# Live cell PNA labelling enables erasable fluorescence imaging of membrane proteins

**DOI:** 10.1101/2020.09.09.286310

**Authors:** Georgina C. Gavins, Katharina Gröger, Michael D. Bartoschek, Philipp Wolf, Annette G. Beck-Sickinger, Sebastian Bultmann, Oliver Seitz

## Abstract

DNA nanotechnology is an emerging field, which promises fascinating opportunities for the manipulation and imaging of proteins on a cell surface. The key to progress in the area is the ability to create the nucleic acid-protein junction in the context of living cells. Here we report a covalent labelling reaction, which installs a biostable peptide nucleic acid (PNA) tag. The reaction proceeds within minutes and is specific for proteins carrying a 2 kDa coiled coil peptide tag. Once installed the PNA label serves as a generic landing platform that enables the recruitment of fluorescent dyes via nucleic acid hybridization. We demonstrate the versatility of this approach by recruiting different fluorophores, assembling multiple fluorophores for increased brightness, and achieving reversible labelling by way of toehold mediated strand displacement. Additionally, we show that labelling can be carried out using two different coiled coil systems, with EGFR and ETBR, on both HEK293 and CHO cells. Finally, we apply the method to monitor internalization of EGFR on CHO cells.

## Main Text

Methods for the selective labelling of cell surface receptors on live cells are needed for investigations of receptor internalization, trafficking and degradation in their native environment. Such information is needed to understand cell signalling from key drug targets such as G-protein coupled receptors (GPCRs) and receptor tyrosine kinases (RTKs), amongst many others. The fusion of fluorescent proteins with the receptor protein of interest allows the visualization via fluorescence microscopy.-^1,2^ Protein tags that are based on self-modifying enzymes such as SNAP,^3^ CLIP^4^ and Halo^5^ proteins or peptide-based tags^6^ extended the scope of protein functional studies to the world of organic fluorescent dyes and techniques beyond fluorescent labelling.

We envisioned that tagging of cell surface proteins with nucleic acid molecules would open up new opportunities for the analysis and, potentially, manipulation of receptors. Interfacing the kingdom of proteins with the area of DNA nanotechnology^7^ offers intriguing applications ranging from advanced imaging methods^8-10^ to the creation of dynamic switches^11-16^ nanomachines ^17^ and logic gates^18-22^. Given the programmability of nucleic acid-nucleic acid interactions, we assumed that labelling with biostable peptide nucleic acid (PNA)^23,24^ tags will provide cell surface proteins with generic landing platforms for the recruitment of fluorescence dyes by means of nucleic acid hybridization (Figure 1a, path A). Moreover, we expected that the use of adaptor strands for multilabelling, and toeholds for strand exchange will provide increased brightness, and allow to reverse labelling, respectively (Figure 1a, path B).

**Figure 1:**
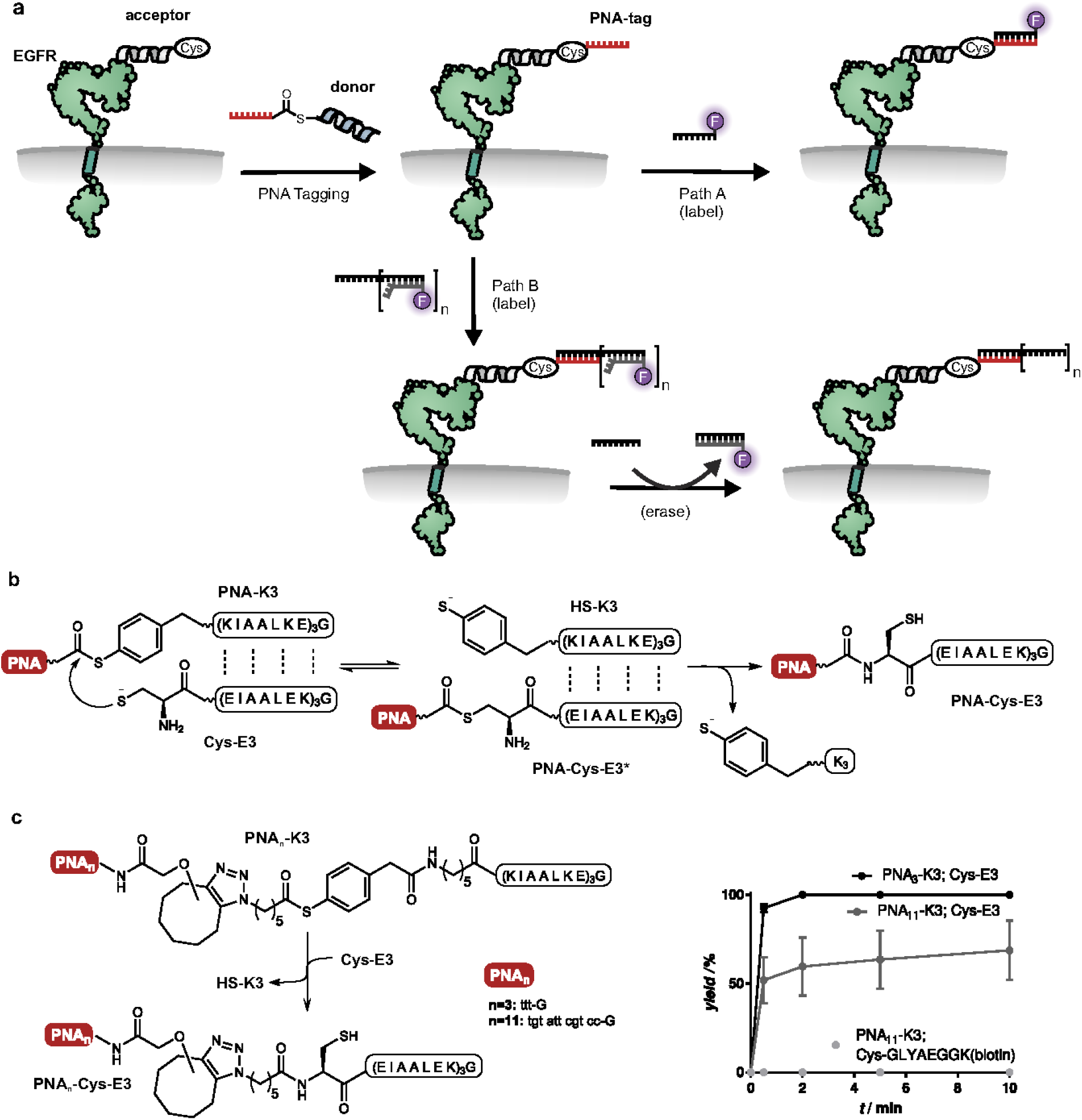
Chemistry of tagging proteins with PNA. **a)** PNA tagging equips cell surface receptors (here: EGFR) with a landing platform that enables recruitment of fluorophores (Path A) or an adaptor strand with multiple DNA strands carrying toeholds for brighter and reversible fluorescent labelling (Path B) by nucleic acid hybridization and toehold-mediated strand exchange. **b)** Principle of PNA tagging by coiled coil induced transfer of PNA from a thioester-linked PNA-coil peptide conjugate (**PNA-K3**) onto a cysteine-coil peptide (**Cys-E3**) linked to a membrane protein of interest. Coiled coil formation triggers a proximity-driven thiol exchange reaction which leads to the thioester-linked PNA-E3 conjugate (**PNA-Cys-E3\***). S→ N acyl transfer yields the stable, amide-linked product **PNA-Cys-E3. c)** Chemical structure of the PNA donor reagents and time course of PNA transfer upon the reaction with the acceptor **Cys-E3** and the control peptide; CGLYAEGGK(biotin). Conditions: 2.5 µM reactants in degassed buffer (100 mM Na_2_HPO_4_, 1 mM TCEP, pH 7.0) at 25°C. Yields were determined by UPLC analysis at 260 nm. Data are presented as mean values +/- SD of n=3 independent experiments.

The most frequently applied procedure used to attach DNA to a protein of interest (POI) on a living cell relies on non-covalent interactions between an antibody-DNA conjugate and the POI, and various chemistries have been developed for linking DNA with antibodies. ^25-30^ However, antibody-based attachment modules introduce a high molecular weight (150 kDa) cargo, which can affect POI function. Metabolic glyco engineering^31,32^ has been used to establish a covalent linkage between DNA and proteins on living cells, but this method does not allow tagging of specific proteins.^33,34^

Live cell covalent labelling of specific proteins with DNA remains a challenge and the two reported methods involve enzymatic fusion proteins. The SNAP-tag methodology^35,36^ introduces a 20 kDa payload in addition to the attached DNA. Another reported method^37^ utilizes a HUH-domain tag to form a phosphotyrosine linkage between itself and ssDNA. Though this method benefits from a smaller, 11.3 kDa fusion protein, for both methods the rapid degradation of oligonucleotides in cell culture remains a problem. Established labelling methods proceeding with smaller expressed peptide tags include FlAsH/ReAsH agents,^38,39^ lipoic acid ligase^40^ or biotin ligase^41^. With these methods, protein tagging typically requires hours and ligase methods call for biorthogonal chemistries to allow secondary labelling of modified conjugates.

Herein we introduce a method that is specific for a 2 kDa large, genetically encoded peptide tag and enables the rapid labelling of membranous proteins with biostable PNA. As opposed to labelling with DNA, PNA provides for improved Watson-Crick base pairing and high stability against degradation by nucleases and proteases. The method is demonstrated on Epidermal Growth Factor Receptor (EGFR) and Endothelial B receptor (ET_B_R) as examples of the RTK and GPCR family of receptors, respectively. Subsequent hybridization with fluorescently labelled PNA or DNA oligomers allows the visualization of receptors on live cells in only a few minutes. We take advantage of DNA nanotechnology techniques to facilitate the analysis of EGF-stimulated EGFR internalization by erasing and fluorescent relabelling of non-internalized EGFRs.

## Results

### PNA is transferred selectively onto peptide coil-tagged EGFR and can be addressed for live cell labelling

The labelling method^42,43^ is based on the interaction between two peptides that form heterodimeric, parallel coiled coils at low nanomolar concentration (Fig. 1b). A notable example are the K3 (KIAALKE)_3_ and the E3 (EIAALEK)_3_ peptides, so called due to their make-up of 3 heptad repeats.^44^ The E3 peptide is equipped with an N-terminal cysteine (**Cys-E3**) and, in analogy to Matsuzakis pioneering work on receptor labelling via coiled coil formation,^45^ serves as the genetically encoded tag for the target protein EGFR. The labelling reagent (**PNA-K3**) is comprised of the K3 peptide segment and a transferable PNA segment conjugated via a thioester linkage. Formation of the coiled coil brings the thioester linkage into proximity of the N-terminal cysteine residue, enabling an acyl transfer reaction to occur at nanomolar concentration of reactants. As a result of the native chemical ligation-type reaction,^46^ the PNA strand will be covalently attached to the Cys-E3-tagged protein (**PNA-Cys-E3**).

For the synthesis of the thioester-linked PNA-K3 conjugate we equipped PNA with an aryl-less cyclooctyne and the K3 peptide with an azidohexanoic acid mercaptoarylthioester and employed strain-promoted azide alkyne cycloaddition^47^ for conjugation (see **PNA**_**n**_**-K3**, Figure 1c; Extended Data Figure 1). To demonstrate the feasibility of the PNA-transfer, we evaluated the reaction by UPLC analysis (Supplementary Figure 5-1). Transfer of the PNA-trimer from conjugate **PNA**_**3**_**-K3** onto the acceptor peptide **Cys-E3** was complete within less than 2 min and succeeded in near quantitative yield despite low (2.5 μM) concentration of reactants (Fig. 1c). The reaction also proceeded smoothly when longer PNA such as the 11-mer in **PNA**_**11**_**-K3** was transferred (Extended Data Figure 2a). As known from native chemical peptide ligation in the absence of thiol additives, attack of the acylating agent at both the amino and the mercapto groups lead to a double acylated product, that is a peptide carrying two PNA strands (Extended Data Figure 2b).^48,49^If required, the S-linked acyl chain can be rapidly removed by treatment with mercaptoethanesulfonate (MESNA) or a low concentration of Cys-K3.9 (Supplementary Figure 5-3). For labelling purposes, this was not deemed necessary. Importantly, no transfer was observed when the PNA donor **PNA**_**11**_**-K3** was allowed to react with a random cysteine peptide (CGLYAEGGK(biotin)) which lacks affinity for the K3 coil peptide (Supplementary Figure 5-2). We inferred that the PNA-labelling reaction will be specific for Cys-E3-tagged proteins.

**Figure 2:**
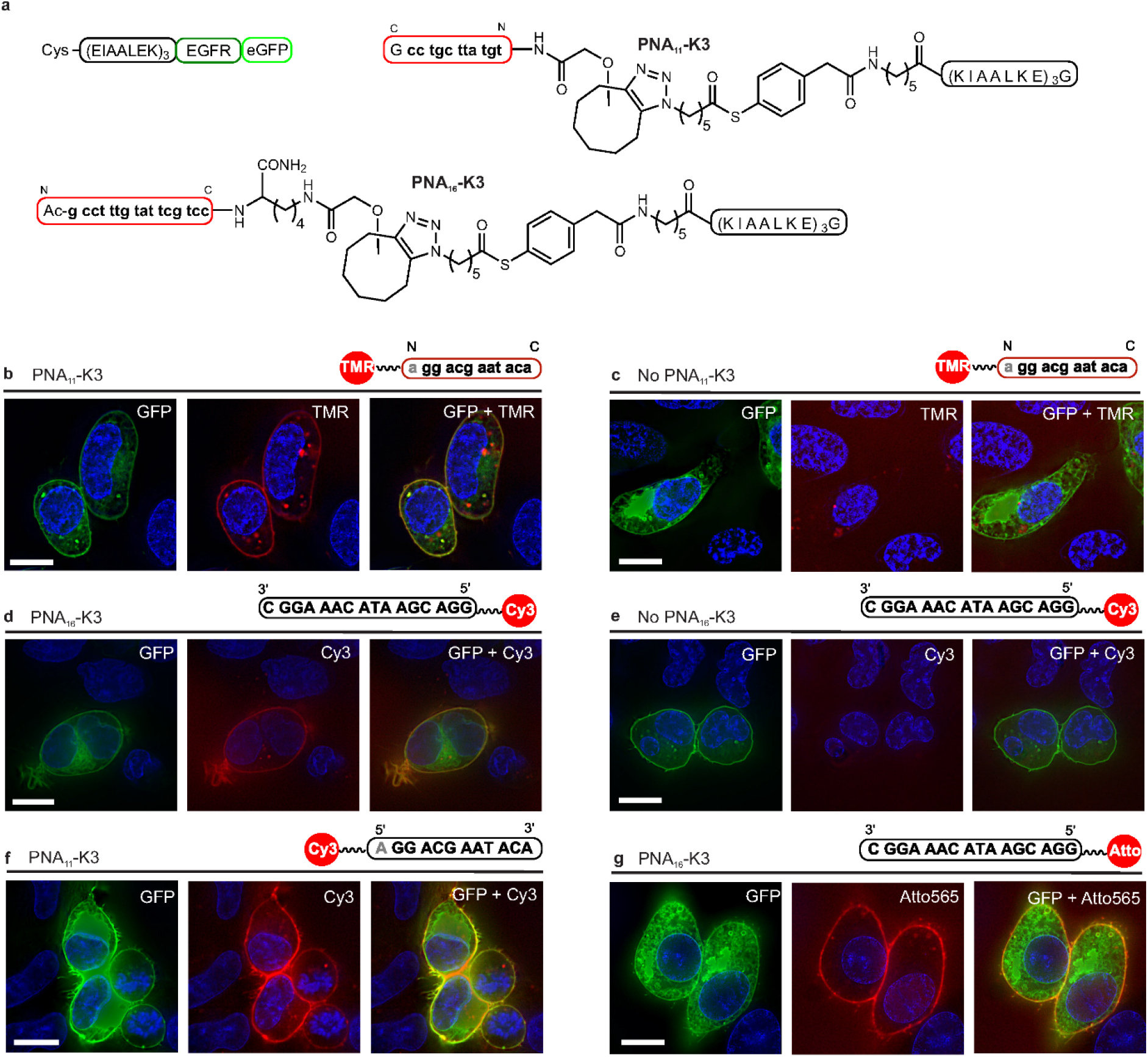
Fluorescence microscopy characterization of PNA-tag enabled fluorescent labelling of Cys-E3-EGFR-eGFP on live HEK293 cells. **a)** Schematic structures of the Cys-E3-EGFR-eGFP construct and the PNA-tagging reagents **PNA**_**11**_**-K3** and **PNA**_**16**_**-K3** used for labelling. **b-f)** After staining of nuclei with Hoechst 33342 (shown in blue), transiently transfected cells were treated with b), f) **PNA**_**11**_**-K3** or d), g) with **PNA**_**16**_**-K3**. For control, PNA tagging was omitted in c) and e). Subsequently, cells were incubated with b), c) TMR-labelled PNA-12mer, d), e) Cy3-labeled DNA-16mer, f) Cy3-labeled DNA-12mer or g) Atto565 labelled 16-mer DNA single strands. Conditions for PNA transfer: 1) 0.1 mM TCEP in PBS (pH 7.0), 2 min, 25°C; 2) 100 nM donor **PNA**_**n**_**-K3** in PBS buffer (pH 7.0), 4 min, 25°C. Conditions for hybridization: 200 nM TMR- / Cy3- / Atto565-labeled PNA/DNA strands in PBS (pH 7.0), 5 min, 25°C. Scale bar = 10 μm. All experiments were repeated independently with similar results 3 times.

To demonstrate the options provided by PNA-tagging we explored the live cell labelling of the EGFR. This receptor tyrosine kinase is an important player in cancer development as its activation can induce proliferation, differentiation, as well as migration and its overexpression is connected to several cancer types such as mammary carcinoma, glioblastoma, non-small cell lung cancer (NSCLC), colorectal, head and neck cancer.^50^Binding of ligands such as EGF triggers EGFR dimerization and signalling via auto-phosphorylation, which eventually leads to endocytosis of the receptor.^51^

HEK293 cells were transiently transfected with an EGF receptor construct comprised of a C-terminal eGFP and an N-terminal Cys-E3-peptide. Fluorescence microscopy of eGFP demonstrated that Cys-E3-EGFR-eGFP correctly localized to the cell membrane (Fig. 2b-g) and intracellular regions where EGFR is biosynthesized and transported. We validated the accessibility of the extracellular Cys-E3 peptide by covalently modifying Cys-E3-EGFR-eGFP with rhodamine (TMR) using a thioester-linked rhodamine-K3 conjugate (TMR-K3, Supplementary Figure 9-1).^37^ The TMR label could not be removed by basic washing, which proved the covalent nature of labelling. We commenced the PNA-tagging studies by incubating Cys-E3-EGFR-eGFP transfected HEK293 cells with 100 nM **PNA**_**11**_**-K3** for 4 min. Additionally, we extended the length of the transferred PNA to a 16mer (**PNA**_**16**_**-K3**) in order to increase the stability of the duplexes formed after nucleic acid hybridization. After washing, we probed the PNA-strands with complementary, fluorescently labelled PNA or DNA (Fig. 2b-g). Both types of oligonucleotides were suitable and cyanine (Cy3), TMR as well as Atto (Atto565) dyes provided sufficient contrast for fluorescence microscopic imaging. While PNA-11mers tags provide sufficient affinity for staining with complementary PNA (Fig. 2b), we found that PNA-16mers are better suited to allow stable interactions with complementary DNA at 37°C (Extended Data Figure 3). We concluded that once a PNA sequence of sufficient length is installed, the colour, brightness and photostability of the label can be easily varied. Staining was restricted to cells showing signals in the GFP channel. Non-transfected cells lack the Cys-E3-EGFR-eGFP construct and therefore serve as internal control demonstrating the selectivity of PNA-transfer for E3-tagged proteins. Staining did not succeed for cells excluded from the PNA tagging step pointing to the specificity of hybridization reactions (Fig. 2c, e). As expected, fluorescence staining by nucleic acid hybridization was limited to the cell surface. However, hydrophobic dye-labelled PNA probes sometimes have low solubility. This can cause problems. For example, the TMR-PNA conjugate showed blob-like structures in control measurements (Fig. 2c). This and the lack of colocalization of the blobs with the features revealed by the eGFP signal points to aggregation. In line with this interpretation, the blobs were not observed in control experiments with the highly hydrophilic DNA conjugates (Fig. 2e).

**Figure 3:**
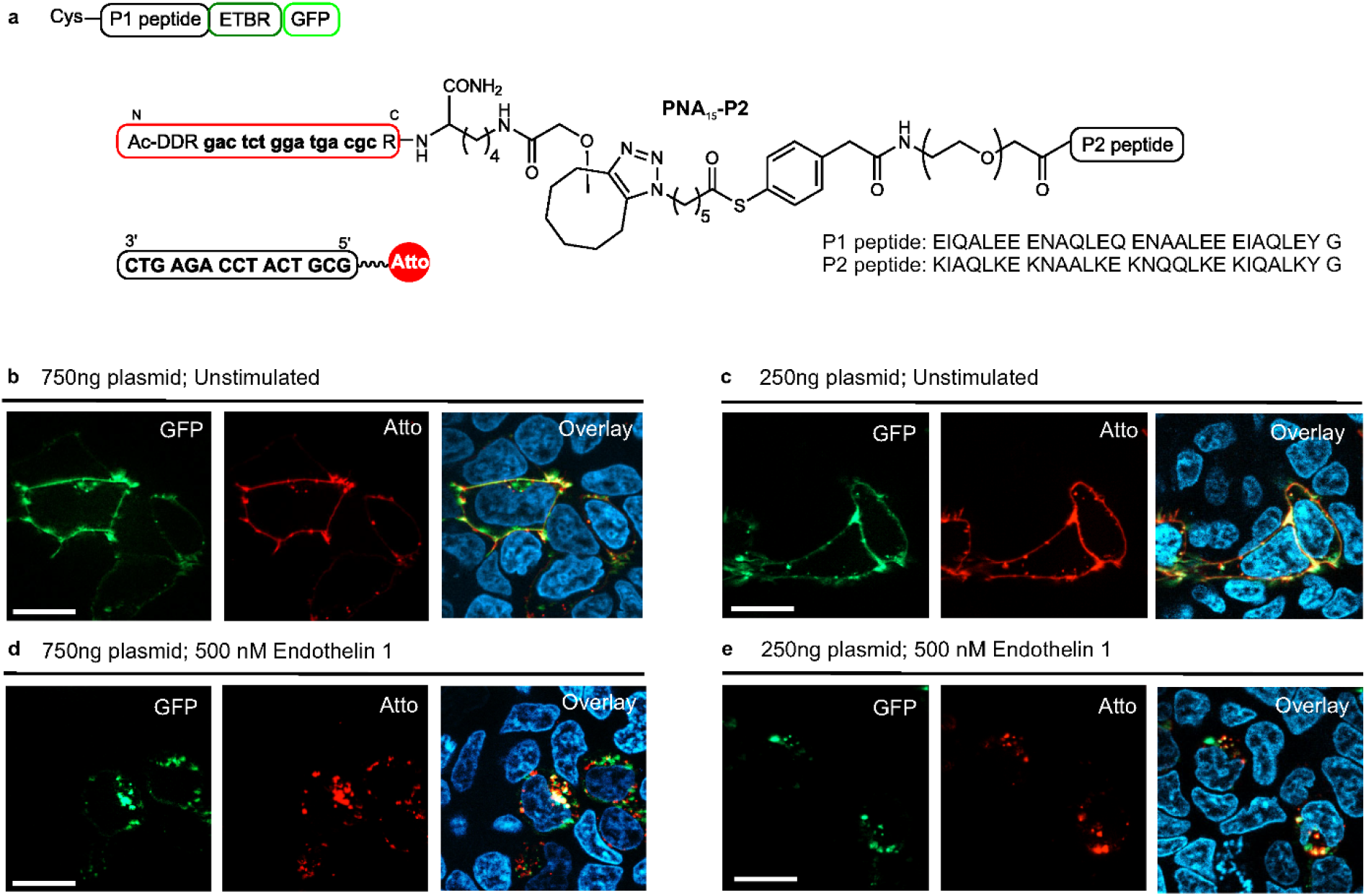
PNA-tag enabled fluorescent labelling of Cys-P1-ET_B_R-GFPspark on HEK293 cells at different transient expression levels. **a)** Schematic structures of the ET_B_R construct, the PNA-tagging reagent **PNA**_**15**_**-P2** and the Atto565 DNA-15mer used for labelling **b-e)** Labelling of ET_B_R transiently expressing Cys-P1-ET_B_R-GFPspark using either b), d) 750 ng or c), e) 250 ng plasmid for transfection. After nuclear staining with Hoechst3342 (shown in blue), PNA-tagging and subsequent hybridization with **Atto565-DNA**_**15**_, cells were b), c) imaged immediately or d), e) stimulated with Endothelin 1 (500 nM) for 60 min at 37°C before imaging. PNA transfer: 100 nM donor **PNA**_**15**_**-P2** in HBSS buffer, 4 min, 37 °C. Hybridization: 200 nM **Atto565-DNA**_**15**_ in HBSS, 4 min, 25°C. Scale bar = 20 μm. Excitation times b) and d): GFPspark 400 ms, Atto565 500 ms. c) GFPspark 340 ms, Atto565 370 ms. e) GFPspark 100 ms, Atto565 170 ms. All experiments were repeated independently with similar results 3 times.

### PNA tagging of ET_B_R at different levels of protein expression

To demonstrate the general applicability of labelling membranous proteins with coiled coil peptide tags, we carried out PNA labelling of transiently expressed Endothelin receptor type B (ET_B_R) on HEK293 cells, using a different coiled coil system. ET_B_R is a GPCR, which mediates vasodilatation and vasoconstriction of endothelial cells for blood pressure regulation. In its ground state the receptor localizes around caveolae on the plasma membrane and upon stimulation with endothelin 1 is extremely rapidly internalized and degraded.^52,53^

Peptides P1 and P2, similarly to E3 and K3, form a heterodimeric parallel coiled coil (Fig. 3a).^54^ The 28mer peptides are composed of four rather than three heptads, resulting in a more stable coiled coil interaction than that of E3/K3.^54^ **PNA**_**15**_**-P2** was synthesized analogously to **PNA**_**16**_**-K3**, with additional charged amino acid residues (Arg and Asp) introduced to the PNA to reduce hydrophobicity and minimize potential non-specific interactions. Transfer reactions in test tubes confirmed similarly rapid PNA transfer at low concentration (200 nM Cys-P1; 500 nM **PNA**_**15**_**-P2**, Supplementary Figure 5-4). Control experiments showed that introducing the Cys-P1-tag behind the endogenous signal peptide of ET_B_R does not alter the G protein-mediated signalling of ET-1 activated ET_B_R (Supplementary Figure 11-1).

HEK293 cells transiently transfected with either 250 or 750 ng of Cys-P1-ET_B_R-GFPspark plasmid were serum starved 30 min prior to PNA tagging with 100 nM **PNA**_**15**_**-P2** for 4 minutes. Hybridization of **Atto565-DNA**_**15**_ followed by fluorescence microscopy confirmed that the staining colocalized with the expression of Cys-P1-ET_B_R-GFPspark on the membrane (Fig 3b, c). Gratifyingly, both expression levels afforded well resolved labelling in the Atto565 channel. In particular, both GFPspark and Atto565 signals exposed clusters of ET_B_R. Stimulation with ET-1 for one hour resulted in complete internalization of the ET_B_R and partial loss of colocalization, which probably is due to the reported rapid degradation of the ET_B_R fusion protein after internalization (Fig 3d, e).^53^

### PNA tagging of EGFR in stable CHO cells accomplishes enhanced brightness by recruiting multiple fluorophore labelled DNA strands

Aiming to demonstrate the advantages of using oligonucleotide-type tagging, we carried out labelling experiments with adaptor DNA strands carrying multiple DNA-fluorophore conjugates. We first generated a stable cell CHO line expressing doxycycline inducible Cys-P1-EGFR-eYFP (Supplementary Figure 8-1). The stable CHO cells were induced with doxycycline 24 hours prior to PNA-tagging with **PNA**_**15**_**-P2** and subsequent staining by nucleic acid hybridization. Fluorescence microscope analysis confirmed that, also in this case, the labelling reaction was specific for Cys-P1-EGFR-eYFP (Supplementary Figures 9-4, 9-5). An immunofluorescence assay measuring EGFR phosphorylation at Y1068 upon EGF stimulation confirmed that the PNA labelling step and DNA hybridization does not disturb EGFR function (Supplementary Figure 10-1).

The use of adaptor strands allows the PNA tag to recruit more than a single fluorescently labelled nucleic acid strand, which allows for enhancements of signal brightness. After PNA tagging with **PNA**_**15**_**-P2**, CHO cells were incubated with a DNA adaptor strand harbouring a 15 nt long stretch for hybridization with the PNA tag, a 3mer gap and either additional 15 nt (**Complex I**) or 87 nt **(Complex II**) long segment for annealing of one or five Cy7-labeled reporter strands (Fig. 4a). Line intensity profiles of Cy7 and eYFP signals revealed that the Cy7-DNA conjugates stain membranous EGFR, while the eYFP signal also highlighted EGFR in intracellular regions (Fig. 4b). The high intensity peaks at membrane regions attest to the increased brightness provided by recruitment of multiple labels with **Complex II** (see Extended Data Fig. 4 and Supplementary Tables 9-1 and 9-2 for quantitative analyses of signal to noise ratios at membrane regions). We performed flow cytometry measurements to analyse the brightness increase upon recruitment of multiple hybridization probes (Supplementary Figure 12-1). In these measurements, CHO cells were treated with mixtures of the 105 nt long adaptor and varied amounts of the Atto647N-labeled oligonucleotides. The ratio of mean fluorescence intensities in the Atto647N and eYFP channels averaged over three independent measurements scaled linearly with the number of Atto647N-DNA equivalents (Figure 4c). This showcases the programmability and robustness of DNA-instructed self-assembly. An extension of this approach to longer adaptor strands or iterative hybridization should enable further brightness enhancements. We speculate that, in analogy to single molecule fluorescence *in situ* hybridization developed for counting of single mRNA molecules,^55^ recruitment of 48 dye-labelled oligonucleotides should be sufficient to achieve single protein sensitivity. A potentially critical issue of labelling by nucleic acid hybridization is stability over time. As opposed to PNA and PNA-DNA complexes, duplex DNA can be subject to degradation by nucleases. The fluorescence microscopic analyses of induced cells starved, after PNA tagging and staining with Atto565-DNA, for varied time in absence of doxycycline indicated that only 22 % of the Atto565 mean intensity was lost within 4 hours, which is comparable to the 16 % loss of the YFP signal (Supplementary Figure 9-10). We note that the use of labelled PNA rather than DNA reporter strands would provide for metabolic stability if label stability was limiting for experiments over longer time courses. However, this was not investigated further since the marginal signal loss over 4 h obtained with the DNA reporter strands does not pose a problem to receptor internalization experiments.

**Figure 4.**
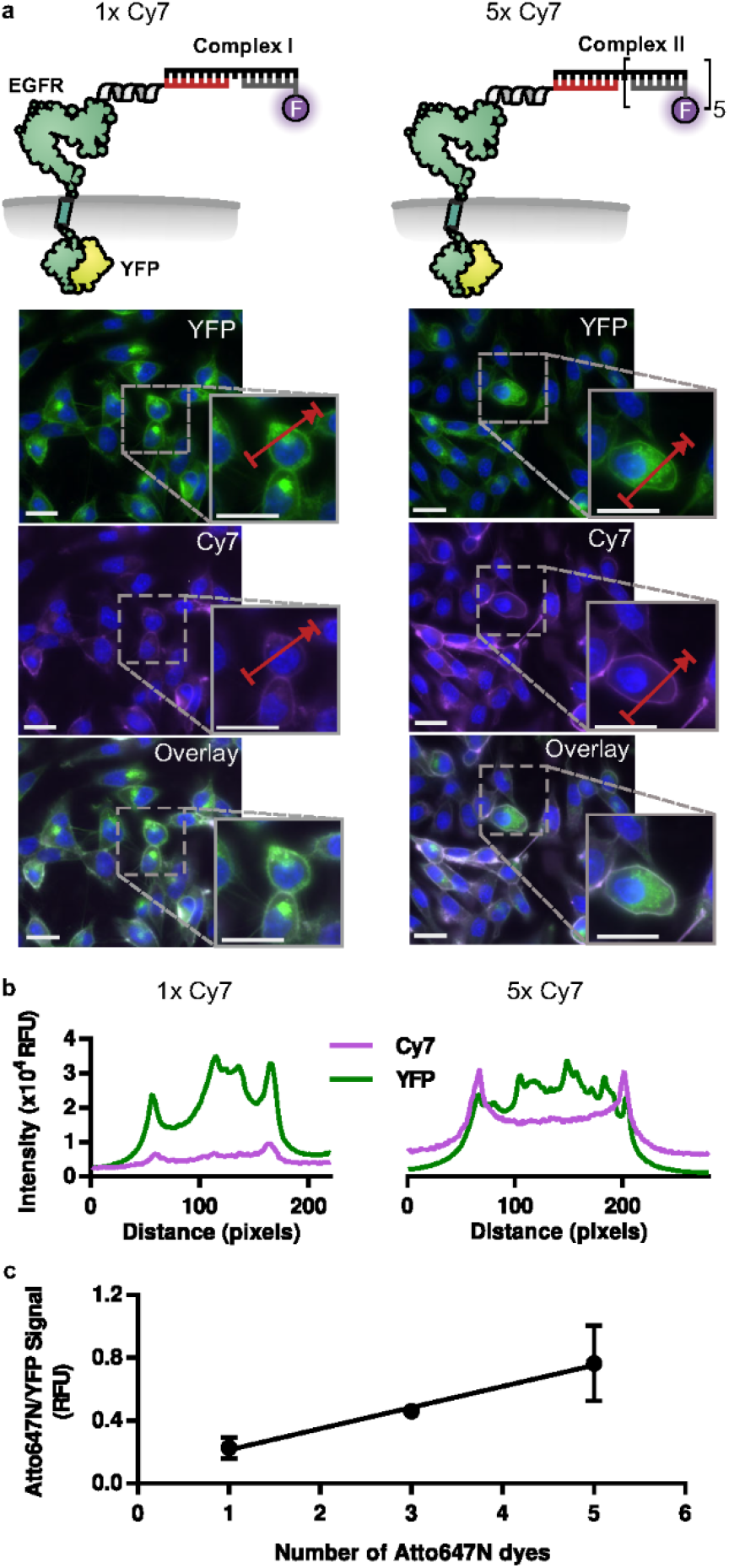
PNA-tag enabled multi fluorophore labelling of Cys-P1-EGFR-eYFP on CHO cells. **a)** Labelling of CHO cells stably expressing Cys-P1-EGFR-eYFP by PNA-tagging and subsequent hybridization with a single Cy7-DNA (**Complex I**, adaptor DNA 33mer with one Cy7-15mer, left) or five Cy7-DNA strands (**Complex II**, adaptor DNA 105mer with five Cy7-15mers, right). Conditions: PNA transfer: 100 nM donor **PNA**_**15**_**-P2** in HBSS buffer, 4 min, 25 °C. Hybridization: 50 nM DNA complex in HBSS-BB (HBSS-Blocking Buffer), 4 min, 25°C. For equally edited pictures see Supplementary Fig. 9-6. Brightness of images depicting 1 × Cy7 labelling was digitally increased relative to 5 × Cy7 for clarity. Scale bar= 20 μm. Experiments were repeated 3 times independently with similar results. **b)** Line intensity profiles for arrowed lines depicted in a). **c)** Dependence of staining intensity on number of Atto647N dyes in complexes used for staining (DNA-105mer + 1 eq or 3 eq or 5 eq Atto647N-DNA-15mer; Complexes IV, V and VI respectively). After labelling as described in a) cells were detached, fixed, and analysed by flow cytometry to determine the ratio of mean Atto647N and YFP fluorescence intensity values, after gating was applied (Supplementary Fig. 12-1). Data is presented as the mean +/- SD of n=3 independent experiments.

**Figure 5:**
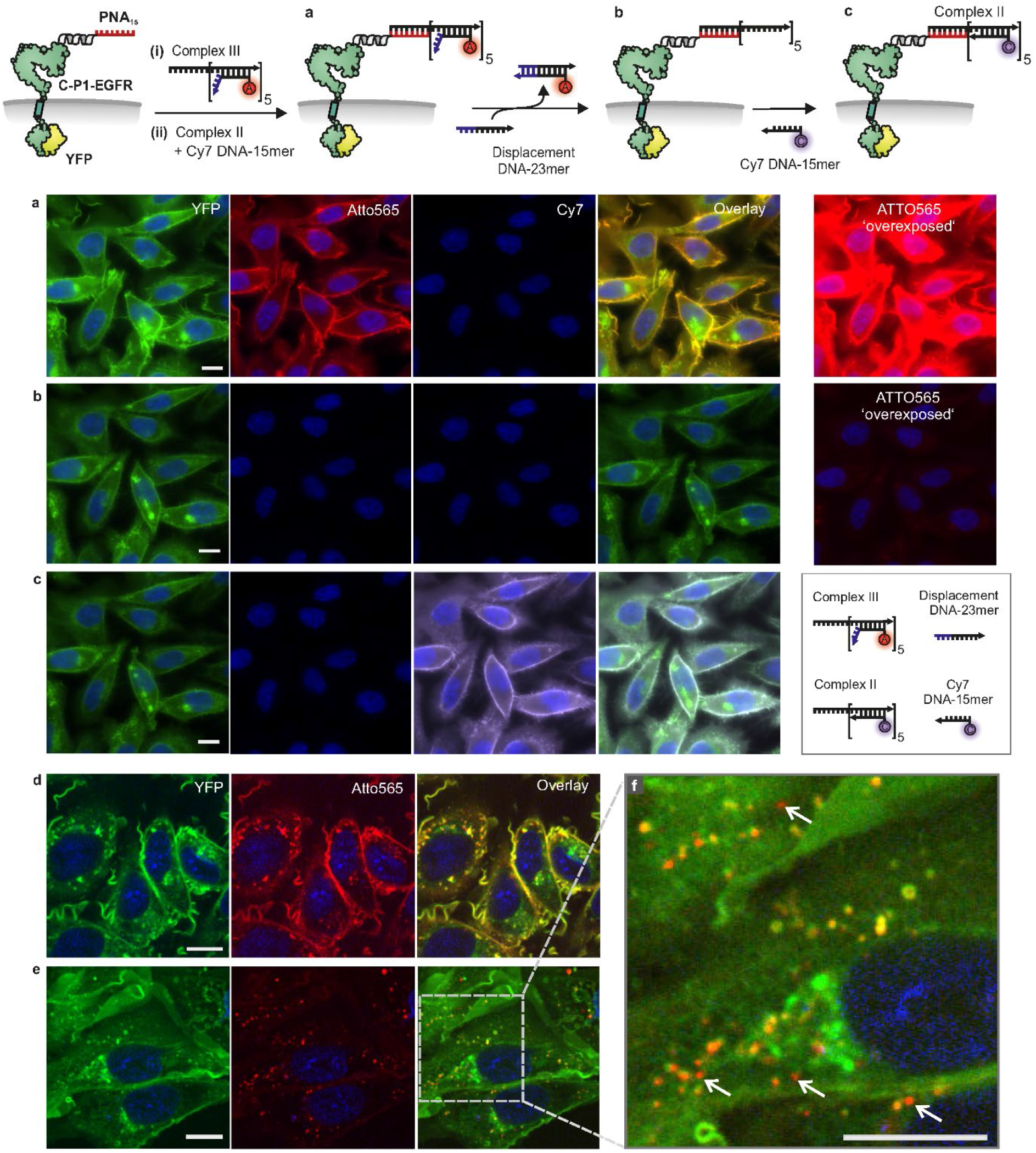
Reversible fluorescent labelling of PNA-tagged Cys-P1-EGFR-eYFP on serum starved CHO cells and visualisation of EGF induced EGFR internalisation. **a-c)** Widefield images of reversible labelling without EGF stimulation. **a)** After staining of nuclei with Hoechst 33342 in HBSS-BB, serum starved Cys-P1-EGFR-eYFP cells were treated with **PNA**_**15**_**-P2** in HBSS for 4 min. Cells were incubated with 50 nM **Complex III** (adaptor DNA-105mer with five Atto565-DNA-23mers containing a toehold for strand displacement) in HBSS-BB for 4 min. Subsequently, 50 nM **Complex II** (adaptor DNA-105mer carrying five Cy7 DNA15mer strands) and 50 nM Cy7 DNA-15mer were added for 4 min. **b)** Toehold mediated strand displacement of the ATTO565-23mer DNA was carried out with 300 nM displacement DNA-23mer for 2 × 5 min in HBSS at 30°C. Brightness of Atto565 images in row a) and b) were digitally increased to give ‘overexposed’ images which illustrate that Atto565 labelling was completely removed in b). **c)** 100 nM Cy7 DNA-15mer was hybridized for 3 min to illustrate restaining. **d-f)** Spinning disc confocal fluorescence microscopy with EGF stimulation **d)** CHO cells labelled with **Complex III** as described in a) were stimulated with EGF (100 nM) for 15 min followed by **e)** toehold mediated strand displacement in the presence of 100 nM EGF for 7 min in HBSS at 30°C. **f)** Zoomed in overlay picture from e). Arrows highlight newly internalized vesicles, which are difficult to identify in the YFP channel. Atto565 brightness is increased f) relative to d) and e). Zoomed out images available in Extended Data Fig. 5. Scale bar= 10 μm. All experiments were repeated 3 times independently with similar results.

### Fluorescent labelling of EGFR is erasable and exchangeable using toehold strand displacement

We next applied the toehold mediated strand displacement^56,57^ to our system to enable the erasure and exchange of fluorescent labelling. After **PNA**_**15**_**-P2** tagging, CHO cells stably expressing Cys-P1-EGFR-eYFP were incubated with 50 nM **Complex III** (Figure 5a); an adaptor DNA-105mer carrying five Atto565 labelled DNA-23mers. Each DNA-23mer bears a 15 nt region complementary to the adaptor strand and an 8 nt toehold region, which seeds the hybridization with an invading ‘displacement DNA-23mer’ strand. To confirm that all the PNA sites were saturated with **Complex III**, and that in turn **Complex III** was saturated with Atto565-DNA-23mer, cells were subsequently incubated with both 50 nM **Complex II** and Cy7 DNA-15mer. No Cy7 signal was observed, confirming that all PNA sites on EGFR were saturated with **Complex III** (Figure 5a). Next, displacement DNA-23mer; that is the reverse complement of the Atto565-DNA-23mer; was added to induce the complete removal of the Atto565-DNA (Figure 5b). Subsequent incubation with Cy7-DNA-15mer for 3 min resulted in relabelling of the adaptor DNA-105mer hybridized to **PNA**_**15**_-tagged EGFR (Figure 5c).

We envisioned that erasable fluorescent labelling provides the opportunity to facilitate the study of membrane protein trafficking by excluding signals from non-internalized proteins. In an exemplary experiment, the CHO cells stably expressing Cys-P1-EGFR-eYFP were starved prior to PNA tagging. Hybridization of **Complex III** with PNA-tagged EGFR allowed visualization of membranous EGFR in the Atto565 channel (Extended Data Figure 5a). Next, cells were stimulated with 100 nM EGF. After 15 min stimulation, internalization of Atto565 labelled EGFR was observed (Fig. 5d, Extended Data Fig.5b). However, under these conditions signals from non-internalized EGFR are as intense, if not brighter, than signals from internalized EGFR. This issue is most pronounced in the eYFP channel, which also reveals signals from freshly transcribed EGFR molecules and EGFRs on their way to the membrane. Treatment with displacement DNA-23mer removed the Atto565-DNA label from remaining membrane-standing EGFR (Fig. 5e, Extended Data Fig. 5c). Only EGFR which was internalized during the defined time frame was observed in the Atto565 channel. This facilitates the identification of transport vesicles that are close to membrane features (Figure 5f, for separate channels of zoomed image see Supplementary Figure 9-9). Additionally, internalized EGFR was visualized in 3D by acquiring Atto565 z-stacks (Supplementary Figure 9-8). The de-labelled EGFR molecules that remained at the membrane despite EGF stimulation were relabelled with 100 nM Atto647N-DNA15mer (Extended Data Fig. 5d), which highlights the non-quantitative nature of EGFR internalization under these conditions. The same experiment was repeated using widefield microscopy analysis as described and also without addition of **PNA**_**15**_**-P2** or DNA (Supplementary Figure 9-7). The YFP images for both experiments give little information of the fate of EGF stimulated, internalized EGFR. Often, multicolour pulse-chase labelling experiments are performed for tracking of membrane receptors.^48^ The ability to erase the label from non-internalized receptor adds extra temporal control to these experiments.

## Discussion

In a growing number of publications, nucleic acid technology is applied to proteins in/on fixed cells; mainly to improve protein detection by barcoding, super-resolution microscopy or DNA-based signal amplification.^7^ A remarkable report described the use of SNAP tag methodology to create truncated DNA-protein hybrid receptors on Jurkat cells.^36^ Recently, metabolic glycoengineering was used to label cell surface carbohydrates with DNA.^33,34^ Lovendahl *et al* presented HUH-endonuclease domain fusion to covalently link DNA to proteins.^36^ We attribute this rather scarce literature precedence to a lack of robust methods that allow conjugation without detriment to the protein function. The herein developed PNA-tagging method allowed us to harness the assets provided by nucleic acid-instructed assembly to reversibly fluorescently label native-like functional proteins on the surface of live cells. The method relies on a genetically encoded 22 or 29 amino acid long peptide tag.^42,43,54,58^ This attachment module has only 10% of the size of the SNAP tag, and around 20% of the HUH domain tag previously used for live cell nucleic acid-protein conjugation. The Cys-E3 or Cys-P1 tags are fused to the N-terminus of the protein of interest, which typically is exposed to the cell exterior for the majority of cell surface receptor proteins. We showed that labelling with PNA succeeded within minutes. The required PNA labelling agent is readily accessible by a strain-promoted ligation of a cycloalkyne-PNA conjugate with a thioester-linked azido-peptide. The modularity of this approach facilitates diversification of the labelling reagents.

Once installed, the PNA tag is extremely versatile. In contrast to DNA, PNA persists nucleases. We showed fluorescent labelling and imaging of EGFR on live cells by means of hybridization with PNA- or DNA reporter strands containing four different dyes i.e. TMR, Cy3, Atto565 and Cy7. We illustrated how common methods utilized in DNA nanotechnology can be easily employed to this system to offer new functionalities i.e. using toehold mediated strand displacement to carry out erasable labelling, allowing more sophisticated analysis of protein trafficking. We believe that the improvement of brightness observed upon recruitment of multiple reporter strands will prove useful for fluorescence microscopy. Of note, the PNA tag and its assembled DNA structures were stable to internalization. PNA-DNA duplexes are protected from nuclease activity, however over long time scales double stranded DNA is vulnerable to degradation by DNases inside the cell. This could be avoided by the use of biostable DNA mimetics such as PNA, LNA or methylated DNA. PNA and LNA is commercially available and LNA-DNA has been shown to be effective in toehold-mediated strand displacement.^59^

The PNA-tagging method may render advanced imaging technologies based on DNA-PAINT^8,10^ or reiterative hybridization^60^ applicable to live cells. The PNA tag could also be used for receptor preorganization. DNA has been used to control distance restraints^61^ and DNA based chemical dimerisers have been used to preorganise cell surface proteins.^62^ After PNA-tagging and staining by nucleic acid hybridization the protein of interest carries a 12 kDa cargo for systems involving a single reporter strand. Similar structures on SNAP tags or HUH domains would lead to a 30 kDa or 21 kDa cargo. In addition, the label dimensions are fundamentally different. Coiled coils and nucleic acid duplexes have a cylinder-shaped 3D structure, which is less bulky than the more globular structures of enzyme tags. We, therefore, expect that protein receptors can tolerate even large nucleic acid duplex cargo. In this regard, it is encouraging that receptor internalization tolerated the 65 kDa molecular mass increase accompanying staining with adaptor and five reporter strands.

We illustrated that the method can be applied with two different coiled coil systems and showcased the versatility by labelling two distinct target proteins. We believe the programmability of coiled coils brings an extra level of control. The *de novo* design of coiled coils has brought about a number of new orthogonal coiled coils^54,63^ which would allow orthogonal labelling. This will be addressed in further work.

In conclusion, the presented PNA tagging approach provides a method to equip proteins on live cells with a generically addressable, biostable control unit. The PNA tag provides an unlimited choice over the kind and, potentially, the number of fluorescence dyes used for labelling. Moreover, by covalent attachment of an oligonucleotide and subsequent nucleic acid self-assembly, we imagine several other application scenarios. While herein we focused on fluorescent labelling, in ongoing work we will explore orthogonal PNA labelling using orthogonal coiled coils, and preorganization of membrane proteins by way of nucleic acid assembly. PNA tagging may prove useful for a broad range of applications in cell biology

## Supporting information

Extended Data Fiigures

## Acknowledgements

We acknowledge financial support from the Deutsche Forschungsgemeinschaft (SPP 1623 and SFB 765). M.D.B is a fellow of the International Max Planck Research School for Molecular Life Sciences (IMPRS-LS). P.W. is a member of the Graduate School Leipzig School of Natural Sciences – Building with Molecules and Nano-objects. We thank Jonathan Lotze (University Leipzig), Stephan Korte and Andreas Herrmann (Humboldt University Berlin) for help with Confocal Laser Scanning Microscopy and Knut Rurack (Bundesanstalt für Materialforschung und Prüfung, Berlin) for providing access to flow cytometry.

## Author Contributions

G.C.G., K.G. and P.W. performed the experiments. G.C.G, K.G. and O.S. designed the experiments and analysed the data. M.D.B. and S.B. constructed the stable CHO cell lines. O.S. conceived the experiments. P.W. and A.G.B-S. designed experiments for labelling of ET_B_R. All authors discussed the results and contributed to the preparation and editing of the manuscript.

## Corresponding authors

Correspondence to Oliver Seitz

## Competing Interests Statement

The authors declare no competing interests.

## Methods

### PNA and DNA sequences (5’->3’ or N->C)

**PNA**_**3**_: ttt; **PNA**_**11**_: tgt att cgt cc; **PNA**_**16**_: gcc ttt gta ttc gtc c; **Atto565/Cy3 DNA-16mer**: GGA CGA ATA CAA AGG C; **Cy3 DNA-12mer**: A GGA CGA AAT ACA; **TMR PNA-12mer**: a gg acg aat aca Gly; **PNA**_**15**_: gac tct gga tga cgc; **Atto565 DNA-15mer**: GCG TCA TCC AGA GTC; **Cy7/Atto647DNA-15mer**: GAC ACC ACT TAC CAG; **DNA-33mer**: GCG TCA TCC AGA GTC CTA CTG GTA AGT GGT GTC; **DNA-105mer**: GCG TCA TCC AGA GTC CTA CTG GTA AGT GGT GTC CTA CTG GTA AGT GGT GTC CTA CTG GTA AGT GGT GTC CTA CTG GTA AGT GGT GTC CTA CTG GTA AGT GGT GTC; **Atto565 DNA-23mer**: GAC ACC ACT TAC CAG ATA GCA CA; **Displacement DNA-23mer**: TGT GCT ATC TGG TAA GTG GTG TC; **ComplexI**: DNA-33mer + Cy7-15mer; **ComplexII**: DNA-105mer + 5(Cy7-15mer); **ComplexIII**: DNA105mer +5(Atto565-DNA23mer) **Complex IV**: adaptor DNA-105mer + Atto647N-15mer, **Complex V:** adaptor DNA-105mer 3(Atto647N-15mer) and **Complex VI:** adaptor DNA-105mer +5(Atto647N-15mer).

DNA complexes were hybridised by premixing to a final complex concentration of 10 μM in HBSS, heating to 70°C and cooling to RT over 30 min.

### Synthesis of thioester-linked peptide-PNA conjugates (PNAn-K3)

The synthesis was performed in three steps (Extended Data Figure 1). Step 1: The K3/P2 peptide was assembled on glycine-loaded TGR resin according to the Fmoc strategy. S-Mmt-protected mercaptophenylacetic acid (MPAA) was coupled to the N-terminal end. After treatment with CH_2_Cl_2_:TFA:TIS (96:2:2) for 2 ×; 1 min, azidohexanoic acid was coupled to the liberated mercaptane by using HCTU as activating reagent in presence of N-methylmorpholine. The thioester-linked azido-peptide was obtained after treatment with TFA:TIS:H_2_O (96:2:2), ether precipitation and HPLC purification. Step 2: A PNA oligomer containing an unprotected N-terminus or an unprotected lysine side chain was prepared by Fmoc-based solid phase synthesis. The crude product obtained after TFA cleavage and ether precipitation was dissolved in DMF. For coupling of aryl-less cycloctyne (ALO), a solution of 2-(cyclooct-2-ine-1-yloxy)acetic acid (20 eq) and DIC (20 eq) in DMF was agitated for 10 min and added to the PNA solution (approx. 10 mM final concentration). After 1 h the mixture was purified by RP-HPLC. Step 3: The purified thioester-linked azido-peptide and the ALO-PNA conjugate were dissolved in a 3:1 stoichiometry in MeCN: H_2_O (3:1) with 0.1% TFA and allowed to react overnight at 0.5 – 1 mM concentration. Purification was performed by RP-HPLC in gradients of MeCN:H_2_O (0.1% TFA).

### PNA tagging and fluorescence microscopic imaging of EGFR with hybridization probes on transiently transfected HEK293 cells

Prior to cell seeding 8-well μ-slides (ibidi, ibiTreat) were coated with 0.01% poly-D-lysine. After 10 min incubation the solution was removed and the slides were allowed to dry. HEK293 cells (10,000) were seeded and incubated in DMEM (10% FBS) over night at 37°C. For transfection, 100 ng vector and 1 μL Lipofectamine®2000 in 200 μL Opti-MEM® was added to each chamber. After 1 h the DMEM (10% FBS; 100 μL) was added and the cells were incubated over night at 37°C. Cells were stained with Hoechst 33342 and treated for 2 min with 0.1 mM TCEP in PBS and then incubated for 4 min with 100 nM PNA donor such as PNA_16_-K3 in PBS (pH 7.0). Cells were washed with PBS (2 x) prior to addition of 200 nM fluorescent labelled DNA or PNA in PBS. Alternatively, 1 μM connector DNA was added along with 3 μM fluorescent labelled PNA. After 5 min cells were washed with PBS. Fluorescence microscopy was performed by using an IX83 microscope from Olympus. Transfected cells were imaged in three different channels (Hoechst 33342: λ_ex_ = 350 ± 50 nm, λ_em_ = 460 ± 50 nm; YFP: λ_ex_ = 500 ± 24 nm, λ_em_ > 520 nm; TMR/Cy7/Atto565: λ_ex_ = 575 ± 25 n,m λ_em_ > 593 nm) in 0.27 μm z stacks and the images were processed by Wiener deconvolution.

### PNA tagging and fluorescence microscopic imaging of ET_B_R with hybridization probes on transiently transfected HEK293 cells

Prior to cell seeding 8-well μ-slides (ibidi, ibiTreat) were coated with 0.01% poly-D-lysine. After reaching 70 % confluence, HEK293 cells were transiently transfected with the pCMV3-Cys-P1-ET_B_R-GFPspark and Lipofectamine™2000 according to the manufacturer’s protocol. Either 250 ng or 750 ng of plasmid DNA were used for transfection, while keeping the total amount of DNA 1000 ng, balanced with pcDNA3 (empty vector). Labelling experiments and microscopic imaging was performed after 24 h post transfection. For microscopic analysis the supernatant was exchanged with OptiMEM and cell nuclei were stained with 1 µL Hoechst33342 for 20 min. For PNA transfer, 100 nM PNA_15_-P2 peptide in HBSS was added after medium exchange for 4 min at 37 °C. The cell monolayer was washed once with HBSS and the cells were incubated with 200 nM ATTO565-DNA_15_ in HBSS for 4 min at room temperature followed by washing the monolayer twice with HBSS.. For stimulation with ET-1, the peptide was dissolved in OptiMEM at a concentration of 500 nM. Cells were incubated for 1 h at 37 °C with ET-1 before microscopic analysis. Fluorescence images were acquired in HBSS after ET-1 removal by washing with HBSS. Fluorescence images were taken by using a Zeiss Axio Observer.Z1 microscope with an ApoTome.2 Imaging System, a C-Apochromat 63x/1.20 W Corr M27 objective, an AxioCamMRm camera and the ZEN 2.0 software. Transfected cells were imaged using different Zeiss filter sets: Hoechst33342 (λ_ex_:365, beamsplitter: 395; λ_em_:420), GFP (λ_ex_: 470/40, beamsplitter: 495: λ_em_:525/50), Atto565 (λ_ex_: 565/30; beamsplitter: 585; emission: λ_em_ 620/60).

### PNA tagging and fluorescence microscopic imaging with hybridization probes on stable CHO cell lines

Prior to cell seeding 8-well μ-slides (ibidi, ibiTreat) were coated with 0.01% poly-D-lysine. After 10 min incubation the solution was removed and the slides were allowed to dry. CHO cells (20,000) were seeded and incubated in Ham’s F12 medium (10% FBS) overnight at 37°C. Cells were induced by addition of 0.1µg/mL doxycycline. For multilabelling and strand displacement experiments, media was replaced after 5 hours with serum free media (0.1µg/mL doxycycline). Cells were stained with Hoechst 33342 in either HBSS-BB for 10 min and treated 4 min with 100 nM PNA donor PNA_15_-P2 in HBSS-BB. Cells were washed with HBSS-BB prior to addition of 100 nM fluorophore labeled DNA or 50 nM DNA complex HBSS-BB. After 4 min cells were washed with HBSS-BB and imaged in HBSS-BB. Fluorescence microscopy was performed by using an IX83 microscope from Olympus. Cells were imaged in four different channels: Hoechst 33342: λex = 350 ± 50 nm λ_em_ = 460 ± 50 nm; YFP: λ_ex_ = 500 ± 24 nm λ_em_ > 520 nm; Atto565: λ_ex_ = 575 ± 25 nm λ_em_ > 593 nm. Cy7: λ_ex_ = 710 ± 75 nm, λ_em_ 810 ± 90 nm. HBSS-BB= HBSS-Blocking Buffer: 0.1mg/mL salmon sperm DNA, 0.2% BSA, 1x ProLong™ Live Antifade in HBSS (+Mg, Ca).

### EGFR internalization and membrane signal erasure on stable CHO cells

Cells were treated as described above. During measurement cells were maintained at 30°C with 5% CO_2_. After hybridization of 50 nM Complex III (adaptor DNA-105mer with five Atto565-DNA-23mers) in HBSS-BB, cells were washed with HBSS-BB then stimulated with EGF (100 nM) in HBSS-BB at 30°C for 15 minutes. For Figure 5a-c) EGF stimulation was omitted. Displacement DNA-23mer was added to a concentration of 300 nM, for 2 × 5 mins. Cells were washed twice with HBSS-BB before adding 100 nM Cy7-DNA-15mer (Widefield microscopy; Fig. 5 a-c), Supporting Figure 9-6) or 100nM Atto647N-DNA-15mer (Confocal microscopy: Fig 5d-f), Extended Data Figure 5). Cells were washed with HBSS-BB before imaging with an a Visitron VisiScope with an *Olympus* IX83 microscope and a *Yokogawa* CSU-W1 spinning disk unit (confocal microscopy) or a widefield IX83 microscope from *Olympus* (widefield microscopy). Confocal imaging: Diode lasers: Hoechst 33342) 405 nm; YFP) 488 nm; Atto565) 561 nm; Atto647N) 640 nm. Dichroic emission filters Hoechst 33342) λ_em_ = 460 ± 50 nm; YFP) λ_em_ 470 ± 24 nm; Atto565) λ_em_ 600 ± 50 nm. Atto647N) λ_em_ = 700 ± 75 nm. Widefield imaging: Hoechst33342: λex = 350 ± 50 nm λem = 460 ± 50 nm; YFP: λex = 500 ± 24 nm λem > 520 nm; TRITC: λex = 575 ± 25 nm λem > 593 nm Cy7.

### Flow cytometry analysis of PNA tagging and hybridization with multi-Atto647N-DNA constructs on stable CHO cell lines

CHO cells (25,000) stably expressing Cys-P1-EGFR-eYFP were seeded in µ-slides (ibidi, ibiTreat) and incubated overnight in Hams F12 media (10% FBS). Media was exchanged to Ham F12 media (-) FBS with 0.1 µg/mL doxycycline for 15-17 hours prior to experiment. Cell were washed once with Dulbecco’s phosphate-buffered saline (DPBS) then incubated **PNA**_**15**_**-P2** (100nM, 4 min) in HBSS. After washing with HBSS-BB cells were incubated with 100 nM DNA complex for 5 mins in HBSS-BB. The DNA complexes used for hybridization were: **Complex IV** (adaptor DNA-105mer with one Atto647N-15mer), **Complex V** (adaptor DNA-105mer with three Atto647N-15mers) and **Complex VI** (adaptor DNA-105mer with five Atto647N-15mers). Cells were washed with DPBS before addition of Trypsin/EDTA (0.25% trypsin / 0.02% EDTA 30 µL) for five minutes at room temperature. 300 µL Hams F12 media (10% FBS) was added and detached cells centrifuged (300g, 8 min), media removed and the cells resuspended in DPBS (200 µL). Paraformaldehyde (8% in PBS, 200 µL) was added for 10 minutes before centrifugation (300 g 8 min). The pellet was resuspended in DPBS (200 or 300 µL) before performing flow cytometry experiments on a BD Accuri™ C6 (BD Biosciences). Excitation lasers: YFP) 488 nm; Atto647N) 640 nm. Emission filters: YFP) 533/30 nm; Atto647N) 675/25 nm. HBSS-BB= HBSS-Blocking Buffer: 0.1mg/mL salmon sperm DNA, 0.2% BSA,in HBSS (+Mg, Ca).

## Data Availability Statement

The data that support the findings of this study are available from the corresponding author upon reasonable request.

## Notes

### Competing Interest Statement

The authors have declared no competing interest.

## References

1 Rizzuto, R., Brini, M., Pizzo, P., Murgia, M. & Pozzan, T. Chimeric green fluorescent protein as a tool for visualizing subcellular organelles in living cells. Curr. Biol. 5, 635–642, doi: 10.1016/S0960-9822(95)00128-X (1995).

2 Chalfie, M., Tu, Y., Euskirchen, G., Ward, W. W. & Prasher, D. C. Green fluorescent protein as a marker for gene expression. Science 263, 802–805, doi: 10.1126/science.8303295 (1994).

3 Keppler, A. et al. A general method for the covalent labeling of fusion proteins with small molecules in vivo. Nat. Biotechnol. 21, 86–89, doi: 10.1038/nbt765 (2003).

4 Gautier, A. et al. An Engineered Protein Tag for Multiprotein Labeling in Living Cells. Chem. Biol. 15, 128–136, doi: 10.1016/j.chembiol.2008.01.007 (2008).

5 Los, G. V. et al. HaloTag: A novel protein labeling technology for cell imaging and protein analysis. ACS Chem. Biol. 3, 373–382, doi: 10.1021/cb800025k (2008).

6 Lotze, J., Reinhardt, U., Seitz, O. & Beck-Sickinger, A. G. Peptide-tags for site-specific protein labelling in vitro and in vivo. Mol. BioSyst. 12, 1731–1745, doi: 10.1039/c6mb00023a (2016).

7 Seeman, N. C. & Sleiman, H. F. DNA nanotechnology. Nat. Rev. Mater. 3, doi: 10.1038/natrevmats.2017.68 (2017).

8 Schnitzbauer, J., Strauss, M. T., Schlichthaerle, T., Schueder, F. & Jungmann, R. Super-resolution microscopy with DNA-PAINT. Nat. Protoc. 12, 1198–1228, doi: 10.1038/nprot.2017.024 (2017).

9 Duose, D. Y. et al. Configuring robust DNA strand displacement reactions for in situ molecular analyses. Nucleic Acids Res. 40, 3289–3298, doi: 10.1093/nar/gkr1209 (2012).

10 Jungmann, R. et al. Single-molecule kinetics and super-resolution microscopy by fluorescence imaging of transient binding on DNA origami. Nano Lett. 10, 4756–4761, doi: 10.1021/nl103427w (2010).

11 Diezmann, F. & Seitz, O. DNA-guided display of proteins and protein ligands for the interrogation of biology. Chem. Soc. Rev. 40, 5789–5801, doi: 10.1039/c1cs15054e (2011).

12 Peri-Naor, R., Ilani, T., Motiei, L. & Margulies, D. Protein-protein communication and enzyme activation mediated by a synthetic chemical transducer. J. Am. Chem. Soc. 137, 9507–9510, doi: 10.1021/jacs.5b01123 (2015).

13 Schade, M. et al. Remote Control of Lipophilic Nucleic Acids Domain Partitioning by DNA Hybridization and Enzymatic Cleavage. J. Am. Chem. Soc. 134, 20490–20497, doi: 10.1021/ja309256t (2012).

14 Röglin, L., Ahmadian, M. R. & Seitz, O. DNA-controlled reversible switching of peptide conformation and bioactivity. Angew. Chem., Int. Ed. 46, 2704–2707, doi: doi: 10.1002/anie.200603889 (2007).

15 Freeman, R. et al. Instructing cells with programmable peptide DNA hybrids. Nat. Comm. 8, doi: 10.1038/ncomms15982 (2017).

16 Ueki, R., Atsuta, S., Ueki, A. & Sando, S. Nongenetic Reprogramming of the Ligand Specificity of Growth Factor Receptors by Bispecific DNA Aptamers. J. Am. Chem. Soc. 139, 6554–6557, doi: 10.1021/jacs.7b02411 (2017).

17 Leung, K., Chakraborty, K., Saminathan, A. & Krishnan, Y. A DNA nanomachine chemically resolves lysosomes in live cells. Nat. Nanotechnol. 14, 176–183, doi: 10.1038/s41565-018-0318-5 (2019).

18 Janssen, B. M. G., Van Rosmalen, M., Van Beek, L. & Merkx, M. Antibody activation using DNA-based logic gates. Angew. Chem., Int. Ed. 54, 2530–2533, doi: 10.1002/anie.201410779 (2015).

19 Qian, L., Winfree, E. & Bruck, J. Neural network computation with DNA strand displacement cascades. Nature 475, 368–372, doi: 10.1038/nature10262 (2011).

20 Elbaz, J. et al. DNA computing circuits using libraries of DNAzyme subunits. Nat. Nanotechnol. 5, 417–422, doi: 10.1038/nnano.2010.88 (2010).

21 Hemphill, J. & Deiters, A. DNA computation in mammalian cells: MicroRNA logic operations. J. Am. Chem. Soc. 135, 10512–10518, doi: 10.1021/ja404350s (2013).

22 You, M. et al. DNA “nano-claw”: Logic-based autonomous cancer targeting and therapy. J. Am. Chem. Soc. 136, 1256–1259, doi: 10.1021/ja4114903 (2014).

23 Egholm, M. et al. Pna Hybridizes to Complementary Oligonucleotides Obeying the Watson-Crick Hydrogen-Bonding Rules. Nature 365, 566–568, doi: 10.1038/365566a0 (1993).

24 Demidov, V. V. et al. Stability of peptide nucleic acids in human serum and cellular extracts. Biochem. Pharmacol. 48, 1310–1313, doi: 10.1016/0006-2952(94)90171-6 (1994).

25 Söderberg, O. et al. Direct observation of individual endogenous protein complexes in situ by proximity ligation. Nat. Methods 3, 995–1000, doi: 10.1038/nmeth947 (2006).

26 Meyer, R., Giselbrecht, S., Rapp, B. E., Hirtz, M. & Niemeyer, C. M. Advances in DNA-directed immobilization. Curr. Opin. Chem. Biol. 18, 8–15, doi: 10.1016/j.cbpa.2013.10.023 (2014).

27 Trads, J. B., Tørring, T. & Gothelf, K. V. Site-Selective Conjugation of Native Proteins with DNA. Acc. Chem. Res. 50, 1367–1374, doi: 10.1021/acs.accounts.6b00618 (2017).

28 Kazane, S. A. et al. Self-Assembled Antibody Multimers through Peptide Nucleic Acid Conjugation. J. Am. Chem. Soc. 135, 340–346, doi: 10.1021/ja309505c (2013).

29 Dickgiesser, S. et al. Self-Assembled Hybrid Aptamer-Fc Conjugates for Targeted Delivery: A Modular Chemoenzymatic Approach. ACS Chem. Biol. 10, 2158–2165, doi: 10.1021/acschembio.5b00315 (2015).

30 Leonidova, A. et al. In vivo demonstration of an active tumor pretargeting approach with peptide nucleic acid bioconjugates as complementary system. Chem. Sci. 6, 5601–5616, doi: 10.1039/C5SC00951K (2015).

31 Mahal, L. K., Yarema, K. J. & Bertozzi, C. R. Engineering chemical reactivity on cell surfaces through oligosaccharide biosynthesis. Science 276, 1125–1128, doi: 10.1126/science.276.5315.1125 (1997).

32 Kayser, H. et al. Biosynthesis of a nonphysiological sialic acid in different rat organs, using N-propanoyl-D-hexosamines as precursors. J. Biol. Chem. 267, 16934–16938 (1992).

33 Chandra, R. A., Douglas, E. S., Mathies, R. A., Bertozzi, C. R. & Francis, M. B. Programmable cell adhesion encoded by DNA hybridization. Angew. Chem., Int. Ed. 45, 896–901, doi: 10.1002/anie.200502421 (2006).

34 Shi, P. et al. Poylvalent Display of Biomolecules on Live Cells. Angew. Chem. Int. Ed. 130, 6916–6920, doi: doi: 10.1002/anie.201712596. (2018).

35 Saccà, B. et al. Orthogonal protein decoration of DNA origami. Angew. Chem., Int. Ed. 49, 9378–9383, doi: 10.1002/anie.201005931 (2010).

36 Taylor, M. J., Husain, K., Gartner, Z. J., Mayor, S. & Vale, R. D. A DNA-Based T Cell Receptor Reveals a Role for Receptor Clustering in Ligand Discrimination. Cell 169, 108-119.e120, doi: 10.1016/j.cell.2017.03.006 (2017).

37 Lovendahl, K. N., Hayward, A. N. & Gordon, W. R. Sequence-Directed Covalent Protein–DNA Linkages in a Single Step Using HUH-Tags. J. Am. Chem. Soc. 139, 7030–7035, doi: 10.1021/jacs.7b02572 (2017).

38 Griffin, B. A., Adams, S. R. & Tsien, R. Y. Specific Covalent Labeling of Recombinant Protein Molecules Inside Live Cells. Science 281, 269–272, doi: 10.1126/science.281.5374.269 %J Science (1998).

39 Spagnuolo, C. C., Vermeij, R. J. & Jares-Erijman, E. A. Improved Photostable FRET-Competent Biarsenical-Tetracysteine Probes Based on Fluorinated Fluoresceins. J. Am. Chem. Soc. 128, 12040–12041, doi: 10.1021/ja063212q (2006).

40 Baalmann, M., Best, M. & Wombacher, R. in Noncanonical Amino Acids: Methods and Protocols (ed Edward A. Lemke) 365–387 (Springer New York, 2018).

41 Chen, I., Howarth, M., Lin, W. & Ting, A. Y. Site-specific labeling of cell surface proteins with biophysical probes using biotin ligase. Nat. Methods 2, 99–104, doi: 10.1038/nmeth735 (2005).

42 Reinhardt, U., Lotze, J., Morl, K., Beck-Sickinger, A. G. & Seitz, O. Rapid Covalent Fluorescence Labeling of Membrane Proteins on Live Cells via Coiled-Coil Templated Acyl Transfer. Bioconjugate Chem. 26, 2106–2117, doi: 10.1021/acs.bioconjchem.5b00387 (2015).

43 Reinhardt, U. et al. Peptide-templated acyl transfer: A chemical method for the labeling of membrane proteins on live cells. Angew. Chem., Int. Ed. 53, 10237–10241, doi: 10.1002/anie.201403214 (2014).

44 Litowski, J. R. & Hodges, R. S. Designing heterodimeric two-stranded α-helical coiled-coils. Effects of hydrophobicity and α-helical propensity on protein folding, stability, and specificity. J. Biol. Chem. 277, 37272–37279, doi: 10.1074/jbc.M204257200 (2002).

45 Yano, Y. et al. Coiled-coil tag - Probe system for quick labeling of membrane receptors in living cells. ACS Chem. Biol. 3, 341–345, doi: 10.1021/cb8000556 (2008).

46 Dawson, P. E., Muir, T. W., Clark-Lewis, I. & Kent, S. B. H. Synthesis of Proteins by Native Chemical Ligation. Science 266, 776–779, doi: 10.1126/science.7973629 (1994).

47 Chang, P. V. et al. Copper-free click chemistry in living animals. Proc. Natl. Acad. Sci. U. S. A. 107, 1821–1826, doi: 10.1073/pnas.0911116107 (2010).

48 Rohde, H., Schmalisch, J., Harpaz, Z., Diezmann, F. & Seitz, O. Ascorbate as an Alternative to Thiol Additives in Native Chemical Ligation. ChemBioChem 12, 1396–1400, doi: 10.1002/cbic.201100179 (2011).

49 Haase, C. & Seitz, O. Internal Cysteine Accelerates Thioester-Based Peptide Ligation. Eur. J. Org. Chem. 2009, 2096–2101, doi: 10.1002/ejoc.200900024 (2009).

50 Roskoski, R. The ErbB/HER family of protein-tyrosine kinases and cancer. Pharmacol. Res. 79, 34–74, doi: 10.1016/j.phrs.2013.11.002 (2014).

51 Schlessinger, J. Cell signaling by receptor tyrosine kinases. Cell 103, 211–225, doi: 10.1016/S0092-8674(00)00114-8 (2000).

52 Yamaguchi, T., Murata, Y., Fujiyoshi, Y. & Doi, T. Regulated interaction of endothelin B receptor with caveolin-1. Eur. J. Biochem. 270, 1816–1827, doi: 10.1046/j.1432-1033.2003.03544.x (2003).

53 Mazzuca, M. Q. & Khalil, R. A. Vascular endothelin receptor type B: Structure, function and dysregulation in vascular disease. Biochem. Pharmacol. 84, 147–162, doi: 10.1016/j.bcp.2012.03.020 (2012).

54 Gradišar, H. & Jerala, R. De novo design of orthogonal peptide pairs forming parallel coiled-coil heterodimers. J. Pept. Sci. 17, 100–106, doi: 10.1002/psc.1331 (2011).

55 Raj, A., van den Bogaard, P., Rifkin, S. A., van Oudenaarden, A. & Tyagi, S. Imaging individual mRNA molecules using multiple singly labeled probes. Nat. Methods 5, 877–879, doi: 10.1038/nmeth.1253 (2008).

56 Yurke, B., Turberfield, A. J., Mills, A. P., Simmel, F. C. & Neumann, J. L. A DNA-fuelled molecular machine made of DNA. Nature 406, 605–608, doi: 10.1038/35020524 (2000).

57 Zhang, D. Y. & Seelig, G. Dynamic DNA nanotechnology using strand-displacement reactions. Nat. Chem. 3, 103, doi: 10.1038/nchem.957 (2011).

58 Lotze, J. et al. Time-Resolved Tracking of Separately Internalized Neuropeptide Y<inf>2</inf>Receptors by Two-Color Pulse-Chase. ACS Chem. Biol. 13, 618–627, doi: 10.1021/acschembio.7b00999 (2018).

59 Olson, X., Kotani, S., Yurke, B., Graugnard, E. & Hughes, W. L. Kinetics of DNA Strand Displacement Systems with Locked Nucleic Acids. J. Phys. Chem. B 121, 2594–2602, doi: 10.1021/acs.jpcb.7b01198 (2017).

60 Schweller, R. M. et al. Multiplexed in situ immunofluorescence using dynamic DNA complexes. Angew. Chem., Int. Ed. 51, 9292–9296, doi: 10.1002/anie.201204304 (2012).

61 Bandlow, V. et al. Spatial Screening of Hemagglutinin on Influenza A Virus Particles: Sialyl-LacNAc Displays on DNA and PEG Scaffolds Reveal the Requirements for Bivalency Enhanced Interactions with Weak Monovalent Binders. J. Am. Chem. Soc. 139, 16389–16397, doi: 10.1021/jacs.7b09967 (2017).

62 Liang, S. I. et al. Phosphorylated EGFR Dimers Are Not Sufficient to Activate Ras. Cell Rep. 22, 2593–2600, doi: 10.1016/j.celrep.2018.02.031 (2018).

63 Thomas, F., Boyle, A. L., Burton, A. J. & Woolfson, D. N. A Set of de Novo Designed Parallel Heterodimeric Coiled Coils with Quantified Dissociation Constants in the Micromolar to Sub-nanomolar Regime. J. Am. Chem. Soc. 135, 5161–5166, doi: 10.1021/ja312310g (2013).

