## Extended Data Fiigures for "Live cell PNA labelling enables erasable fluorescence imaging of membrane proteins"

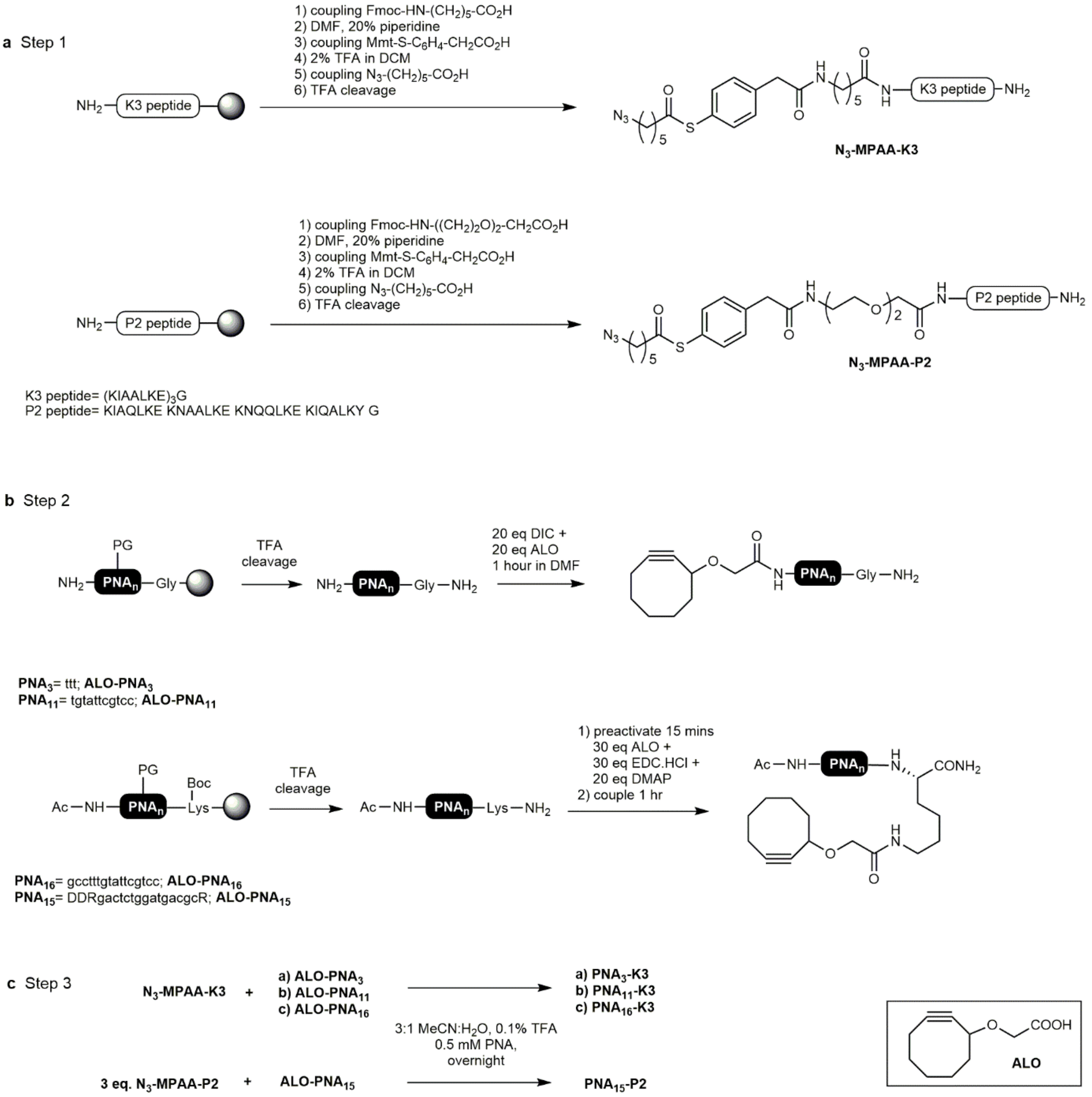


Extended Data Figure 1: Synthesis of thioester-linked K3 and P2 peptide-PNA conjugates via SPPS and strained cycloaddition. a) Solid-phase synthesis affords K3 and P2 peptides containing thioester-linked azido hexanoic acid. b) After solid-phase assembly of PNA strands by using Fmoc/Bhoc-protected PNA monomers, PNA is cleaved by TFA treatment and submitted to an in-solution coupling with ALO (pictured), which provides the ALO-PNA conjugate. c) Strain-promoted azide-alkyne cycloaddition produces the thioester-linked PNA-peptide conjugates.

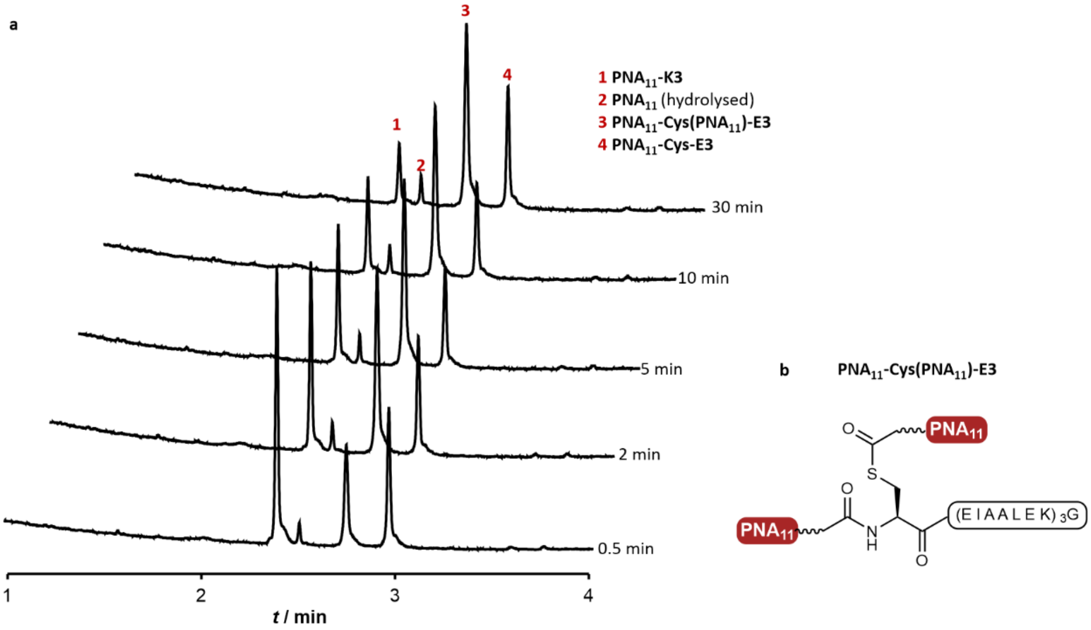


Extended Data Figure 2. Time course experiment of transfer reaction between PNA$_{11}$-K3 and Cys-E3. a) UPLC$^{TM}$ traces of reaction time points. UPLC gradient 20-50 % in 4 min, detection at 260 nM. Experiment was repeated three times with similar results b) Structure of S-acylated side product PNA$_{11}$-Cys(PNA$_{11}$)-E3.

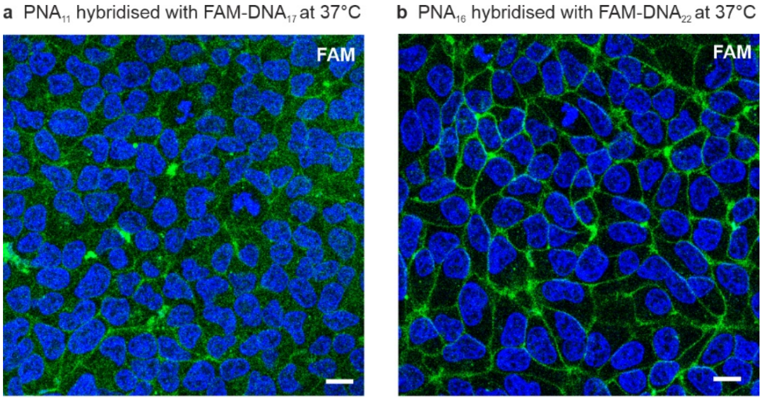


**Extended Data Figure 3.** Analysis of PNA:DNA duplex stability by fluorescence microscope imaging of Cys-E3-h$Y_2$R tagged with $PNA_{11}$ or $PNA_{16}$ and hybridized with complementary FAM-DNA at 37°C. **a)** HEK293-Cys-E3-h$Y_2$R labelled with an 11mer duplex ($PNA_{11}$-K3 tagging and FAM-$DNA_{17}$ hybridisation). **b)** HEK293-Cys-E3-h$Y_2$R labelled with a 16mer duplex ($PNA_{16}$-K3 tagging and FAM-$DNA_{22}$ hybridisation). Stably transfected HEK293-Cys-E3-h$Y_2$R cells[42,58] labelled with Hoechst33342 (shown in blue) were treated with $PNA_{11}$-K3 or $PNA_{16}$-K3 (100 nM) in buffer (HBSS with 0.1 mM TCEP, 20 mM HEPES, pH 7) for 4 min. After washing with basic buffer (200 mM $NaHCO_3$ in DPBS, pH 8.4) for 1.5 min, complementary FAM-DNA (1 µM) containing 6mer overhangs (FAM-$DNA_{17}$ or FAM-$DNA_{22}$) was added for 5 min in 20 mM HEPES buffer. Cells were washed and microscopic studies were performed in OptiMEM at 37°C. Hoechst33342 ($\lambda_{ex}$:365, $\lambda_{em}$:420), 5/6-Carboxyfluorescein (FAM) ($\lambda_{ex}$: 470/40, $\lambda_{em}$:525/50). Scale bar= 10 µm. See Supplementary 9.3 for full method and DNA sequences. Experiment was repeated 3 times independently with similar results.

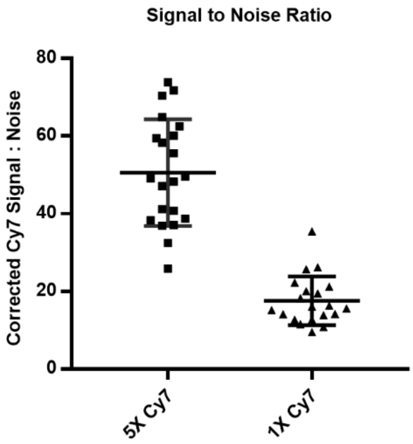


Extended Data Figure 4. Fluorescence microscope analysis of Signal to Noise Ratio (SNR) of $PNA_{15}$ labelled Cys-P1-EGFR-CHO cells stained with either one or five Cy7 fluorophores. After $PNA_{15}$ labelling, cells were incubated with 50 nM Complex I (1x Cy7: adaptor DNA-33mer with a single Cy7-15mer) or 50 nM Complex II (5x Cy7: adaptor DNA-105mer with five Cy7-15mers) in HBSS-BB before washing with HBSS-BB and imaging. From three independent experiments, 6-8 cells were analyzed by line intensity profiles spanning a whole cell. For each line intensity profile, Cy7 or YFP signal was calculated as the max peak height at the membrane regions, and the noise calculated as the standard deviation of the signal from an empty background region. Dot plot is presented as the mean +/- SD with each point representing SNR for one cell (n=20).

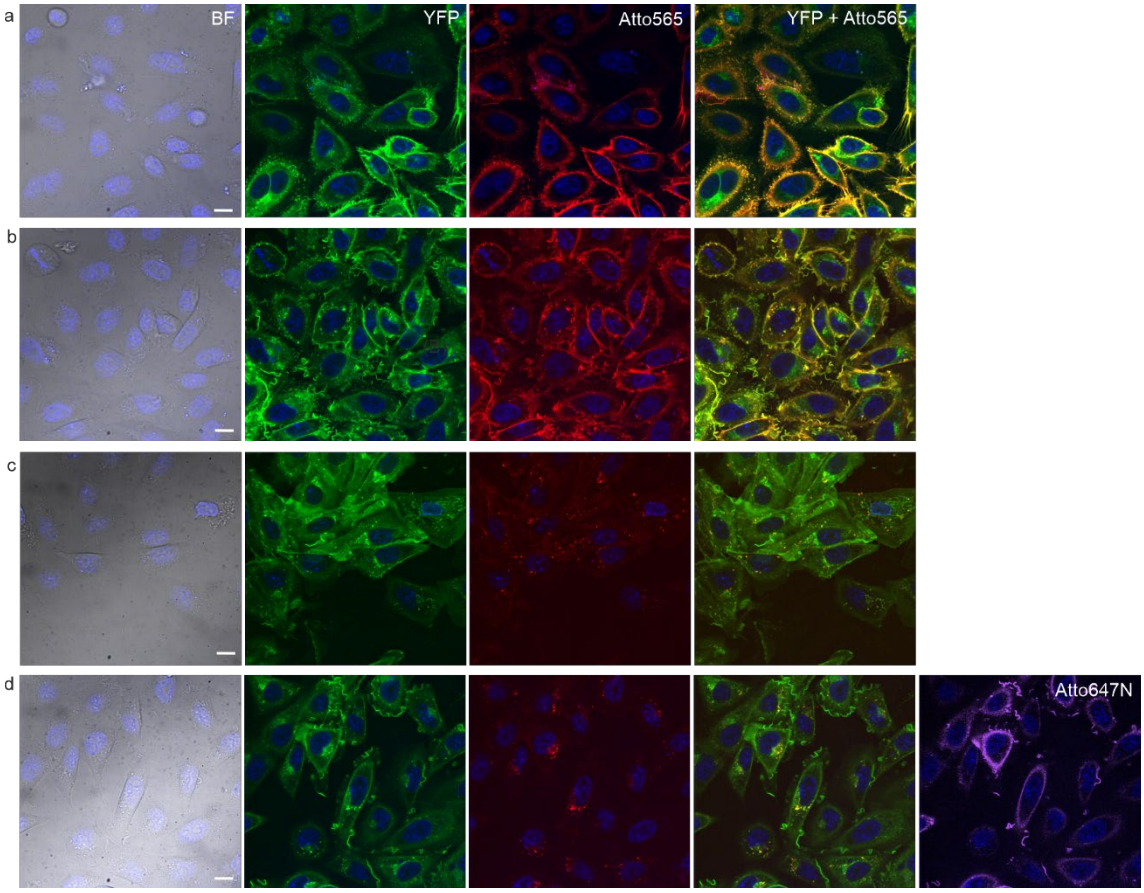


Extended Data Figure 5. Spinning disk confocal microscopy analysis of PNA enabled reversible labelling of Cys-P1-EGFR-eYFP on CHO cells. **a)** After staining of nuclei with Hoechst 33342, serum starved Cys-P1-EGFR-eYFP cells were treated with $PNA_{15}$-P2 in HBSS for 4 minutes. Cells were then incubated with 50 nM **Complex III** (adaptor DNA-105mer + five Atto565-DNA-23mers) in HBSS-BB for 4 min. **b)** Stimulation with EGF (100 nM) for 15 mins. **c)** Toehold mediated strand displacement of Atto565-23mer DNA with 300 nM displacement DNA-23mer in presence of 100 nM EGF for 2 x 5 min in HBSS at 30°C. **d)** Hybridisation with 100 nM Atto647N-DNA-15mer, 3 min. Excitation times: ATTO565: 200 ms YFP: 100 ms, Hoechst 33342:100 ms Atto647N: 300 ms. Diode lasers: Hoechst 33342) 405 nm; YFP) 488 nm; Atto565) 561 nm; Atto647N) 640 nm. Dichroic emission filters Hoechst 33342) $\lambda_{em}$ = 460 ± 50 nm; YFP) $\lambda_{em}$ 470 ± 24 nm; Atto565) $\lambda_{em}$ 600 ± 50 nm. Atto647) $\lambda_{em}$ = 700 ± 75 nm. Scale bar= 10µm. Experiments were repeated 3 times independently with similar results.
