## Supporting Information for "Live cell PNA labelling enables erasable fluorescence imaging of membrane proteins"

### 1 Reagents for chemical synthesis

Standard Fmoc protected amino acids were purchased from *Carbolutions* (St. Ingbert, Germany). PNA monomers; Fmoc-PNA-T-OH, Fmoc-PNA-A(Bhoc)-OH, Fmoc-PNA-C(Bhoc)-OH and Fmoc-PNA-G(Bhoc)-OH were purchased from *LGC Genomics* (Teddington, UK). Fmoc-aminoethyloxyethyloxyacetic acid (Fmoc-Ahz) and [2-[2-(Fmoc-amino)ethoxy]ethoxy]acetic acid (Fmoc-AEEA), were purchased from *Iris Biotech* (Marktredwitz, Germany). TentaGel® Rink amide (TGR) Resin was from *RAPP Polymers* (Tübingen, Germany) with a loading of 0.2 mmol/g. Coupling reagents PyBOP, HCTU, HATU, DIC, EDC.HCl were from *Carl Roth* (Karlsruhe, Germany), Oxyma from *Carbolutions* (Saarbrücken, Germany) and HOBt from *Angene* (Nanjing, China). Triisopropylsilane (TIS) and triscarboxyethylphosphine (TCEP) were purchased from *Aldrich* (Schnelldorf, Germany), piperidine and N,N-Diisopropylethylamine (DIPEA) from *Alfa Aesar* (Karlsruhe, Germany), acetic anhydride from *VWR Chemicals* (Darmstadt, Germany) and N-methylmorpholine (NMM) from *Fisher Bioreagents* (Geel, Belgium). 5,6-Carboxytetramethylrhodamine (TMR) was obtained from *ChemPep Inc.* (Wellington, USA). Buffer components were purchased from *Carl Roth* (Triton™ X-100, Ethylenediaminetetraacetic acid (EDTA)), *Fisher Bioreagents* (Tris), *VWR Chemicals* (NaCl), *Merck* (Sodium hydrogen phosphate, Tween-20 polysorbate), *Iris Biotech* (Dithiotreitol), and *Merck* (SDS, Glycerol). Sinapic acid was purchased from *Acros Organics*. DMF for solid phase synthesis of peptides and Fmoc-PNA-OH monomers were purchased from *VWR Chemicals*. The ALO (2-(cyclooct-2-yn-1-yloxy)acetic acid) building block was prepared as described.[1] The S-Mmt-protected mercaptophenyl acetic was prepared as described.[2] The control peptide C-GLYAEGGK(biotin) was a kind donation from Robert Zitterbart.[3]

### 2 HPLC instruments and mass spectrometry

#### 2.1 HPLC for small scale purification

Purification of products in nanomole amounts (PNA-peptide conjugates) was performed on a HPLC Elite LaChrom from *Merck-Hitachi*. Separation was carried out using a Polaris C18 column (5 µm,C18-A 100 x 4.6) from *Agilent Technologies* with a flow rate of 0.8 mL / min at 55 °C. The mobile phase was a mixture of solutions A (98.9% water, 1.0% ACN, 0.1% TFA) and B (98.9% ACN, 1.0% water, 0.1% TFA). For detection, Elite LaChrom L-2450 from *Merck Hitachi* was used, with a spectral measuring range of 200 - 350 nm.

#### 2.2 HPLC for large scale purification

Semi-preparative HPLC purification was performed on a 110 Series HPLC system from *Agilent Technologies.* Separation was carried out using a Polaris C18 column (5 µm, 250 x 10, pore size: 220 Å) from *Varian* at a flow rate of 6 mL / min. The mobile phase was composed as a mixture of solutions A (98.9% water, 1.0% ACN, 0.1% TFA) and B (98.9% ACN, 1.0% water, 0.1% TFA). Detection using a UV variable wavelength detector was carried out at wavelengths λ = 210 nm and λ = 260 nm.

#### 2.3 UPLC™:

***UPLC™-MS:*** Analysis of products was performed using an Aquity UPLC™-MS system from *Waters.* The mass spectrometry analysis was carried out using a quadrupole analyzer with electron spray ionization (ESI); detection via a diode array detector (DAD) at the wavelengths λ = 210 nm and λ = 260 nm. The stationary phase was a *Waters* Acquity UPLC™ C18 column (1.7 µm, 50 x 2.1 mm). The mobile phase was a mixture of solutions A (98.9% water, 1.0% ACN, 0.1% TFA) and B (98.9% ACN, 1.0% water, 0.1% TFA), which was pumped at the flow rate of 0.5 mL / min, at 50 ° C.

***FLR-UPLC™***: Analysis of products by fluorescence was performed using an ACQUITY UPLC™ Fluorescence (FLR) Detector. The stationary phase was a *Waters* Acquity UPLC™CSH C18 column (1.7 µm, 50 x 2.1 mm). The mobile phase was a mixture of solutions A (98.9% water, 1.0% ACN, 0.1% TFA) and B (98.9% ACN, 1.0% water, 0.1% TFA), which was pumped at the flow rate of 0.5 mL / min, at 50 ° C.

#### 2.4 Matrix-assisted laser desorption ionization-time of flight mass spectrometry

MALDI-TOF mass spectrometry was performed by using a *Shimadzu* Biotech Axima Confidence instrument in linear mode. Ionization was carried out with a nitrogen laser. In each spectrum 10 profiles of 50 shots each were used, with a relative laser intensity of approx. 90 in positive measuring mode. The matrix was a solution of sinapinic acid (10 mg / mL in acetonitrile : $H_2O$: 1:1 with 0.1% TFA).

### 3 Synthesis methods

#### 3.1 General peptide synthesis

Peptide synthesis was carried out in a 10 µmol or 25 µmol scale using a MultipepRS synthesizer from *Intavis* (Köln, Germany). The appropriate amount of TantaGel® Rink amide (TGR) resin (approx. 0.2 mmol/g) was allowed to swell in DMF for at least 30 minutes prior to synthesis. Fmoc/tBu SPPS strategy was employed, using Fmoc-protected amino acids. The first deprotection step of the resin was used for Fmoc monitoring and the calculated value for reaction scale used to obtain final synthesis yields.

*Fmoc Deprotection:* The resin was treated with a mixture of DMF:piperidine (4:1) twice for 5 minutes and again for 3 minutes before washing three times with DMF.

*Coupling:* Coiled-coil peptides E3, K3, P1 and P2 were prepared by using 5 eq. Fmoc protected amino acid, 5 eq. Oxyma, 4.5 Eq. HCTU and 10 eq NMM in each coupling step. The following stock solutions were prepared: 0.4 M

Fmoc-amino acid with Oxyma in DMF; 0.4 M HCTU in DMF; 4M NMM in DMF, and amino acids were activated immediately before coupling. Single couplings for 30 minutes were used up to the 8$^{th}$ amino acid, after which double couplings for 20 minutes each were used. The resin was then washed with DMF 3 times after coupling

*Capping:* Unreacted terminal amines from the coupling step were capped by treatment with a solution of acetic anhydride: 2,6-lutidine: DMF (5:6:89) for 5 minutes before washing 3 times with DMF.

*5(6)-Carboxytetramethylrhodamine (TMR) coupling:* TMR was coupled by hand using 4 eq. of 5(6)-Carboxytetramethylrhodamine, 4 eq. PyBOP and 8 eq. NMM in DMF for 30 mins. The coupling was repeated, and no capping step was carried out. The resin was washed 5 times each with DMF and DCM.

*N-terminal cysteine coupling:* N-terminal cysteine was coupled by hand using 4 eq. Boc-L-Cys(Mmt)-OH, 3.6 eq. HATU, 4 eq. DIPEA for 20 mins. Mmt and Boc protecting groups were removed during the final global deprotection and cleavage.

#### 3.2 General PNA synthesis

Synthesis of PNA oligomers was carried out on TGR resin with a loading of approx. 0.2 mmol/g using using a ResPep synthesizer from *Intavis* (Köln, Germany). An appropriate amount of resin was swollen in a syringe reactor for 30 minutes prior to synthesis. The first deprotection step of the resin was used for Fmoc monitoring and the calculated value for reaction scale used to calculate the final yield.

*Fmoc Deprotection:* The resin was treated with a mixture of DMF:piperidine (4:1) twice for 5 minutes before washing 5 times with DMF.

*Coupling:* 4 eq. Fmoc-protected PNA monomer, 4 eq. PyBOP and 8 eq. NMM in DMF was used. The volume of DMF was kept as low as the solubility of the PNA monomer allowed, however, no less than 100 µL was used. The activated PNA monomer was reacted for 30 minutes. Double couplings were carried out throughout, and coupling was followed by washing 5 times with DMF

*Capping:* Unreacted terminal amines from the coupling step were capped by treatment with a solution of acetic anhydride: 2,6-lutidine: DMF (5:6:89) for 5 minutes followed by washing 5 times each with DMF.

#### 3.3 Peptide and PNA cleavage from the resin

After synthesis, peptides and PNA oligomers were cleaved from the resin together with side group deprotection. Cleavage was carried out in syringe reactors and prior to cleavage resin was either dried under reduced pressure or washed 10 times with DCM.

*Cleavage from TGR resin*: Peptides and PNA oligomers prepared on TGR resin were cleaved and deprotected using TFA:TIS:$H_2O$ (96:2:2) for 2 hours, before rinsing twice with 1 mL TFA for 30 minutes. For the P2 thioester, cleavage was limited to 2 hours maximum and the resin was rinsed once with DCM for 10 mins after cleavage, and precipitated directly without concentration. For the other peptides/PNAs, the combined TFA fractions were concentrated to around 100 µL per 1 µmol oligomer. The product was precipitated in ten times the volume of cold ether (-20 °C), centrifuged for 15 minutes at 4 °C and the precipitate dried under compressed air. The crude product obtained was purified by HPLC.

#### 3.4 Synthesis of thioester peptides

The Fmoc-K3 and Fmoc-P2 peptides were assembled on TGR resin as previously described. After Fmoc deprotection, to the N terminus were coupled by hand: Fmoc-aminohexanoic acid, or 2-(2-(Fmoc-amino)ethoxy)ethoxy]acetic acid (Fmoc-AEEA-OH), followed by Mmt-mercaptophenylacetic acid (Mmt-MPAA-OH) and finally azidohexanoic acid to generate the thioester. In between coupling steps was a capping step according to the standard protocol, followed by deprotection of Fmoc or Mmt.

*Coupling of the aminohexanoic acid, 2-(2-amino)ethoxy)ethoxy acetic acid and mercaptophenylacetic acid:* 4 eq. Fmoc-aminohexanoic acid, Fmoc AEEA-OH or Mmt-MPAA-OH; 3.6 eq. HCTU and 8 eq. NMM in DMF was used. After double couplings for 30 minutes each, and test cleavage had ensured full conversion, the resin was washed with DMF, DCM and finally DMF again before capping. The resin was washed with DMF 5 times and DCM 10 times.

*Removal of the Mmt Protecting Group on MPAA:* Resin was treated DCM:TFA:TIS (96:3:1) 5 times for 2 minutes each, until the discard was colourless, before washing 5 times with DCM and 5 times with DMF.

*Coupling of azidohexanoic acid to form the thioester:* A mixture of 4 eq. azidohexanoic acid with 3.6 eq HCTU and 8 eq. NMM in DMF was added to the resin and reacted for 30 minutes before washing with DMF 5 times and DCM 10 times.

*ALO coupling to $PNA_3$, $PNA_{11}$, $PNA_{16}$:* A PNA oligomer containing an unprotected N-terminus or an unprotected lysine side chain was prepared by Fmoc-based solid phase synthesis as described. The crude product obtained after TFA cleavage and ether precipitation was dissolved in DMF. For coupling of aryl-less cycloctyne (ALO), a solution of 2-(cyclooct-2-ine-1-yloxy)acetic acid (20 eq) and DIC (20 eq) in DMF was preactivated for 10 min with shaking and added to the PNA solution (approx. 10 mM final concentration). After 1 h the mixture was purified by RP-HPLC.

*ALO coupling to $PNA_{15}$:* A $PNA_{15}$ oligomer containing an unprotected lysine side chain was prepared by Fmoc-based solid phase synthesis as described. The crude product obtained after TFA cleavage and ether precipitation was was purified by RP-HPLC and dissolved in DMSO. For coupling of aryl-less cycloctyne (ALO), a solution of ALO (2-(cyclooct-2-ine-1-yloxy)acetic acid, 30 eq) and EDC.HCl (30 eq) with DMAP (20 eq) was preactivated in DMSO for 15 min with shaking and added to the PNA solution (approx. 10 mM final concentration). After 1 h the mixture was purified by RP-HPLC.

Strain promoted Azide-Alkyne cycloaddition (SPAAC): The purified thioester-linked azido-peptide and the ALO-PNA conjugate were dissolved in a 3:1 stoichiometry in MeCN: H2O (3:1) with 0.1% TFA and allowed to react overnight at 0.5 – 1 mM concentration. Purification was performed by RP-HPLC.

### 4 Compound Characterization

**Cys-E3 peptide**, Cys-$(EIAALEK)_3G-NH_2$

The peptide was prepared as previously described.[4]

**Cys-K3.9,** Cys-AEEA-(KIAALKE)3-KIAALG-$NH_2$

Yield: 6% (362 nmol). $\varepsilon_{214nm=}$ 27227 L·mol$^{-1}$cm$^{-}$. LC gradient: 10-70% B in 4 min, 210 nm. ESI-MS: *m/z* = 1027.94, 771.28, 617.32. ($C_{140}H_{255}N_{37}O_{37}S$ 3079.05 g·mol$^{-1}$ calculated: $[M+3H]^{3+}$ = 1027.35, $[M+4H]^{4+}$ =770.76), $[M+5H]^{5+}$ =6176.81.)

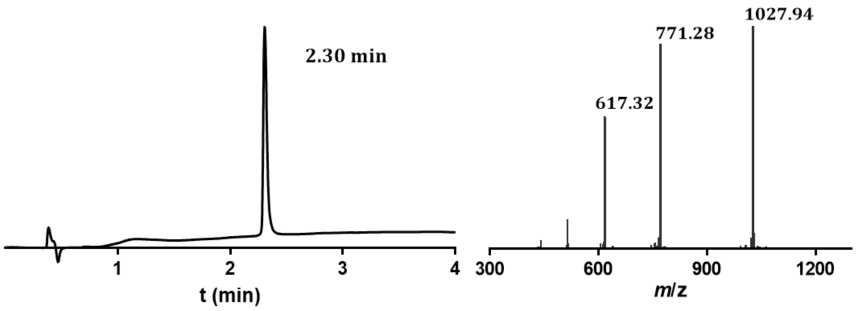

**Cys-P1-K(TMR),** Cys-EIQALEE ENAQLEQ ENAALEE EIAQLEY G K(TMR)-$NH_2$

Yield: 9.9% (990 nmol). $\varepsilon_{260m=}$ 32484 L·mol$^{-1}$cm$^{-1}$. LC gradient: 20-70% B in 4 min, 280 nm. ESI-MS: *m/z* = 658.93, 790.03, 987.28. ($C_{174}H_{257}N_{41}O_{62}S_{1,}$ 3945.09, calculated: $[M+6H]^{6+}$ =658.52; $[M+5H]^{5+}$ =790.03; $[M+4H]^{4+}$ =987.28; $[M+3H]^{3+}$**.**

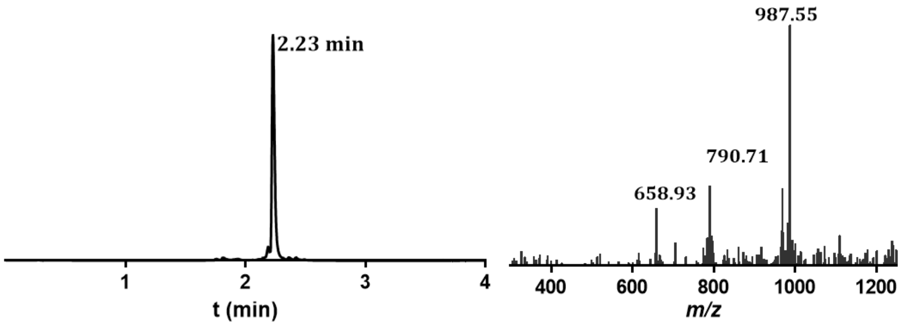

**$N_3$-C($N_3$)-E3,** $N_3(CH_2)_5$(CO)-Cys((CO)$(CH_2)_5N_3$)-(EIAALEK)$_3$G-$NH_2$

Yield: 7% (350nmol). $\varepsilon_{214nm=}$24083. LC gradient: 40-90% B in 4 min, 210 nm. ESI-MS: *m/z* = 907.57. ($C_{119}H_{203}N_{33}O_{37}S$, 2718,47 g·mol$^{-1}$, calculated: $[M+2H]^{2+}$ =907.16).

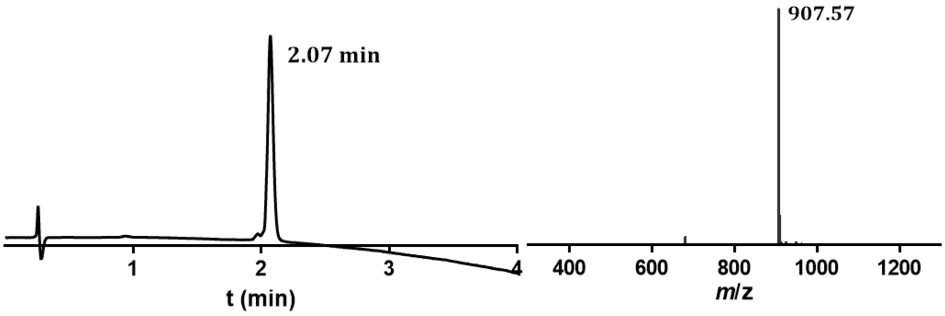

**$N_3$-MPAA-K3**, $N_3(CH_2)_5$(CO)-MPAA-Ahx-(KIAALKE)$_3$G-$NH_2$

Yield: 48% (206 nmol) based on loading of resin with K3 peptide. $\varepsilon_{214nm}$ = 28396 L·mol$^{-1}$cm$^{-1}$. LC gradient: 3-80% B in 4 min, 210 nm. MALDI-TOF-MS: *m/z* = 2739. ($C_{127}H_{221}N_{33}O_{31}S$, $[M+H]^+$, calculated: 2738.39 g·mol$^{-1}$).

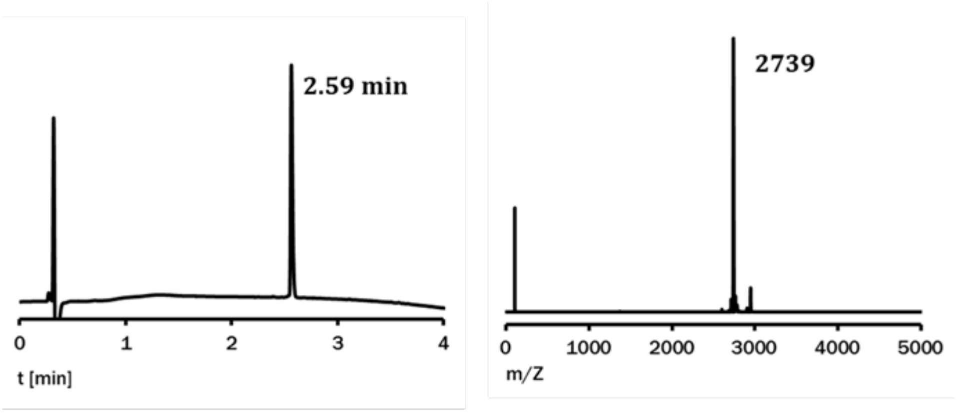

**$N_3$-MPAA-P2,** $N_3(CH_2)_5$(CO)-MPAA-AEEA-KIAQLKE KNAALKE KNQQLKE KIQALKY G-$NH_2$

Yield: 8% (1.5 µmol) $\varepsilon_{214nm=}$ 41008 L·mol$^{-1}$cm$^{-1}$ . LC gradient: 20-80% B in 4 min, 210 nm. ESI-MS: *m/z* = 948.56, 759.29 ($C_{170}H_{288}N_{48}O_{47}S$ 3788.48, calculated: $[M+4H]^{4+}$ = 948.12, $[M+5H]^{5+}$ =758.70).

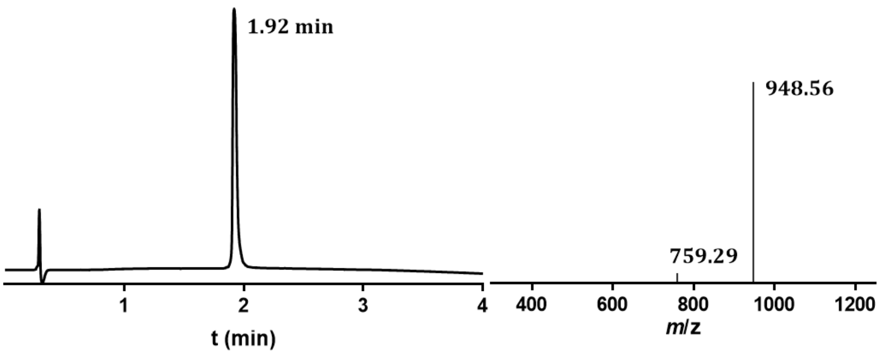

**ALO-PNA$_3$**, ALO-ttt-Gly-NH$_2$

Yield: 80% (800nmol). $\varepsilon_{260nm}$ = 25800 L·mol$^{-1}$cm$^{-1}$. LC gradient: 3-80% B in 4 min, 260 nm. ESI-MS: *m/z* = 519.4, 1038.4. ($C_{45}H_{60}N_{14}O_{15}$, 1307.0, calculated: $[M+H]^{1+}$ = 1038.0, $[M+2H]^{2+}$ =519.5).

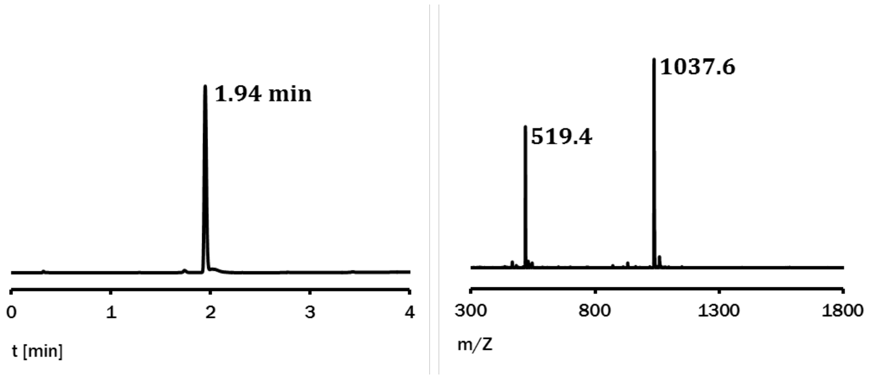

**ALO-PNA$_{11}$**, ALO- tgt att cgt cc-Gly-NH$_2$

Yield: 36% (360 nmol). $\varepsilon_{260nm}$ = 99800 L·mol$^{-1}$cm$^{-1}$ . LC gradient: 3-80% B in 4 min, 260 nm. (Peak shape may be due to an accidental resolution of isomers formed after the strain-promoted click reaction). MALDI-TOF-MS: *m/z* = 3180. ($C_{130}H_{166}N_{58}O_{40}$, $[M+H]^+$, calculated: 3181.08 g·mol$^{-1}$).

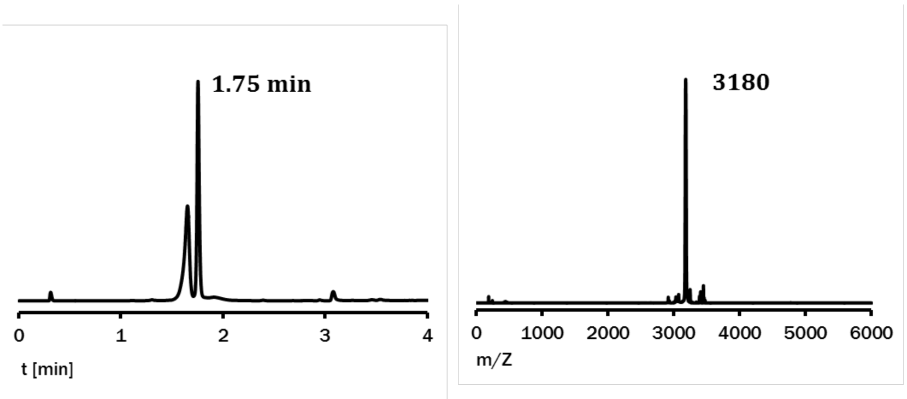

**ALO-PNA$_{16}$**, Ac-gcc ttt gta ttc gtc c-Lys(ALO)-NH$_2$

Yield: 13% (480 nmol). $\varepsilon_{260nm}$ = 141900 L·mol$^{-1}$cm$^{-1}$ . LC gradient: 3-80% B in 4 min, 260 nm. ESI-MS: *m/z* = 925.2, 1155.7 ($C_{117}H_{207}N_{29}O_{31}$, 4620.50 g·mol$^{-1}$, calculated: $[M+4H]^{4+}$ = 1156.1, $[M+5H]^{5+}$ =925.1).

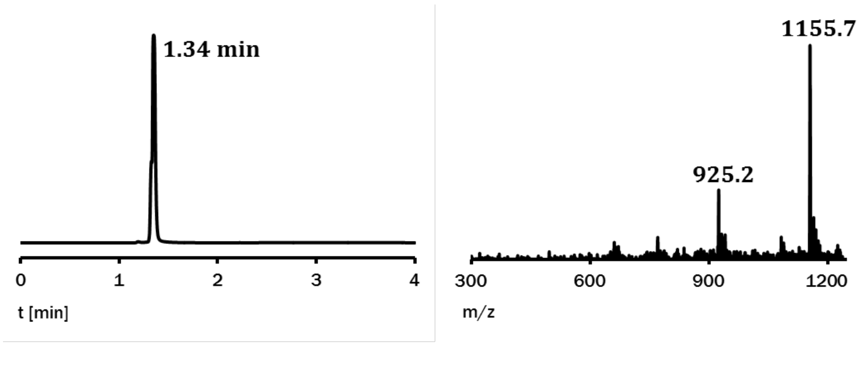

**ALO-PNA$_{15}$**, Ac-D D R $^{N}$gac tct gga tga cgc$^{C}$ R-Lys(ALO)-NH$_2$

Yield: 2% (20 nmol) $\varepsilon_{260nm=}$ 151800 L·mol$^{-1}$cm$^{-1}$. LC gradient: 10-60% B in 4 min, 280 nm. ESI-MS: *m/z* = 931:9, 996:91 ($C_{199}H_{261}N_{101}O_{57}$, 4978.97 g·mol$^{-1}$, calculated: $[M+5H]^{5+}$ = 966.79, $[M+6H]^{6+}$ =830.83).

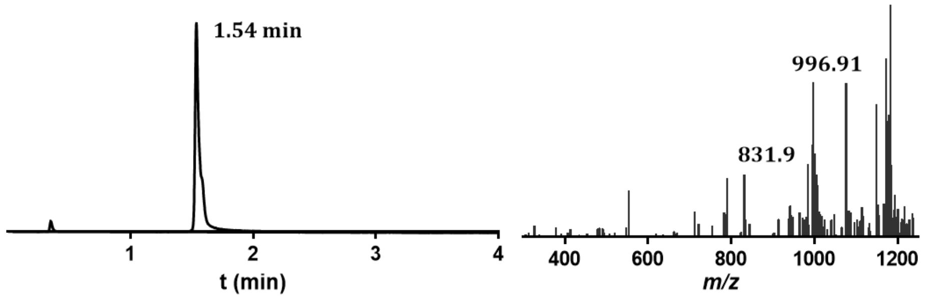

**PNA$_3$-K3**, Gly-$^{C}$ttt$^{N}$-ALON$_3$(CH$_2$)$_5$(CO)-MPAA-Ahx-(KIAALKE)$_3$G-NH$_2$

Yield: 20% (6 nmol). $\varepsilon_{260nm}$ = 25800 L·mol$^{-1}$cm$^{-1}$. LC gradient: 3-80% B in 4 min, 260 nm. MALDI-TOF-MS: *m/z* = 3777. ($C_{172}H_{281}N_{47}O_{46}S$, $[M+H]^{+}$, calculated: 3775.42 g·mol$^{-1}$

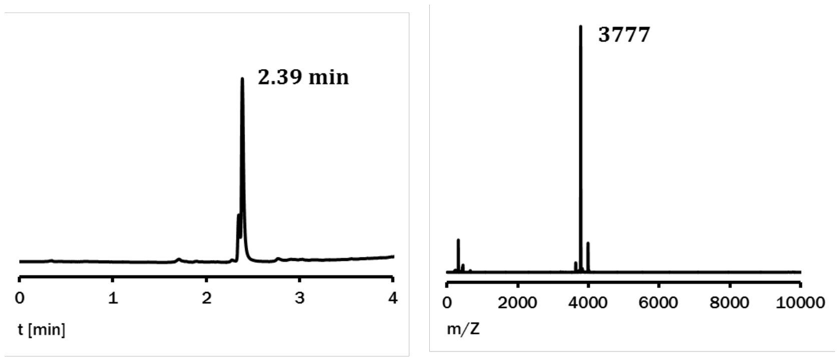

Despite the impurity visible on the UPLC chromatogram, it was decided not to further purify this compound.

**PNA$_{11}$-K3,** Gly-$^{C}$cc tgc tta tgt$^{N}$-Lys(ALON$_3$(CH$_2$)$_5$(CO)-MPAA-Ahx-(KIAALKE)$_3$G-NH$_2$)

Yield: 29% (8.6 nmol). $\varepsilon_{260nm}$ = 99800 L·mol$^{-1}$cm$^{-1}$. LC gradient: 3-80% B in 4 min, 260 nm. MALDI-TOF-MS: *m/z* = 5922. ($C_{257}H_{387}N_{91}O_{71}S$, $[M+H]^{+}$, calculated: 5919.46 g·mol$^{-1}$)

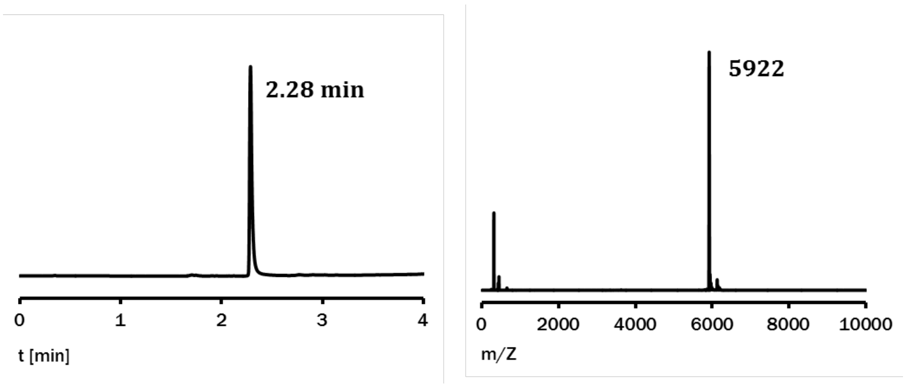

**$PNA_{16}$-K3**, Ac-$^{N}$gcc ttt gta ttc gtc c$^{C}$-Lys(ALO)$N_3(CH_2)_5$(CO)-MPAA-Ahx-(KIAALKE)$_3$G-$NH_2$

Yield: 4% (16 nmol). $\varepsilon_{260nm}$ = 141900 L·mol$^{-1}$cm$^{-1}$. LC gradient: 3-80% B in 4 min, 260 nm. ESI-MS: *m/z* = 1227.1, 1052.2, 921.1 ($C_{316}H_{465}N_{117}O_{89}S$, 7358.88 g·mol$^{-1}$, calculated: $[M+6H]^{6+}$ = 1227.5, $[M+7H]^{7+}$ =1052.3, $[M+8H]^{8+}$ = 920.9.

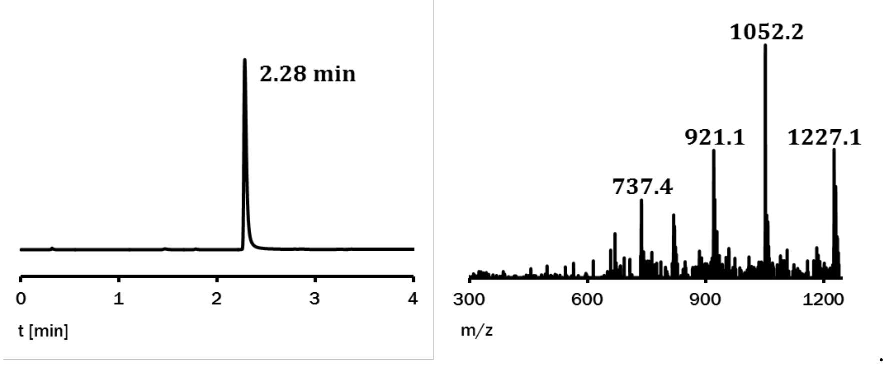

**$PNA_{15}$-P2,** Ac- DDR $^{N}$gac tct gga tga cgc$^{C}$ R-Lys((ALO)$N_3(CH_2)_5$(CO)-MPAA-AEEA-KIAQLKE KNAALKE KNQQLKE KIQALKY G-$NH_2$)

Yield: 7% (10 nmol). $\varepsilon_{260nm=}$ 154423 L·mol$^{-1}$cm$^{-1}$. LC gradient: 3-80% B in 4 min, 280 nm. ESI-MS: *m/z* = 798.58, 877.63, 975.68, 1097.01 ($C_{369}H_{549}N_{149}O_{104}S$, 8768.28 g·mol$^{-1}$, calculated: $[M+8H]^{8+}$ = 1096.93, $[M+9H]^{9+}$ =975.16), $[M+10H]^{10+}$ =877.75, $[M+11H]^{11+}$ = 798.04)

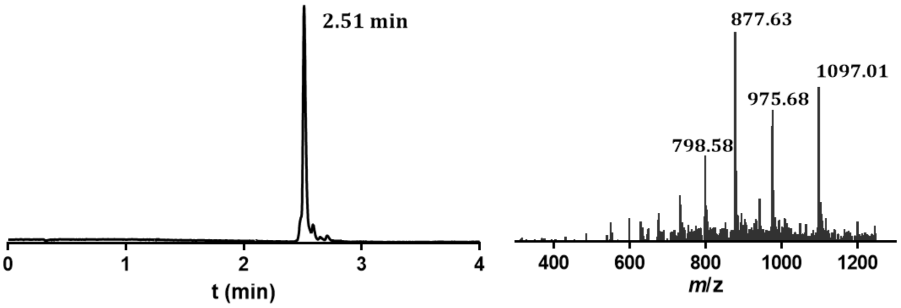

**TMR-$PNA_{12}$,** TMR-NH$(CH_2)_5$(CO)a gga cga ata ca-Gly-$NH_2$

Yield: 10% (250 nmol). $\varepsilon_{260nm}$ = 159539L·mol$^{-1}$cm$^{-1}$. LC gradient: 3-80% B in 4 min, 260 nm. MALDI-TOF-MS: *m/z* = 3899 ($C_{162}H_{195}N_{80}O_{38}$, $[M+H]^+$, calculated: 3896.42 g·mol$^{-1}$)

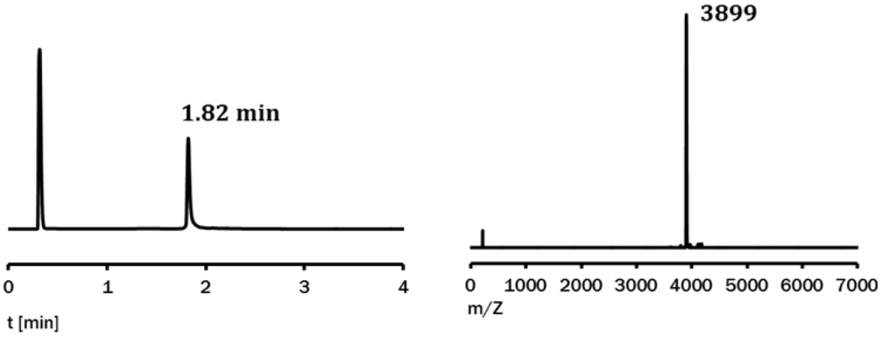

**TMR-PNA₁₆**, TMR-NH($CH_2$)$_2$O($CH_2$)$_2$(CO)-c gag caa gcg cct gag-Gly-$NH_2$

Yield: 6% (170 nmol). $\varepsilon_{260nm}$ = 198200 L·mol$^{-1}$cm$^{-1}$. LC gradient: 3-80% B in 4 min, 260 nm. ESI-MS: *m/z* = 992.7, 827.5 ($C_{202}H_{243}N_{104}O_{52}$, 4959.75 g·mol$^{-1}$, calculated: [M+5H]$^{5+}$ = 993.0, [M+6H]$^{6+}$ =827.6

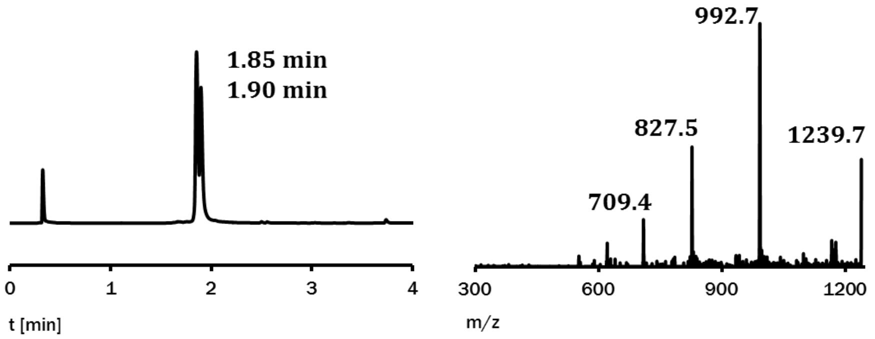

### 5 Transfer reactions

#### 5.1 PNA$_n$-K3

Time dependent studies of transfer reactions with PNA-peptide conjugates were carried out as independent triplicates, with aliquots taken at defined time points and analysed by UPLC™ and later verified by UPLC-MS. The buffer was prepared by addition of TCEP (from a 100 mM stock to a final concentration of 1 mM) to degassed phosphate buffer (100 mM $Na_2HPO_4$, pH 7). The cysteinyl peptide was dissolved in the buffer to a final concentration of 2.5 µM and incubated for 10 min. Timing began with the addition of the PNA$_n$-K3 to a final concentration of 2.5 µM. Aliquots of 20 µL were withdrawn at the respective time and quenched upon addition of 0.4 µL TFA. A volume of 10 µL was injected into the UPLC™. The area under the relevant peaks were weighted by taking the extinction coefficients of the respective compound into consideration.

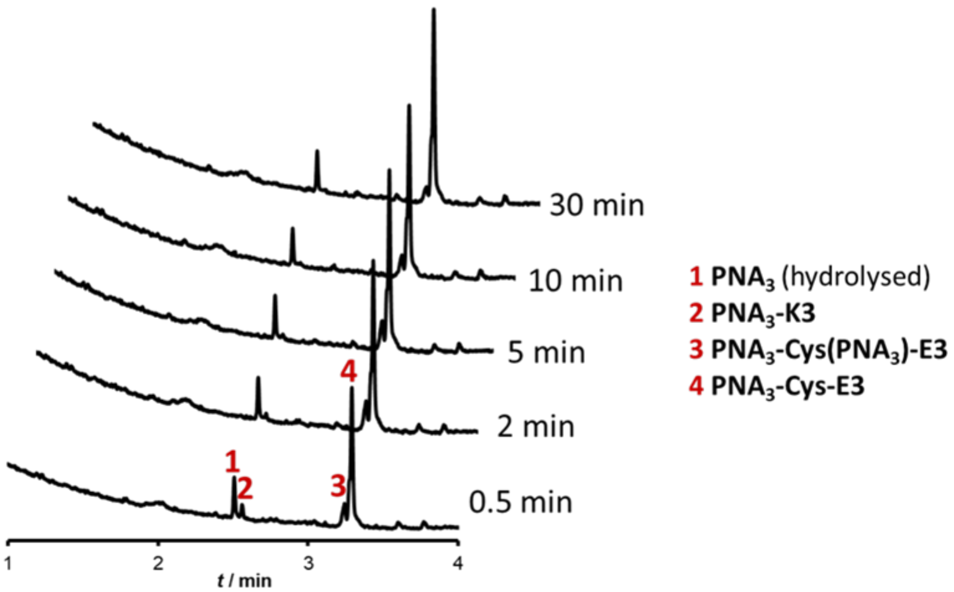

Figure 5-1 **Time course experiment of transfer reaction between PNA$_3$-K3 and Cys-E3.** UPLC$^{TM}$ traces of reaction time points were recorded as described in section 5.1. Peptide concentration: 2.5 µM. UPLC gradient 0-60 % in 4 min, 260 nM. Experiment was repeated three times with similar results.

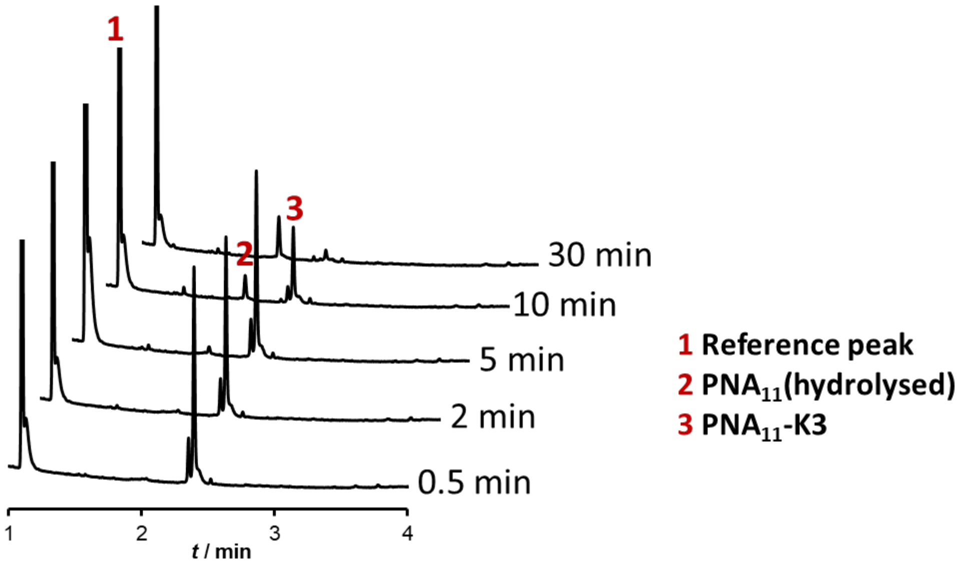

Figure 5-2 **Control experiment of transfer reaction between PNA$_{11}$-K3 and random cysteine peptide (C-GLYAEGGK(biotin).** UPLC$^{TM}$ traces of reaction time points were recorded as described in section 5.1. No transfer product was observed under the given conditions. UPLC$^{TM}$ gradient 0-60 % in 4 min, 260 nM. Experiment was repeated 3 times with similar results.

##### 5.1.1 Stability of S-acylated NCL product

**$N_3$-C($N_3$)-E3** was used as a model compound to estimate the stability of the double N- and S-acylated side product **$PNA_{11}$-Cys($PNA_{11}$)-E3**, when treated with reagents which induce thiolysis of the thioester.

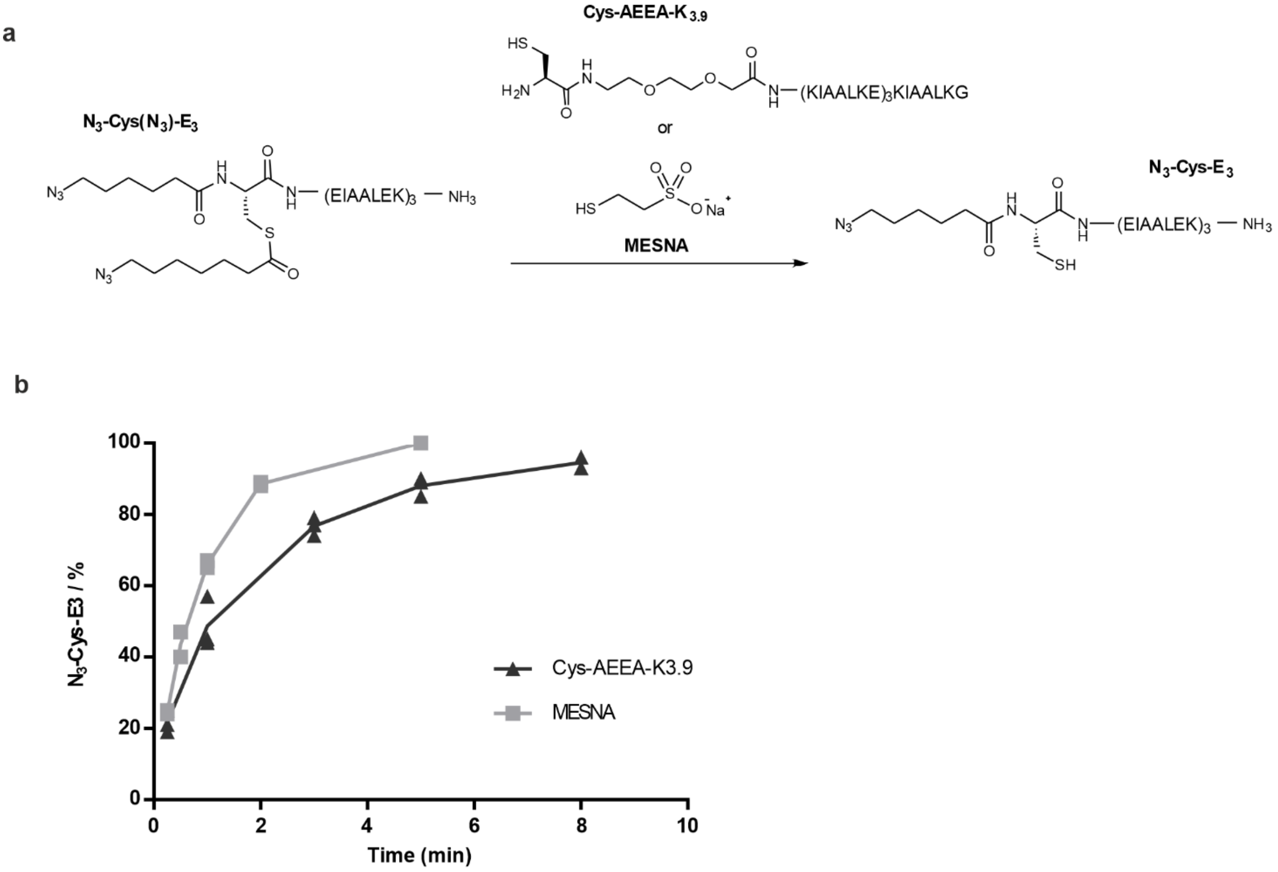

Figure 5-3 **Cleavage of the thioester in $N_3$-C($N_3$)-E3 upon treatment with MESNA or Cys-AEEA-K3.9. a)** Reaction scheme **b)** Rate of formation of **$N_3$-Cys-E3**. Treatment of 25µM of **$N_3$-C($N_3$)-E3** with either (i) 25µM (equimolar) **Cys-AEEA-K3.9** in 100mM PBS, pH 7.4 or (ii) 20 mM (excess) MESNA with 10 mM TCEP in 100mM PBS, pH 7.4. Data is presented with points representing independent replicates, with a line connecting the mean of the replicates (Independent replicates; Cys-AEEA-K3.9, n=3; MESNA, n=2.)

#### 5.2 $PNA_{15}$-P2

Reaction time courses of transfer reactions with **$PNA_{15}$-P2** and **Cys-P1-K(TMR)** were analysed by FLR-UPLC. Reactions were carried in phosphate buffer (200 mM $Na_2HPO_4$, 2 mM TCEP, pH 7.4, 30°C). Prior to experiments, 200 nM **Cys-P1-K(TMR)** was shaken for 10 minutes in buffer before addition of 500 nM **$PNA_{15}$-P2.** Aliquots were taken at different time points and diluted by 50 % upon addition of 4% TFA in water.

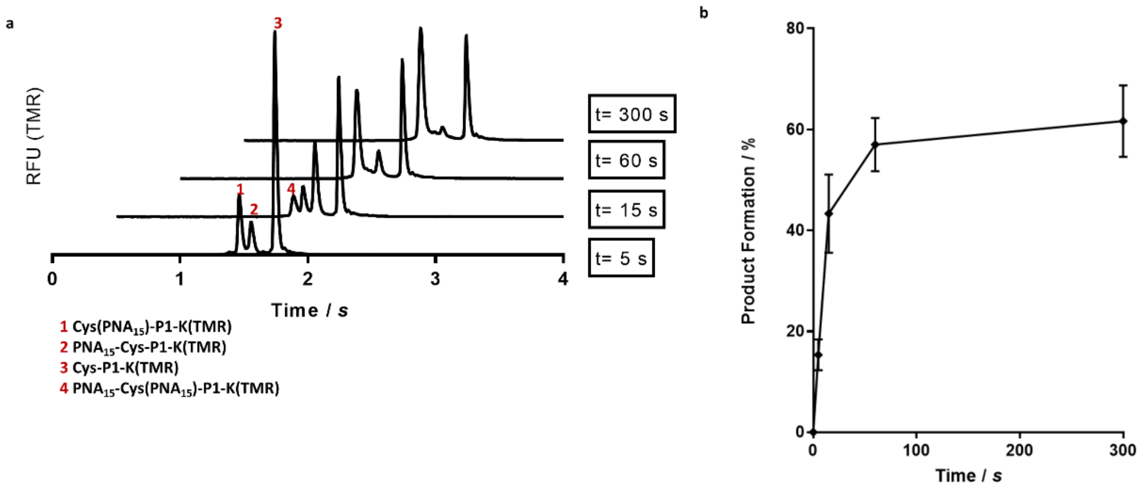

Figure 5-4 **Time course experiment of transfer reaction between $PNA_{15}$-P2 and C-P1-K(TMR). a)** FI-UPLC™ traces of reaction time points were recorded as described in section 5.2. Peptide concentration: 200 nM **Cys-P1-K(TMR)**, 500 nM **$PNA_{15}$-P2**. FI-UPLC™ gradient 10-70 % in 4 min, Ex: 550 nm, Em: 580 nm. Experiment was repeated three times with similar results. **b)** Rate of formation of $PNA_{15}$-Cys-P1-K(TMR) and $PNA_{15}$-Cys($PNA_{15}$)-P1-K(TMR) products calculated from area peaks shown in a). Data is presented as the mean +/- SD of n=3 independent measurements.

### 6 PNA and DNA Sequences

Table 6-1 **DNA and PNA Sequences**. Bold-faced nucleotides remain unpaired after hybridization. Toehold portion of DNA shown in italics.

| **Name** | **DNA/PNA** (5′->3′ or N->C) **used in Figure 1 and 2 in main article** |
|---|---|
| $PNA_3$ | ttt Gly |
| $PNA_{11}$ | tgt att cgt cc Gly |
| $PNA_{16}$ | gcc ttt gta ttc gtc c Lys |
| Atto565 DNA-16mer | Atto565-GGA CGA ATA CAA AGG C |
| Cy3 DNA-16mer | Cy3-GGA CGA ATA CAA AGG C |
| Cy3 DNA-12mer | Cy3-A GGA CGA AAT ACA |
| TMR DNA-20mer | TMR-GC CAC GAG CAA GCG CCT GAG |
| DNA-34mer | GGA CGA ATA CAA AGG C **AT** CTC AGG CGC TTG CTC G |
| DNA-66mer | GGA CGA ATA CAA AGG C **AT** (CTC AGG CGC TTG CTC G)$_3$ |
| TMR PNA-16mer | TMR-c gag caa gcg cct gag Gly |
| TMR PNA-12mer | TMR- a gg acg aat aca Gly |
| FAM-$DNA_{17}$ | FAM-GG ACG AAT ACA CAT CTC |
| FAM-$DNA_{22}$ | G CGG CAC GGA AAC ATA AGC AGG-FAM |

| **DNA/PNA (5‘->3‘ or N->C) used in Figure 3, 4, 5 in main article** | |
|---|---|
| $PNA_{15}$ | Asp Asp Arg gac tct gga tga cgc Arg Lys |
| Atto565 DNA-15mer | ATTO565-GCG TCA TCC AGA GTC |
| Cy7 DNA-15mer | Cy7- GAC ACC ACT TAC CAG |
| Atto647N DNA-15mer | Atto647N- GAC ACC ACT TAC CAG |
| DNA-33mer (multilabelling) | GCG TCA TCC AGA GTC **CTA** CTG GTA AGT GGT GTC |
| DNA-105mer (multilabelling) | GCG TCA TCC AGA GTC **CTA** CTG GTA AGT GGT GTC **CTA** CTG GTA AGT GGT GTC **CTA** CTG GTA AGT GGT GTC **CTA** CTG GTA AGT GGT GTC **CTA** CTG GTA AGT GGT GTC |
| Atto565 DNA-23mer (erasable label) | ATTO565- GAC ACC ACT TAC CAG *ATA GCA CA* |
| Displacement DNA-23mer (erasure) | *TGT GCT AT*C TGG TAA GTG GTG TC |

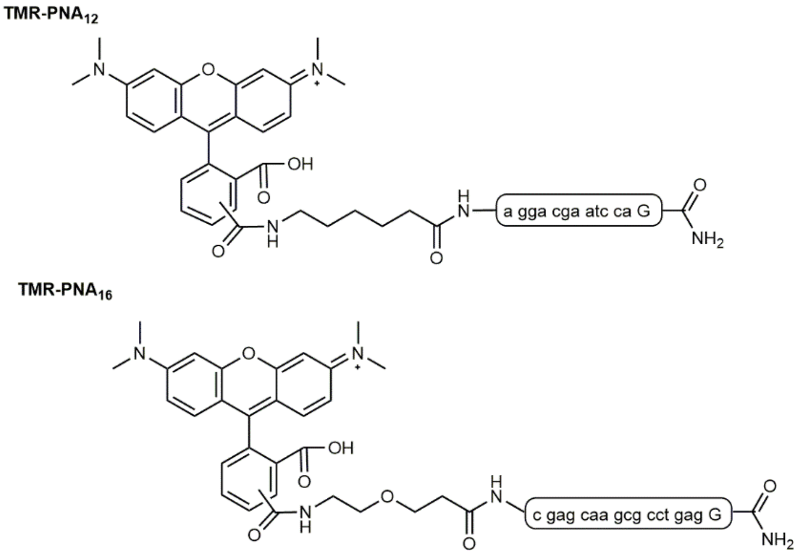

Figure 6-1 **Stuctures of 5,6 -Carboxytetramethylrhodamine (TMR) conjugated PNA used for hybridisation with PNA-tagged Cys-E3-EGFR-eGFP** in Figure 1 of the main article (TAMRA-$PNA_{12}$) and Supplementary Chapter 9.1.3 (TAMRA-$PNA_{16}$).

### 7 Cloning of Cys-E3 and Cys-P1 tagged proteins.

#### 7.1 Cloning of Cys-E3-EGFR-eGFP

A vector containing EGFR-eGFP was kindly donated by Dr. Donna Arndt-Jovin (Max-Planck-Institut für Biophysikalische Chemie, Göttingen). This vector was used for cloning of Cys-E3-EGFR-eGFP which was carried out by *GeneScript* (Piscataway, NJ, USA). The first 24 amino acids of EGFR code for a signal peptide (UniProt Q504U8), which is cleaved off during biosynthesis. The DNA sequence of the Cys-E3 peptide was therefore inserted after amino acid 24; between the C terminal end of the EGFR signal peptide and the N terminal end of the mature protein (Figure 7-1). After biosynthesis and cleavage of the signal peptide, the mature protein with N terminal Cys E3 -tag should be revealed.

GeneScript carried out cloning of the vector. Fot this, the EGFR-eGFP vector was restricted by NheI (GCTAGC (591 – 596)) and BglII (AGATCT (3519 – 3524)) and the tag sequence added by directed mutagenesis with suitable primers, followed by ligation into the opened vector. Restriction mapping with KpnI and BglII (*New England Biolabs,* Ipswitch, U.S.A) confirmed the insertion of the tag sequence (Figure 7-2).

EGFR Signal peptide (N → C): MRPSGTAGAA LLALLAALCP ASRA

DNA sequence of signal peptide (5' → 3'): ATG CGA CCC TCC GGG ACG GCC GGG GCA GCG CTC CTG GCG CTG CTG GCT GCG CTC TGC CCG GCG AGT CGG GCT

Cys-E3 Peptide (N → C): C E I A A L E K E I A A L E K E I A A L E K G G

Inserted DNA sequence of Cys-E3 Peptide (5' → 3'): TGC GAG ATC GCC GCC CTG GAG AAG GAG ATC GCC GCC CTG GAG AAG GAG ATC GCC GCC CTG GAG AAG GGC GGC

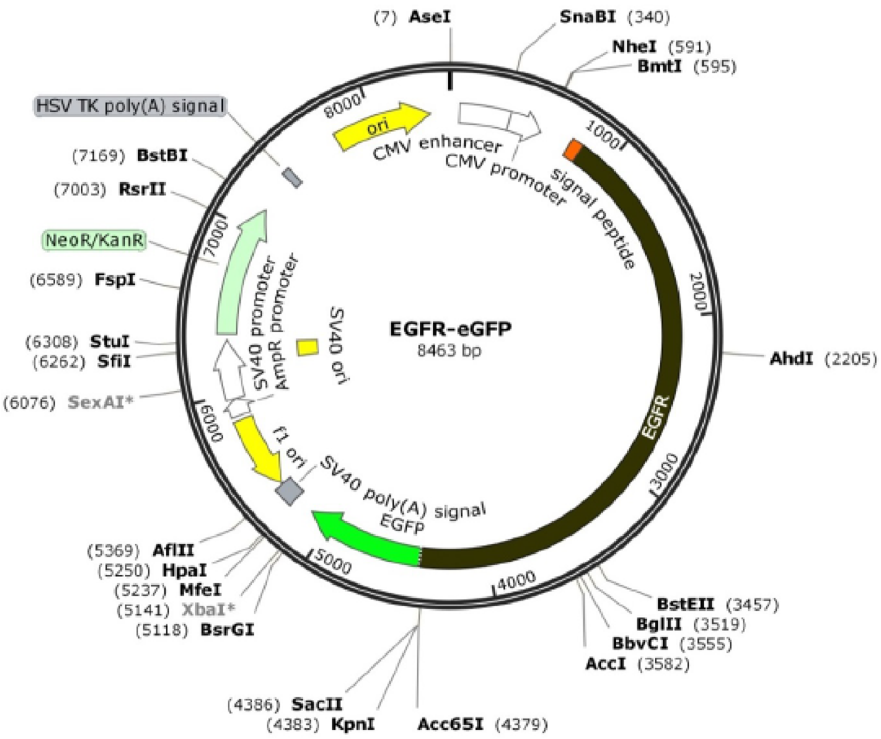

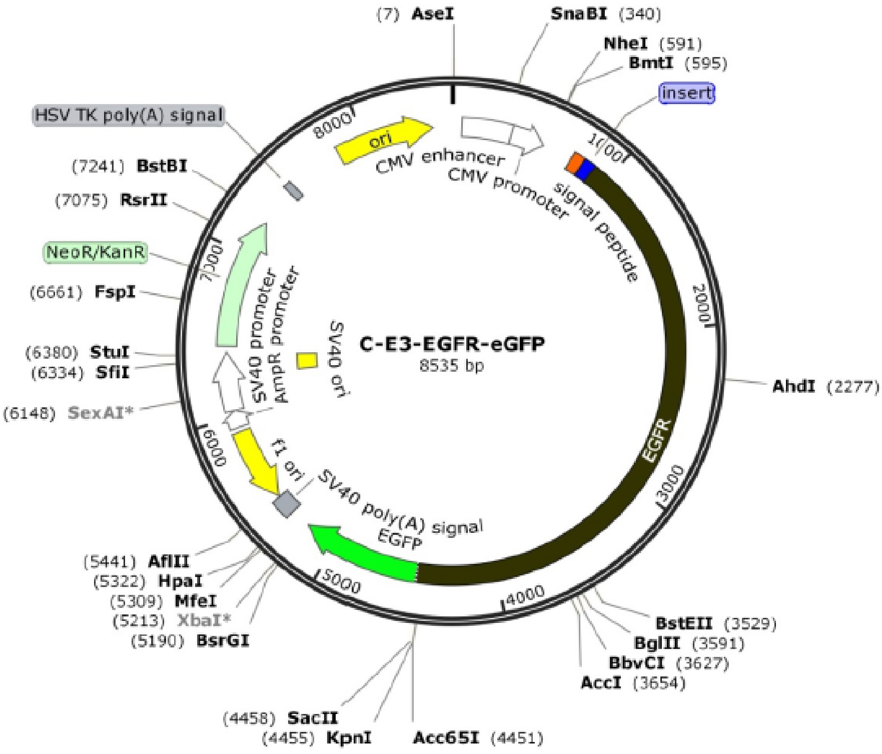

Figure 7-1 Vector maps for EGFR-eGFP kindly donated by Dr. Donna Arndt-Jovin (above) and C-E3-EGFR-eGFP (below) cloned by by *GeneScript*. Images made by SnapGene Viewer 3.0.3.

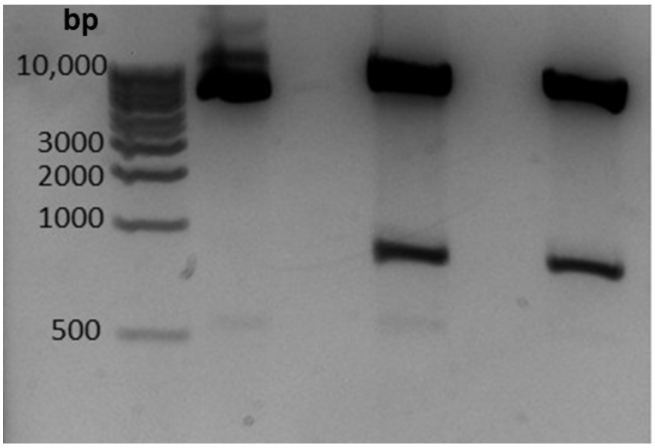

Figure 7-2 **Digestion of Cys-E3-EGFR-eGFP cloned by by *GeneScript* with restriction enzymes KpnI and BglII.** Lane1: KB ladder (*New England Biolabs*, U.S.A.) molecular weight shown in bp. Lane2: Cys-E3-EGFR-eGFP plasmid. Lanes 3 and 4: Kpn1, BglII digestion. Digestion was carried out once.

#### 7.2 Cloning of Cys-P1-EGFR-eYFP

##### 7.2.1 Cloning Cys-P1 into EGFR-eYFP-N1

A vector containing EGFR-eYFP-N1 was kindly donated by Prof. Thorsten Wohland (Centre for Bioimaging Sciences, National University of Singapore). The cloning strategy was similar as described in Chapter 7.1, with the Cys-P1 inserted after the signal peptide to ensure its location directly at the N terminus of EGFR after signal peptide cleavage. The gene for Cys-P1 was ordered from GeneScript (Piscataway, NJ, USA) and cloned by GeneScript into the plasmid EGFR-eYFP-N1 directly after the C terminal of the signal peptide (1323-1394, Figure 7-3)

EGFR Signal peptide: (N → C): MRPSGTAGAA LLALLAALCP ASRA

DNA sequence of EGFR signal peptide 1323-1394: (5' → 3'): ATG CGA CCC TCC GGG ACG GCC GGG GCA GCG CTC CTG GCG CTG CTG GCT GCG CTC TGC CCG GCG AGT CGG GCT

Cys-P1 Peptide (N → C): C EIQALEE ENAQLEQ ENAALEE EIAQLEY GG

Inserted DNA sequence of Cys-P1 Peptide (5' → 3'): TGC GAG ATC CAG GCC CTG GAG GAG GAG AAC GCC CAG CTG GAG CAG GAG AAC GCC GCC CTG GA GGA GGA GAT CGC CCA GCT GGA GTA CGG CGG C

##### 7.2.2 Cloning Cys-P1-EGFR-eYFP into the PiggyBac plasmid.

Cys-P1-EGFR-eYFP was inserted into the PiggyBac plasmid pPBtet-3xFLAG-IRES-DsRed-Express-PuroR donor vector[5] by GenScript (Piscataway, NJ, USA). Cys-P1-EGFR-eYFP was amplified with the addition of two unique SfiI restriction sites directly before and after the gene, before cloning into the pPBtet-3xFLAG-IRES-DsRed-Express-PuroR donor vector using the SfiI restriction endonuclease to give the vector pPBtet-Cys-P1-EGFR-eYFP-PuroR (Figure 7-4) used for generation of the stable cell line.

DNA sequence inserted before ATG start codon of Cys-P1-EGFR-eYFP:  GGCCTCTGAGGCC

DNA sequence inserted before TAA stop codon of Cys-P1-EGFR-eYFP: GGCCTGTCAGGCC

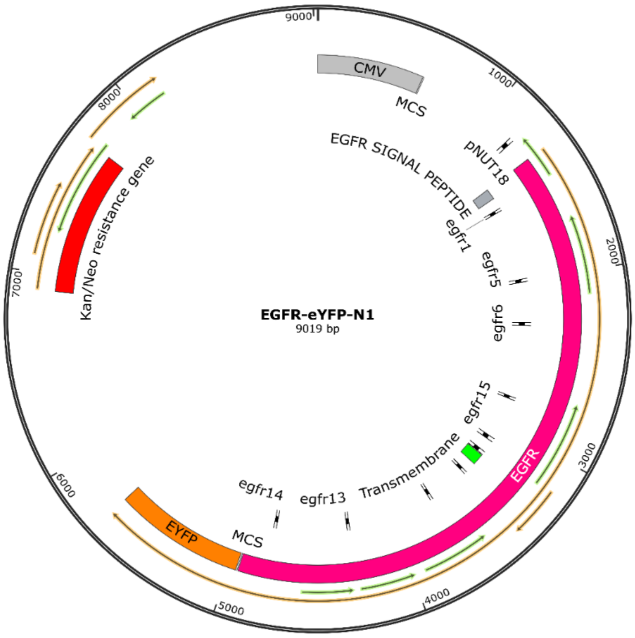

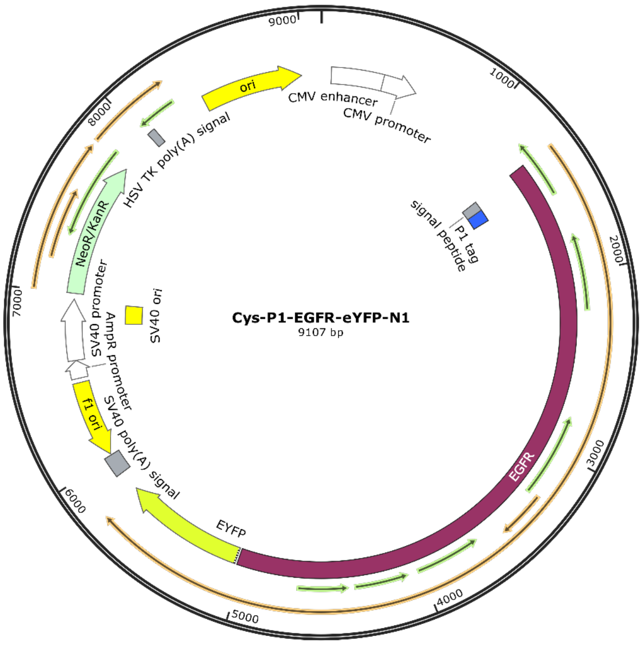

Figure 7-3 Plasmid map of EGFR-eYFP-N1 kindly donated by Prof. Thorsten Wohland (above) and plasmid map of Cys-P1-EGFR-eYFP-N1 cloned by *GeneScript* (Piscataway, NJ, USA). (below). Cys-P1-peptide was cloned directly before the signal peptide

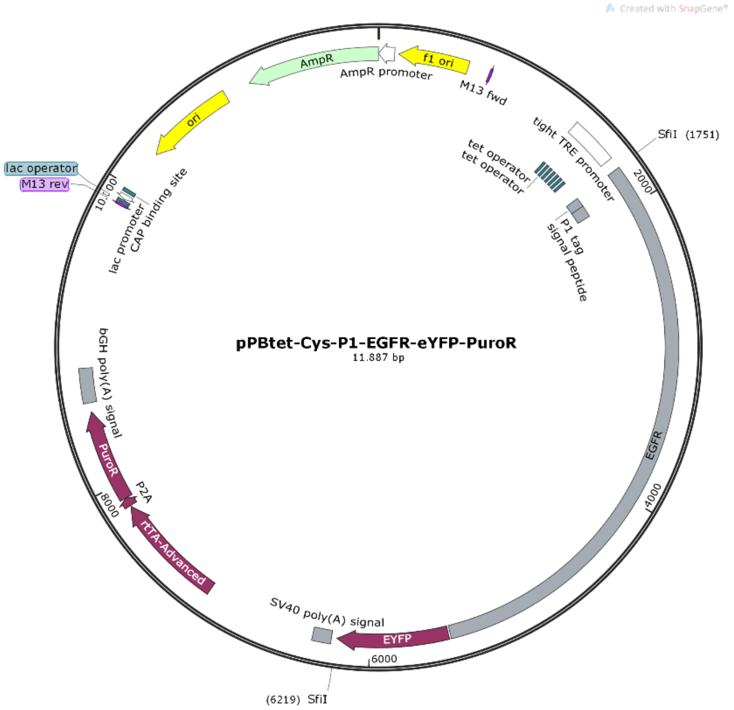

Figure 7-4 Plasmid map of pPBtet-Cys-P1-EGFR-eYFP-PuroR donor vector[5] used for PiggyBac transposition, cloned by *GeneScript* (Piscataway, NJ, USA).

#### 7.3 Cloning of Cys-P1-$ET_B$R-GFPspark

*Reagents*: The plasmid pCMV3-EDNRB-GFPspark was purchased from Sino Biological Inc. (Peking, China). Modifying and restriction enzymes were purchased from Thermo Fischer Scientific (Waltham, MA, USA). PureYield™ Plasmid Miniprep System and Wizard® SV Gel and PCR Clean-Up System were obtained from Promega (Madison, WI, USA). All oligonucleotides were purchased from Biomers (Ulm, Germany). Hygromycine B Gold (100 µg/mL) was purchased from InvivoGen (San Diego, CA, USA) and kanamycine sulfate was acquired from Carl Roth GmbH + Co. KG (Karlsruhe, Germany).

*Cloning:* The plasmid encoding the human endothelin B receptor ($ET_B$R) carrying a C-terminal GFP (pCMV3-EDNRB-GFPspark) was obtained from Sino Biological Inc. (Peking, China). To allow expression of the GPCR construct, the Cys-P1-tag was introduced after the endogenous signalpeptide of the endothelin B receptor (amino acid 1-26 of the encoded protein). The Cys-P1 tag was introduced by site-directed mutagenesis based on the restriction-free ligation protocol[6]. The following primers were used for introducing the Cys-P1 tag into the cDNA of the receptor:

Forward primer for Cys-P1-$ET_B$R-GFPspark generation, containing the C-terminal part of the signal peptide and the N-terminal part of the Cys-P1 tag:

5’-CGGCCTGTCGCGGATCTGGGGATGCGAGATCCAGGCCCTG-3’

Reverse primer for Cys-P1-$ET_B$R-GFPspark generation, containing the C-terminal part of the Cys-P1 tag, a Gly-Gly-Ser linker and the N-terminal part of the receptor:

5’-CAGGCGGGAAGCCTCTCTCCTCTGAGCCGCCGTACTCCAG-3’

Mutagenesis was carried out in two separate polymerase reactions. First PCR was used to generate the mutagenesis primer, using the overlapping forward and reverse primer. The newly synthesized primer was purified by gel extraction (Wizard® SV Gel and PCR Clean-Up System). Next, the obtained mutagenesis primer was used in a second PCR with the pCMV3-EDNRB-GFPspark plasmid to insert the Cys-P1 tag in frame. Clone selection was carried out in *E. coli* DH5α and positive clones were isolated using the PureYield™ Plasmid Miniprep System and sequenced (sequencing unit Leipzig University).

##### 7.4 Cloning of Cys-E3-hY$_2$R

The plasmids for the Cys-E3-hY$_2$R-eYFP and Cys-E3-hY$_2$R construct were previously described [7,8]

### 8 Cell culture

#### 8.1 Reagents and media for cell culture, fluorescence microscopy and cellular assays

##### 8.1.1 CHO and HEK293 Flp-In™ T-REx™ cells

Dulbecco’s Modified Eagle’s Medium (DMEM), Nutrient Mixture F-12 Ham **(**N6658), Fetal Bovine Serum (FBS) superior, L-Glutamine, penicillin, streptomycin, doxycycline, trypsin, TRIS and EDTA were purchased from *Merck* (Darmstadt, Germany).  Trypsin / EDTA (0.25%) , OptiMEM, Phosphate buffered saline (PBS, 100 mM phosphate, 150 mM sodium chloride), Dulbecco’s phosphate buffered saline (DPBS), HEPES  buffer, Hanks balanced salt solution (HBSS, calcium, magnesium, no phenol red), Opti-MEM, Puromycin Dihydrochloride, Lipofectamine®2000, Lipofectamine®3000, Poly-D-Lysine (0.1 mg/mL), Hoechst 33342 0.5mg/mL, Normal goat serum, Pierce™ 16% Formaldehyde (w/v) Methanol-free, ProLong™ Live Antifade, were purchased from *Thermo Fischer Scientific* (Waltham, U.S.A.) . Triton™ X-100, Salmon Sperm DNA sodium salt, BSA (Albumin Fraction V for use in blocking buffer) was purchased from *Carl Roth* (Karlsruhe, Germany). 8-well µ-slides were acquired from *ibidi GmbH* (Martinsried, Germany). Fluorescence labeled and non-labeled oligonucleotides were purchased from *Biomers* (Ulm, Germany).

##### 8.1.2 HEK293, HEK293-Cys-E3-hY$_2$R and COS-7 cells

Dulbecco’s modified Eagle’s medium (DMEM, containing 4.5 g/L glucose), Ham’s F12 (containing L-glutamine), Dulbecco’s phosphate buffered saline (DPBS), Hank’s balanced salt solution (HBSS), trypsin / EDTA, and OptiMEM were purchased from *Lonza Group Ltd.* (Basel, Switzerland). FBS superior was obtained from *Biochrom GmbH* (Berlin, Germany). Metafecene® Pro was obtained from *Biontex Laboratories GmbH* (Munich, Germany). Hygromycin B Gold was purchased from InvivoGen (San Diego, CA, USA). Bovine serum albumin (BSA) was acquired from *Sigma Aldrich* (Taufkirchen, Germany). Endothelin 1 was purchased from Bachem (Bubendorf, Switzerland). Lipofectamine® 2000 was from *Invitrogen* (Carlsbad, CA, USA). Poly-D-lysine hydrobromide was acquired from *Sigma Aldrich* (Taufkirchen, Germany). 8-well µ-slides were acquired from *ibidi GmbH* (Martinsried, Germany). Hoechst 33342 (20 mM) was purchased from *Thermo Fischer Scientific* (Waltham, U.S.A.) Fluorescence labeled and non-labeled oligonucleotides were purchased from *Biomers* (Ulm, Germany).

#### 8.2 HEK293 cell culture

HEK293 (human embryonic kidney cells) cells were grown as monolayers at 37 °C, 5 % $CO_2$, and 95% humidity if not stated otherwise.

##### 8.2.1 HEK293 Flp-In™ T-REx™ cell culture

HEK293 (Flp-In™ T-REx™ 293 Cell Line, R78007, *Thermo Fischer Scientific*) were cultured in Dulbecco's Modified Eagle's Medium (DMEM) containing 10% FBS and penicillin / streptomycin (10,000 units/ mL). After reaching around 95 % confluence, cells were washed twice with DPBS and detached by incubation with trypsin / EDTA at 37 °C for roughly 2 min at 37 °C. Cells were resuspended in cell culture medium and reseeded as needed.

##### 8.2.2 HEK293 cell culture

HEK293 cells were cultured in DMEM/Ham's F12 (1:1 v/v), supplied with 15% (v/v) heat-inactivated FBS. After reaching around 95 % confluence, cells were washed twice with DPBS and detached by incubation with trypsin / EDTA at 37 °C. Cells were resuspended in cell culture medium and reseeded as needed.

##### 8.2.3 HEK293-Cys-E3-h$Y_2$R cell culture

HEK293 Cys-E3-h$Y_2$R cells were cultured in DMEM/Ham's F12 (1:1 v/v), supplied with 15% (v/v) heat-inactivated FCS and 200 µg/mL Hygromycin B. After reaching confluence, cells were washed twice with PBS and detached by incubation with trypsin / EDTA at 37 °C. Cells were resuspended in cell culture medium and reseeded as needed.

##### 8.2.4 Generation of HEK293 with stable Cys-E3-h$Y_2$R expression

The generation of the HEK293 Cys-E3-h$Y_2$R cell line was previously reported.[7,8]

#### 8.3 CHO-Cys-P1-EGFR-eYFP cell culture

Chinese Hamster Embryo (CHO) cells were grown as monolayers at 37 °C, 5 % $CO_2$, and 95% humidity if not stated otherwise. Cells were cultured in Hams F12 from *Thermo Fischer Scientific* with additional 2mM L-Glutamine, and penicillin / streptomycin (10,000 units/ mL), puromycin (8 µg/mL) and 10 % FBS at 37 °C and 5% $CO_2$. For plating cells were washed with PBS and detached with 1 mL 0.25% trypsin / 0.02% EDTA solution for 2 min at 37 °C. Cells were centrifuged and medium was exchanged to fresh medium.

##### 8.3.1 Generation of a doxycycline inducible Cys-P1-EGFR-eYFP stable CHO cell line by PiggyBac transposition

To generate a stable cell line carrying doxycycline inducible Cys-P1-EGFR-eYFP, wild-type CHO cells (CHO Cell Line from Chinese hamster ovary, 85050302*, Merck)* were transfected with a 3:1 ratio of donor plasmid pPBtet-Cys-P1-EGFR-eYFP-PuroR and PiggyBac transposase (System Biosciences, #PB200PA-1) vector using Lipofectamine 3000 (Thermo Fisher Scientific) according to the manufacturer's instructions. Two days after transfection, cells were transferred to puromycin selection (8 µg/mL) for 13 days before fluorescence-activated cell sorting (FACS). To generate the Cys-P1-EGFR-eYFP stable CHO clone 12.3, transfected CHO cells were subjected to two rounds of FACS: First, eYFP negative uninduced cells were sorted to ensure tight expression of Cys-P1-EGFR-eYFP only upon doxycycline induction. Second, cells were treated for 18 h with doxycycline (100 ng/mL), 10,000 cells sorted based on thresholded levels of Cys-P1-EGFR-eYFP expression and plated at clonal density into a p100 cell culture dish without doxycycline. After 5 days, single colonies were picked, transferred to a 96-well plate, and expanded. Before starting experiments, the Cys.P1-EGFR-eYFP stable CHO clone 12.3 was analyzed by flow-cytometry for homogeneous levels of Cys-P1-EGFR-eYFP after doxycycline induction (100 ng/mL) for 18 h (Figure 8-1). To maintain expression of the pPBtet-Cys-P1-EGFR-eYFP-PuroR cassette, stable CHO cells were continuously cultured with puromycin (8 µg/mL).

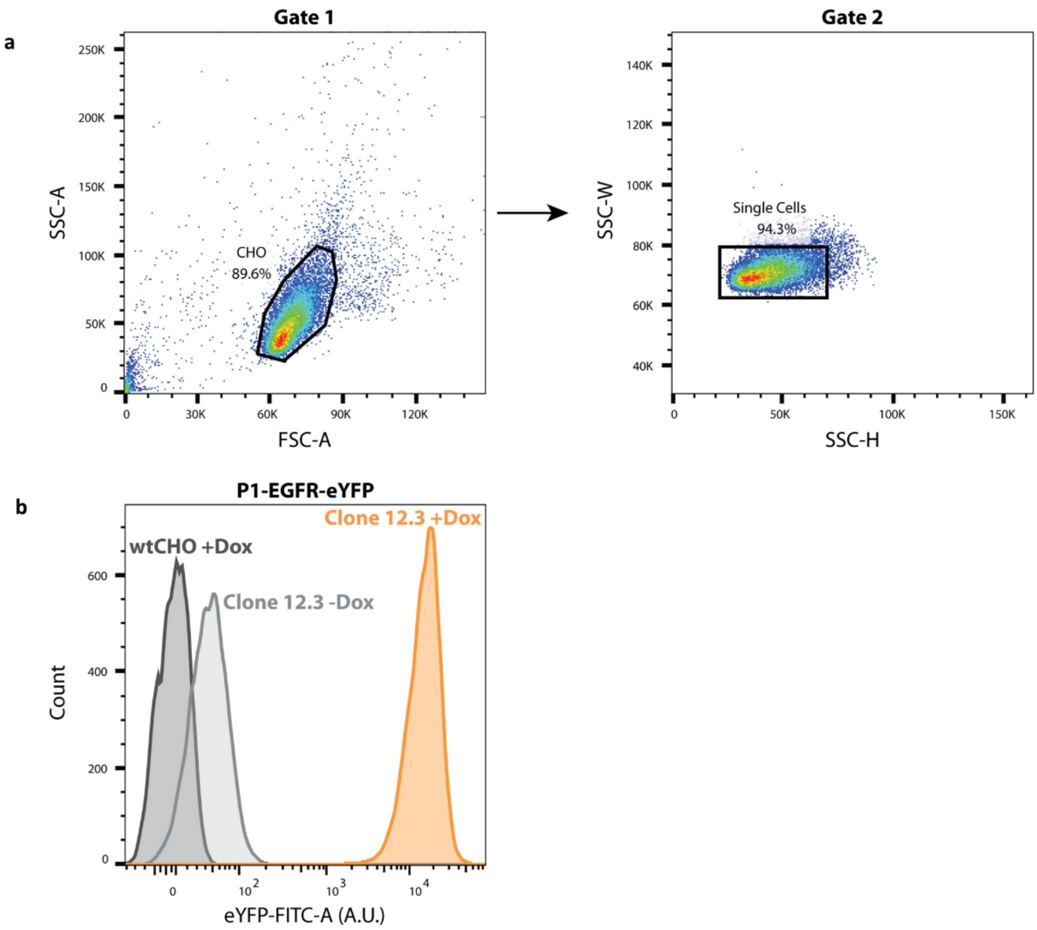

Figure 8-1 **Generation of doxycycline inducible, stable Cys-P1-EGFR-eYFP CHO cell line. a)** Exemplified gating strategy displaying wild-type CHO (wtCHO). Debris (FSC-A, SSC-A; Gate 1) and doublets (SSC-H, SSC-W; Gate 2) were excluded and 18,000 cells per sample analyzed for eYFP fluorescence intensity. **b)** Expression of P1-EGFR-eYFP was analyzed by fluorescence flow cytometry in the stable CHO clone 12.3 in the absence of doxycycline (-Dox) and 18 h after induction with 100 ng/mL doxycycline (+Dox). wtCHO +Dox are plotted as negative control. Fluorescence intensity of eYFP is indicated in arbitrary units (A.U.).

#### 8.4 COS-7 cell culture

COS-7 cells (African green monkey) were grown as monolayers at 37 °C, 5 % $CO_2$, and 95% humidity if not stated otherwise. COS-7 cells were cultured in DMEM supplied with 10 % (v/v) heat-inactivated FCS in cell culture flasks until reaching confluence. Then, cells were washed twice with PBS and detached by incubation with trypsin / EDTA at 37 °C. Cells were resuspended in cell culture medium and reseeded as needed.

### 9 Fluorescence microscopic analysis

Widefield fluorescence microscopy to investigate EGFR was performed using an IX83 microscope from *Olympus*. Transfected cells were imaged in four different channels (Hoechst33342: $\lambda ex$ = 350 ± 50 nm $\lambda em$ = 460 ± 50 nm; YFP: $\lambda ex$ = 500 ± 24 nm $\lambda em$ > 520 nm; TRITC: $\lambda ex$ = 575 ± 25 nm $\lambda em$ > 593 nm CY7). For HEK293 cells, 0.27 µm For experiments with Cys-E3-EGFR-eGFP, z stacks were imaged and the images were processed by Wiener deconvolution. Cell Sens Dimension V1.17 *(Olympus)* software was used for image manipulation. ImageJ software was used to generate line intensity profiles.

Inverted Scanning Disc Confocal Laser microscopy to investigate EGFR was carried out on a Visitron VisiScope with an *Olympus* IX83 microscope and a *Yokogawa* CSU-W1 spinning disk unit and detection with an *Andor* iXon Ultra 888 EM-CCD. All pictures were collected as Z stacks through at cell using *VisiView®* Software. Images shown are of a single layer. Diode lasers: Hoechst 33342) 405 nm; YFP) 488 nm; Atto565) 561 nm; Atto647N) 640 nm. Dichroic emission filters Hoechst 33342) $\lambda_{em}$ = 460 ± 50 nm; YFP) $\lambda_{em}$ 470 ± 24 nm; Atto565) $\lambda_{em}$ 600 ± 50 nm. Atto647N) $\lambda_{em}$ = 700 ± 75 nm.

Fluorescence images to investigate the GPCRs $hY_2R$ and $ET_BR$ were acquired by using a *Zeiss* Axio Observer.Z1 microscope with an ApoTome.2 Imaging System, a C-Apochromat 63x/1.20 W Corr M27 objective, an AxioCamMRm camera and the ZEN 2.0 software. Transfected cells were imaged using different Zeiss filter sets: Atto565 ($\lambda_{ex}$: 565/30; beamsplitter: 585; emission: $\lambda_{em}$ 620/60).

#### 9.1 Labelling of Cys-E3-EGFR-eGFP on HEK293 Flp-In™ T-REx™ cells

##### 9.1.1 Coiled coil templated TMR labelling of Cys-E3-EGFR-eGFP on HEK293 Flp-In™ T-REx™ cells.

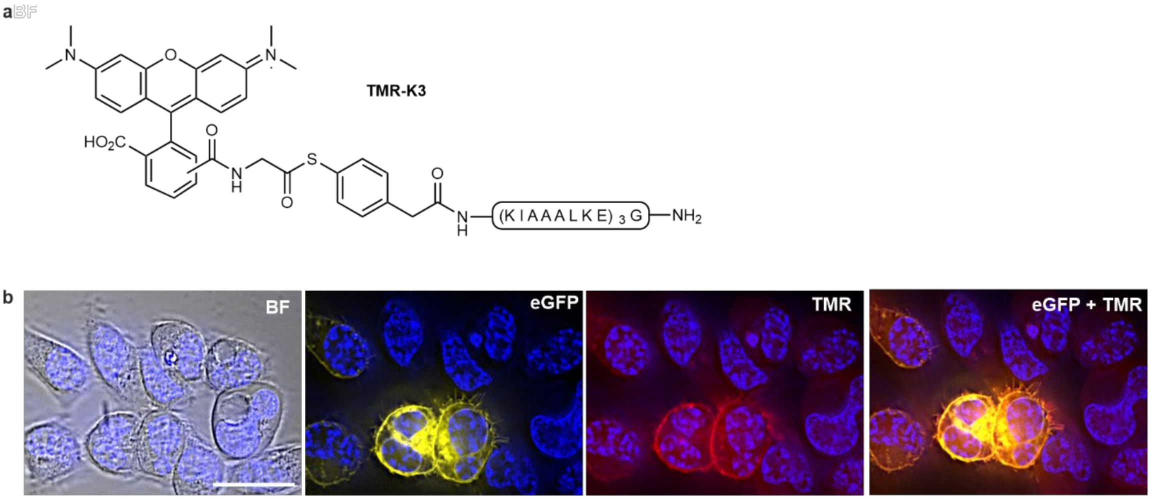

Figure 9-1 **Coiled coil templated fluorescent labelling of Cys-E3-EGFR-eGFP with 5,6 - Carboxytetramethylrhodamine (TMR). a)** Structure of TMR-K3 hybridization probe. **b)** Fluorescence microscope characterization labelling of Cys-E3-EGFR-eGFP on HEK293 cells. After staining of nuclei with Hoechst 33342, transiently transfected cells were treated with TCEP (0.1 mM) for 2 min and TMR-K3 (100 nM) for 4 min. Prior to imaging, cells were washed with basic buffer (200 mM NaHCO3 in PBS, pH 9.0, 2 min). All pictures show Hoechst 33342 stain (λex = 350 ± 50 nm, λem = 460 ± 50 nm, 10 ms excitation). From left to right; Bright field (117 ms), eGFP signal (λex = 500 ± 24 nm, λem > 520 nm, 1 s excitation), labelled hybridization probe signal (λex = 575 ± 25 nm, λem > 593 nm, 5 s excitation). Scale bar is 10 μM. Experiment was repeated three times independently with similar results.

##### 9.1.2 PNA tagging of Cys-E3-EGFR-eGFP on HEK293 Flp-In™ T-REx™ cells and fluorescence microscopic imaging with hybridization probes.

**a)** PNA tagging with **$PNA_{11}$-K3**, incubation with **TMR PNA-11mer**. Histogram intensity window: 2000-7000

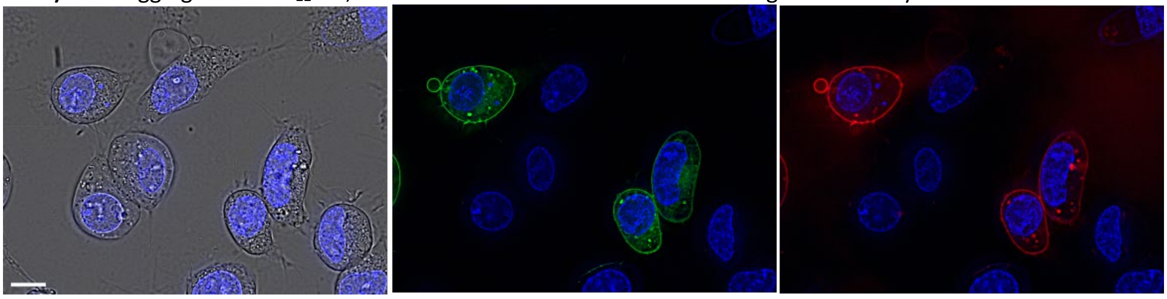

**b)** no PNA tagging, incubation with **TMR PNA-11mer** (control). Histogram intensity window: 2000-7000

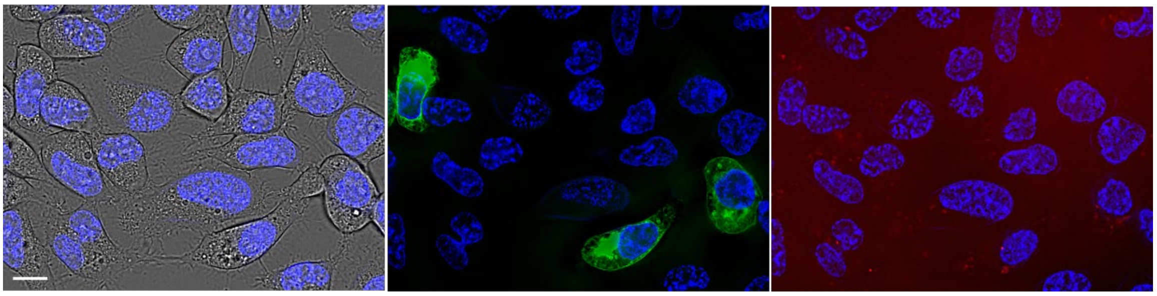

**c)** PNA tagging with **PNA$_{16}$-K3**, incubation with **Cy3 DNA-16mer**. Histogram intensity window: 4000-10000

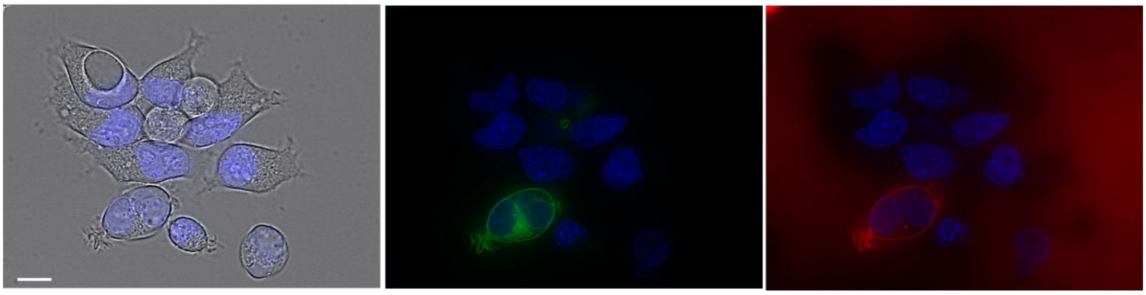

**d)** no PNA tagging, incubation with **Cy3 DNA-16mer** (control). Histogram intensity window: 4000-10000

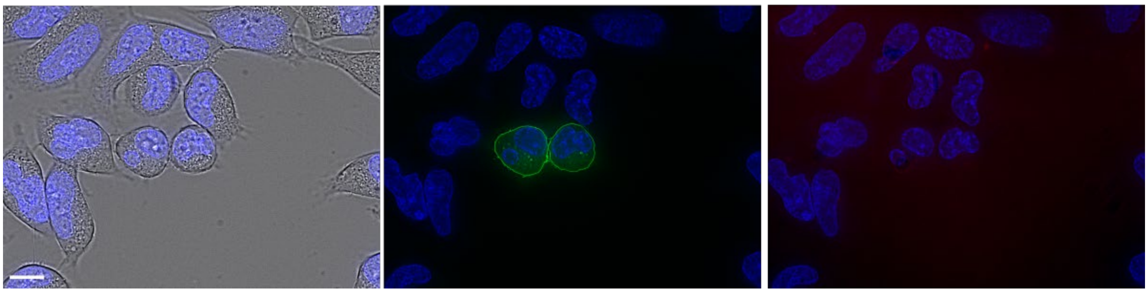

**f)** PNA tagging with **PNA$_{16}$-K3**, incubation with **Cy3 DNA-11mer**. Histogram intensity window: 2000-6000

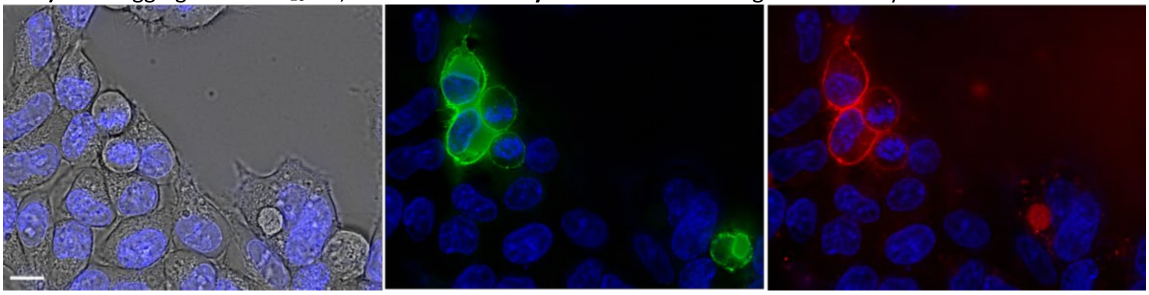

**g)** PNA tagging with **PNA$_{16}$-K3**, incubation with **ATTO565 DNA-16mer**. Histogram intensity window: 16000-26000

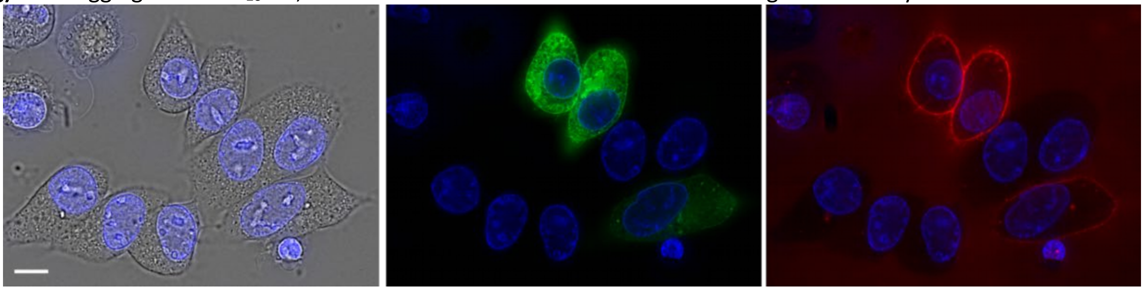

Figure 9-2 **Zoomed out fluorescence microscopy images used in Figure 2 in main article.** Transiently transfected Cys-E3-EGFR-eGFP HEK293 Flp-In™ T-REx™ cells were stained with Hoechst 33342 for 10 mins at 37℃ and treated for 2 min with TCEP (0.1 mM) in PBS then incubated with donor **PNA$_{16}$-K3** or **PNA$_{11}$-K3** (100 nM) in PBS (pH 7.0), 4 min. Cells were washed with PBS, incubated with fluorescently labelled DNA or PNA (100 nM) for 4 min before washing with PBS and imaging. From left to right: Bright field (117 ms), Hoechst 33342 stain ($\lambda_{ex}$ = 350 ± 50 nm, $\lambda_{em}$ = 460 ± 50 nm, 10 ms excitation), eGFP signal ($\lambda_{ex}$ = 500 ± 24 nm, $\lambda_{em}$ > 520 nm, 1 s excitation), labelled hybridization probe signal ($\lambda_{ex}$ = 575 ± 25 nm, $\lambda_{em}$ > 593 nm, 5 s excitation). Histogram intensity windows given for the TRITC channel. All other histogram settings were adjusted linearly. Scale bars are 10 µM. Experiment was repeated three times independently with similar results.

##### 9.1.3 Multilabelling of Cys-E3-EGFR-eGFP on HEK293 Flp-In™ T-REx™ cells with TMR-PNA constructs

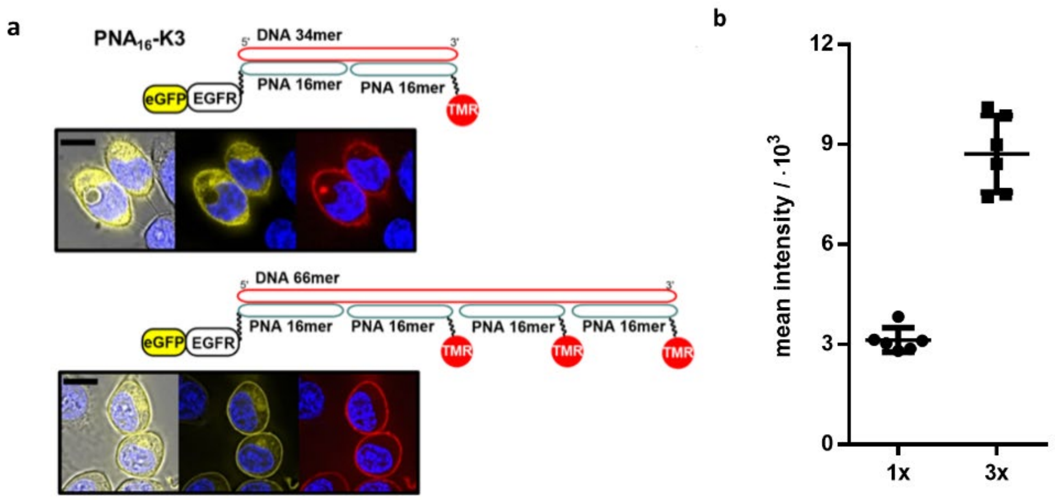

Figure 9-3 **Multilabelling of Cys-E3-EGFR-eGFP on HEK293 Flp-In™ T-REx™ cells with TMR-PNA constructs. a)** Fluorescence microscopic characterization of PNA-tag enabled fluorescence labelling of Cys-E3- EGFR-eGFP on HEK293 cells. After staining of nuclei with Hoechst 33342, transiently transfected cells were treated with **$PNA_{16}$-K3.** Subsequently, cells were incubated with **a DNA-34mer** (top) or **DNA-66mer** adapter (bottom) and a **TMR-labeled PNA-16mer**. Conditions for PNA transfer: i) 0.1 mM TCEP in PBS (pH 7.0), 2 min, 25°C; ii) 100 nM donor $PNA_{16}$-K3 in PBS buffer (pH 7.0), 4 min, 25°C. Conditions for hybridization: 1 µM DNA 34mer or 66mer and 3 µM TMR-PNA 16mer in PBS (pH 7.0). 5 min, 25°C. Scale bars are 10 µM. Brightness of 1x TMR picture was increased relative to the 3x TMR picture. **b)** Mean TMR/eGFP intensities of cell membrane regions labeled with either one or three TMR-DNA-probes are shown at the right. Values were collected from from three distinct experiments, two cells from each experiment (n=6). Plot is presented as the mean +/- SD.

#### 9.2 Labelling of Cys-P1-EGFR-eYFP on CHO cells

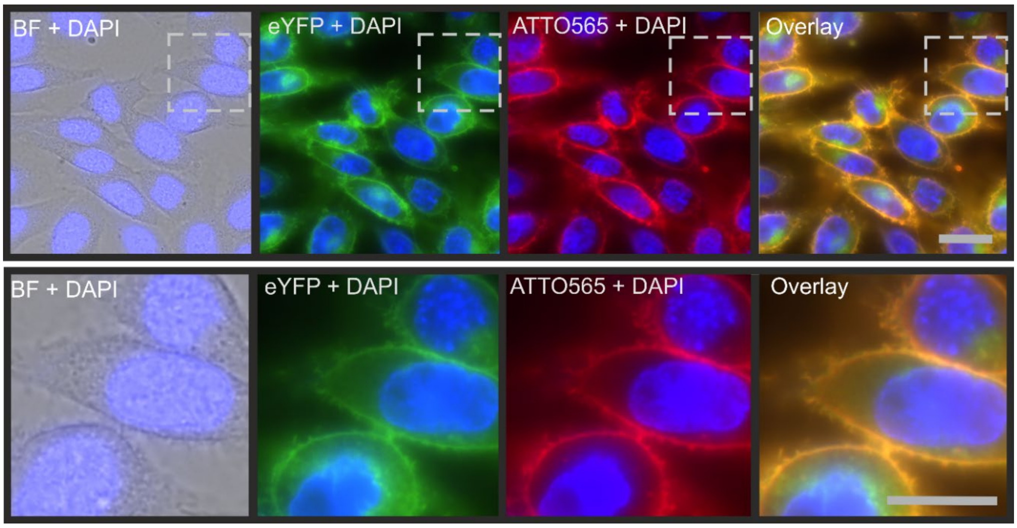

Figure 9-4 **Fluorescence microscopic characterization of PNA-tag enabled fluorescence labelling of stable Cys-P1-EGFR-eYFP expressing CHO cells**.After staining of nuclei with Hoechst 33342, doxycycline induced cells were treated with **PNA$_{15}$-P2** for 4 min in HBSS. Subsequently, cells were incubated with Atto565-labeled DNA-15mer. Conditions for PNA transfer: 100 nM donor **PNA$_{15}$-P2** in HBSS buffer, 4 min. Conditions for hybridization: 100 nM Atto565 DNA-15mer in HBSS, 4 min, 25°C. Excitation times: Hoechst 33342 15ms; YFP; 150 ms Atto565; 500 ms .Filter settings : Hoechst 33342) $\lambda_{ex}$ = 350 ± 50 nm, $\lambda_{em}$ = 460 ± 50 nm; CFP) $\lambda_{ex}$ = 438 ± 24 nm, $\lambda_{em}$ 483 ± 24 nm, YFP) $\lambda_{ex}$ = 500 ± 24 nm, $\lambda_{em}$ 545 ± 40 nm, Atto565) $\lambda_{ex}$ = 575 ± 25 nm, $\lambda_{em}$ 623 ± 40 nm. Scale bar= 20 µm. Experiment was repeated three times independently with similar results.

##### 9.2.1 Control experiments for labelling of CHO cells with PNA$_{15}$-P2

For control experiments, CHO cells stably expressing Cys-P1-EGFR-eYFP were seeded and prepared as described in the Methods section but without doxycycline addition. Labelling reactions were carried out with **PNA$_{15}$-P2** and Atto565 DNA-15mer as described in Fig. S 9.4.

Cys-P1-EGFR-eYFP (-) DOX

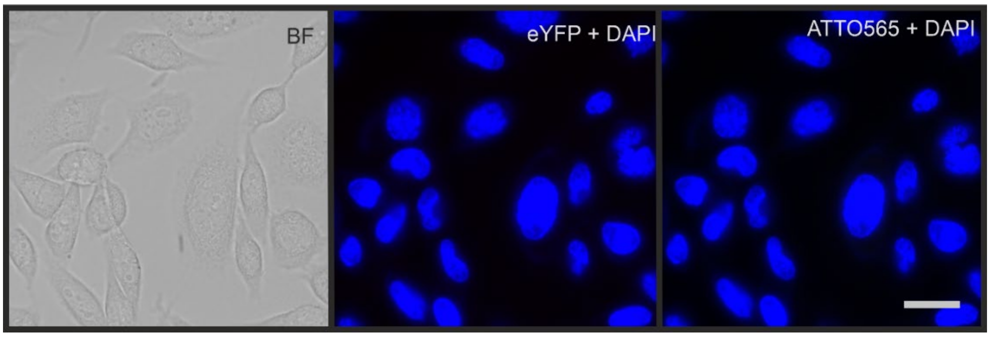

Figure 9-5 **Attempted labelling of CHO cells with PNA$_{15}$-P2 and Atto565-DNA 15mer upon treatment of Cys-P1-EGFR-eYFP CHO cells, without prior DOX induction**. No signal is seen in the Atto565 channel. Labelling procedure and fluorescence picture settings are identical to standard methods and that shown in Figure 9-4: After staining of nuclei with Hoechst 33342, Cells were treated with **PNA$_{15}$-P2** for 4 min in HBSS. Subsequently, cells were incubated with Atto565-labeled DNA-15mer. Conditions for PNA transfer: 100 nM donor **PNA$_{15}$-P2** in HBSS buffer, 4 min. Conditions for hybridization: 100 nM Atto565 DNA-15mer in HBSS, 4 min, 25°C. Excitation times: Hoechst 33342 15ms; YFP; 150 ms Atto565; 500 ms .Filter settings : Hoechst 33342) $\lambda_{ex}$ = 350 ± 50 nm, $\lambda_{em}$ = 460 ± 50 nm; CFP) $\lambda_{ex}$ = 438 ± 24 nm, $\lambda_{em}$ 483 ± 24 nm, YFP) $\lambda_{ex}$ = 500 ± 24 nm, $\lambda_{em}$ 545 ± 40 nm, Atto565) $\lambda_{ex}$ = 575 ± 25 nm, $\lambda_{em}$ 623 ± 40 nm. Scale bar= 20 µm. The experiment was repeated three times independently with similar results.

##### 9.2.2 Labelling of Cys-P1-EGFR-eYFP on CHO cells followed by staining with Cy7-DNA constructs for microscopy analysis

CHO cells stably expressing Cys-P1-EGFR-eYFP were seeded, prepared and labelled as described in the methods section with either **Complex I**; adaptor DNA-33mer with a single Cy7-15mer or **Complex II**; adaptor DNA-105mer with five Cy7-15mers. For Signal to Noise Ratio (SNR) calculations, cells (n=20) taken from three independent experiments were analyzed by taking a line intensity profile throughout the cell (Figure 9-6). For each cell, Cy7 or YFP signal was calculated as the max peak height at the membrane region, and the Noise calculated as the standard deviation of the signal from a background region. The background region was deemed as a nearby 'empty' region in the line intensity profile, with no cells, and where little to no YFP or Cy7 signal was observed. YFP signal was normalized to a 'Relative YFP signal' according to the equation

$$\text{Relative YFP signal} = \frac{\text{YFP Signal}}{\text{YFP(min)}}$$

Where YFP(*min*) is the minimum YFP signal (4841 RFU). To account for variation in peak signal between single cells due to different expression levels, Cy 7 Signal was corrected according to the equation,

$$\text{Corrected Cy7 signal} = \frac{\text{Cy7 Signal}}{\text{Relative YFP signal}}$$

Finally, SNR was calculated (Tables 9-1, 9-2):

$$\text{SNR} = \frac{\text{Corrected Cy7 Signal}}{\text{Cy7 Noise}}$$

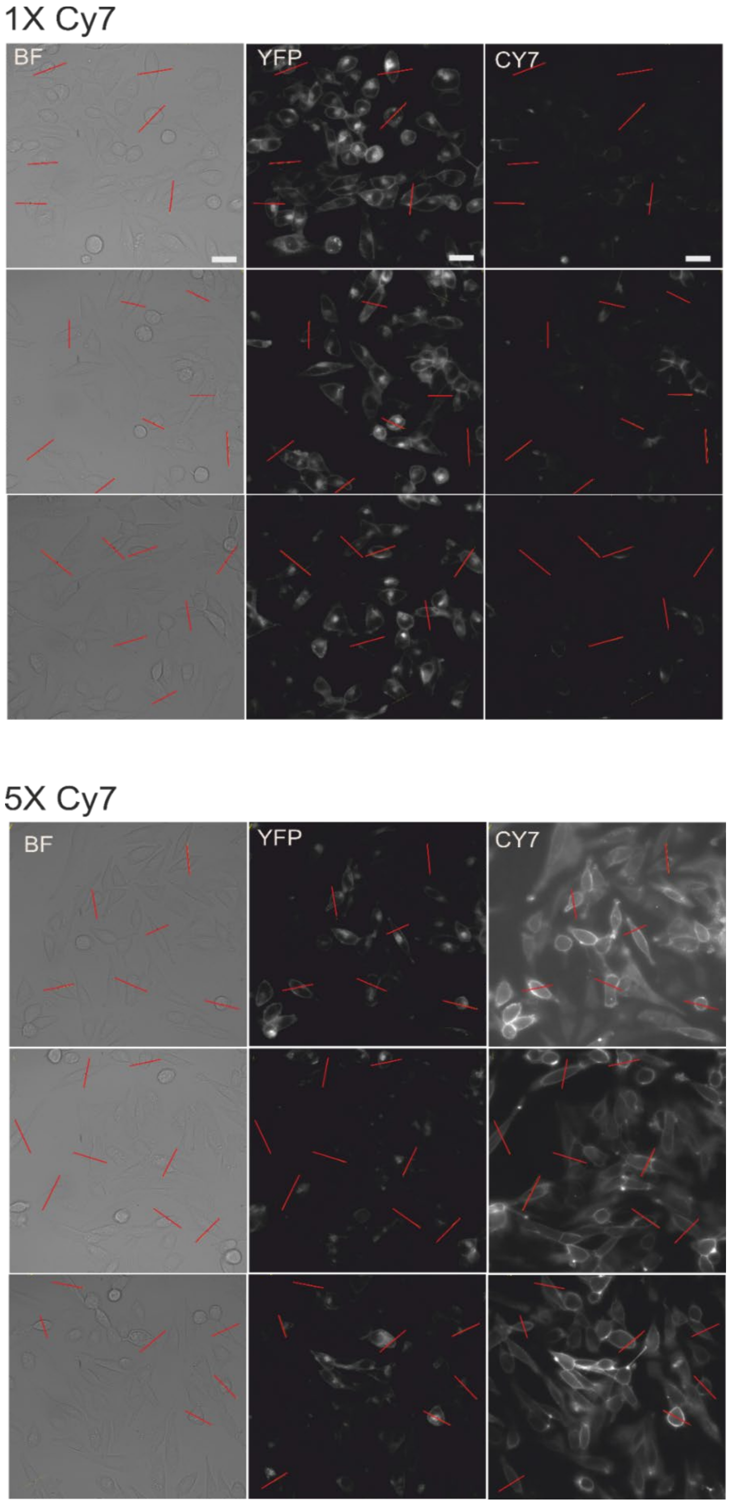

Figure 9-6 **Greyscale images used for calculation of Signal to Noise ratios depicted in Extended Data Figure 4.** Stably expressing Cys-P1-EGFR-eYFP cells were treated with with **PNA$_{15}$** for 4 minutes before washing and hybridization with 1xCy7- **Complex I** (adaptor DNA-33mer with a single Cy7-15mer; 1x Cy7) or **Complex II** (adaptor DNA-105mer with five Cy7-15mers; 5xCy7) in HBSS-BB. With the resultant images, ImageJ was used to generate line intensity profiles (shown as red lines). Excitation times: Cy7: 500 ms, YFP: 50 ms, Hoechst 33342:10 ms. Filter settings: Hoechst 33342) $\lambda_{ex}$ = 350 ± 50 nm, $\lambda_{em}$ = 460 ± 50 nm; YFP) $\lambda_{ex}$ = 500 ± 24 nm, $\lambda_{em}$ 545 ± 40 nm, Cy7) $\lambda_{ex}$ = 710 ± 75 nm, $\lambda_{em}$ 810 ± 90 nm. Scale bar= 20 µm. Image shows the n= 20 cells which were examined over 3 independent experiments, each experiment giving similar results.

Table 9-1 **Data used for calculation of Signal to Noise Rations (SNR) for 1x Cy7 labelling of PNA tagged Cys-P1-EGFR-eYFP**. Cells from three independent experiments were analyzed, with a line intensity profile taken throughout the cell (Figure 9-6). Signal was calculated as the max peak height at the membrane regions, and the Noise calculated as the standard deviation of the signal from a background region.

| | YFP | | | Cy7 1X | | | |
|---|---|---|---|---|---|---|---|
| **CELL** | **Signal (RFU)** | **Relative Signal** | **Noise (RFU)** | **Signal (RFU)** | **Corrected signal (RFU)** | **Noise (RFU)** | **Corrected Signal Cy7/ Noise Cy7 (RFU)** |
| **1.1** | 15623 | 3,23 | 55,0 | 8812 | 2730 | 194 | 14,1 |
| **1.2** | 12865 | 2,66 | 44,8 | 10238 | 3852 | 147 | 26,2 |
| **1.3** | 11863 | 2,45 | 35,0 | 6988 | 2851 | 134 | 21,2 |
| **1.4** | 16396 | 3,39 | 36,9 | 8447 | 2494 | 155 | 16,1 |
| **1.5** | 23603 | 4,88 | 59,6 | 9739 | 1997 | 158 | 12,7 |
| **1.6** | 9467 | 1,96 | 91,6 | 6639 | 3395 | 245 | 13,9 |
| **1.7** | 11094 | 2,29 | 18,4 | 6146 | 2682 | 138 | 19,5 |
| **2.1** | 15616 | 3,23 | 31,9 | 7788 | 2414 | 108 | 22,3 |
| **2.2** | 7131 | 1,47 | 67,4 | 4545 | 3085 | 324 | 9,5 |
| **2.3** | 15135 | 3,13 | 57,3 | 7344 | 2349 | 166 | 14,2 |
| **2.4** | 19554 | 4,04 | 68,4 | 6443 | 1595 | 147 | 10,8 |
| **2.5** | 19580 | 4,05 | 33,7 | 7547 | 1866 | 160 | 11,6 |
| **2.6** | 22881 | 4,73 | 54,5 | 11011 | 2329 | 184 | 12,7 |
| **3.1** | 13439 | 2,78 | 38,7 | 4481 | 1614 | 81 | 20,0 |
| **3.2** | 11274 | 2,33 | 36,8 | 7455 | 3201 | 91 | 35,4 |
| **3.3** | 16597 | 3,43 | 42,3 | 8741 | 2549 | 163 | 15,6 |
| **3.4** | 8193 | 1,69 | 36,8 | 5581 | 3297 | 128 | 25,7 |
| **3.5** | 12985 | 2,68 | 105,2 | 5343 | 1992 | 131 | 15,2 |
| **3.6** | 16054 | 3,32 | 70,0 | 7285 | 2196 | 135 | 16,3 |
| **3.7** | 16884 | 3,49 | 32,8 | 7504 | 2151 | 118 | 18,2 |
| **MEAN** | 14812 | 3,06 | 51 | 7404 | 2532 | 155 | 17,56 |
| **σ** | 4448 | 0,92 | 22 | 1764 | 610 | 54 | 6,28 |

Table 9-2 **Data used for calculation of Signal to Noise Rations (SNR) for 5x Cy7 labelling of PNA tagged Cys-P1-EGFR-eYFP**. Cells from three independent experiments were analyzed, with a line intensity profile taken throughout the cell (Figure 9-6). Signal was calculated as the max peak height at the membrane regions, and the Noise calculated as the standard deviation of the signal from a background region.

| | YFP | | | Cy7 5X | | | |
|---|---|---|---|---|---|---|---|
| CELL | Signal (RFU) | Relative Signal | Noise (RFU) | Signal (RFU) | Corrected signal | Noise | Corrected Signal Cy7/ Noise Cy7 |
| 1.1 | 8794 | 1,82 | 51,3 | 15254 | 8396 | 178 | 47,1 |
| 1.2 | 8563 | 1,77 | 100,8 | 20561 | 11623 | 209 | 55,6 |
| 1.3 | 20229 | 4,18 | 45,9 | 53918 | 12902 | 206 | 62,5 |
| 1.4 | 17820 | 3,68 | 47,1 | 28171 | 7652 | 200 | 38,3 |
| 1.5 | 7897 | 1,63 | 66,6 | 15547 | 9529 | 231 | 41,2 |
| 1.6 | 6555 | 1,35 | 16,9 | 18174 | 13420 | 226 | 59,5 |
| 1.7 | 8825 | 1,82 | 24,8 | 17370 | 9527 | 197 | 48,3 |
| 2.1 | 9099 | 1,88 | 21,9 | 21893 | 11646 | 200 | 58,3 |
| 2.2 | 6028 | 1,25 | 32,0 | 15731 | 12632 | 210 | 60,1 |
| 2.3 | 10702 | 2,21 | 50,6 | 31832 | 14398 | 201 | 71,8 |
| 2.4 | 6943 | 1,43 | 30,6 | 17063 | 11896 | 292 | 40,8 |
| 2.5 | 4841 | 1,00 | 20,3 | 15178 | 15178 | 216 | 70,4 |
| 2.6 | 10504 | 2,17 | 91,7 | 30142 | 13891 | 280 | 49,6 |
| 2.7 | 6905 | 1,43 | 14,1 | 17715 | 12418 | 168 | 73,9 |
| 2.8 | 6516 | 1,35 | 17,5 | 19977 | 14840 | 229 | 64,9 |
| 3.1 | 13451 | 2,78 | 30,0 | 38551 | 13873 | 282 | 49,1 |
| 3.2 | 13605 | 2,81 | 26,2 | 38438 | 13676 | 529 | 25,9 |
| 3.3 | 18689 | 3,86 | 50,6 | 49648 | 12859 | 332 | 38,7 |
| 3.4 | 7430 | 1,54 | 31,5 | 16566 | 10792 | 291 | 37,1 |
| 3.5 | 12470 | 2,58 | 50,3 | 37622 | 14604 | 449 | 32,5 |
| 3.6 | 12224 | 2,53 | 44,3 | 37034 | 14665 | 398 | 36,9 |
| MEAN | 10385 | 2,15 | 41 | 26494 | 12401 | 263 | 50,59 |
| σ | 4341 | 0,90 | 23 | 11962 | 2172 | 94 | 13,77 |

##### 9.2.3 EGF stimulated internalisation of Cys-P1-EGFR-eYFP on CHO cells with membrane label removal: Analysis by widefield microscopy.

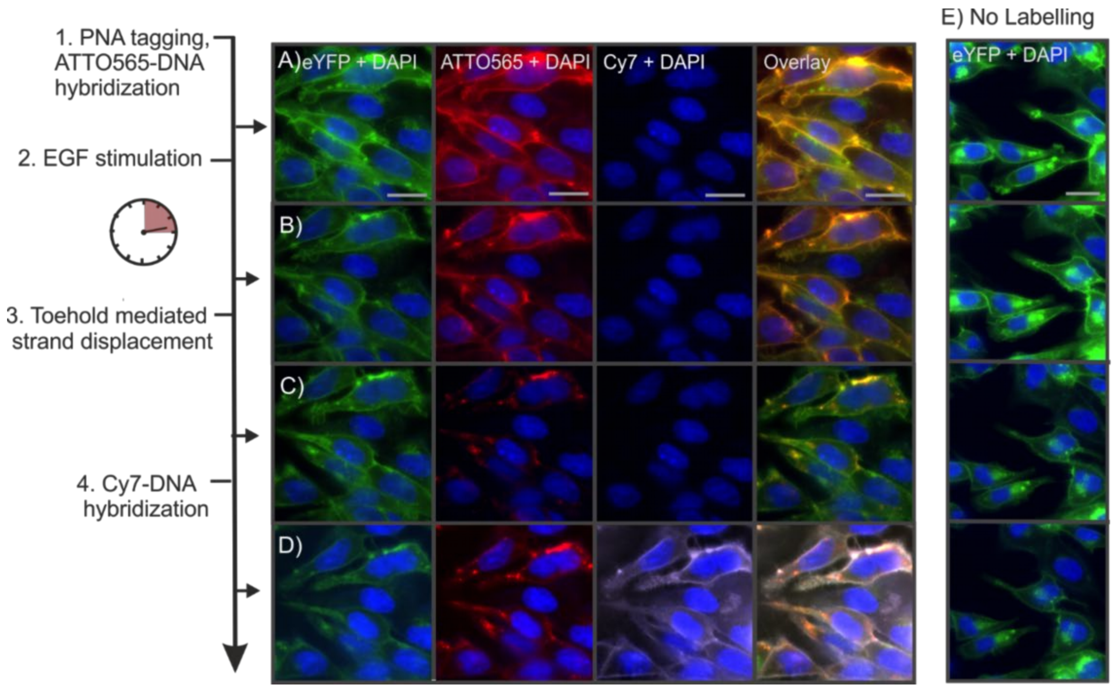

Figure 9-7 **Widefield fluorescence microscope characterization of PNA-tag enabled reversible fluorescence labelling of Cys-P1-EGFR-eYFP on serum starved CHO cells, stimulated with EGF**. After staining of nuclei with Hoechst 33342 in HBSS-BB, cells were treated with **PNA$_{15}$-P2** in HBSS, 4 min. **A)** Cells were incubated with 50 nM **Complex III**; adaptor DNA-105mer with five Atto565-DNA-23mers; in HBSS-BB for 4 min. **B)** 100 nM EGF stimulation for 15 mins **C)** Toehold mediated strand displacement of Atto565-23mer DNA with 300 nM displacement DNA-23mer and 100 nM EGF for 2 x 5 min in HBSS at 30°C. **D)** 100 nM Cy7-DNA-15mer, 3 min. **E)** The same experiment was carried out without addition of PNA or DNA.BB: Blocking Buffer: 0.1mg/mL salmon sperm DNA, 0.2% BSA, 1x ProLong™ Live Antifade in HBSS. Excitation times: Cy7: 500 ms, ATTO565: 50 ms YFP: 50 ms, Hoechst 33342:10 ms. Filter settings: Hoechst 33342) $\lambda_{ex}$ = 350 ± 50 nm, $\lambda_{em}$ = 460 ± 50 nm; YFP) $\lambda_{ex}$ = 500 ± 24 nm, $\lambda_{em}$ 545 ± 40 nm, Cy7) $\lambda_{ex}$ = 710 ± 75 nm, $\lambda_{em}$ 810 ± 90 nm. Scale bar= 20 µm. Experiments were repeated 3 times independently with similar results.

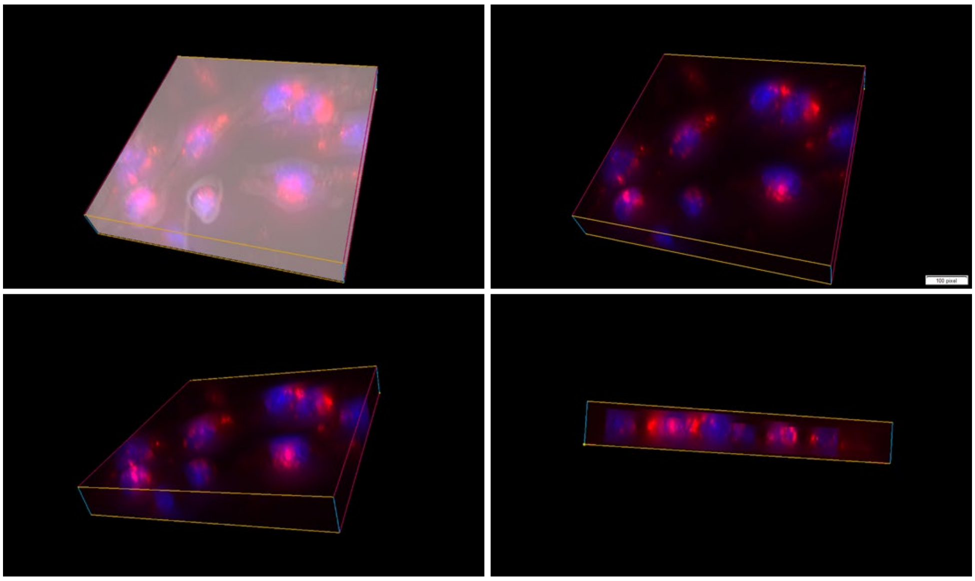

Figure 9-8 **Widefield fluorescence three-dimensional images of Cys-P1-EGFR-eYFP CHO cells reversibly labelled and stimulated with EGF.** 3D view of Figure9-6C. After staining of nuclei with Hoechst 33342, starved cells were treated with (i) **PNA$_{15}$-P2** in HBSS, 4 min (ii) 50 nM **Complex III**; adaptor DNA-105mer with five Atto565-DNA 23mers; in HBSS-BB for 4 min. (iii) 100 nM EGF was added, at 30°C 15 mins (iv) 300 nM displacement DNA-23mer and 100 nM EGF for 2 x 5 min in HBSS at 30°C. BB: Blocking Buffer: 0.1mg/mL salmon sperm DNA, 0.2% BSA, 1x ProLong™ Live Antifade in HBSS. 40 Z stacks were taken with an height of 0,36 µm. Excitation times: ATTO565: 50 ms YFP: 50 ms, Hoechst 33342:10 ms. Filter settings: Hoechst 33342) $\lambda_{ex}$ = 350 ± 50 nm, $\lambda_{em}$ = 460 ± 50 nm; YFP) $\lambda_{ex}$ = 500 ± 24 nm, $\lambda_{em}$ = 545 ± 40 nm. Scale bar= 20 µm. Top left image shows brightfield, Hoechst 33342 and Atto565 channels. The remaining images show Hoechst 33342 and ATTO565 channels from different angles. Experiments were repeated 3 times independently with similar results

##### 9.2.4 EGF stimulated internalisation of Cys-P1-EGFR-eYFP on CHO cells with membrane label removal; analysis by confocal microscopy

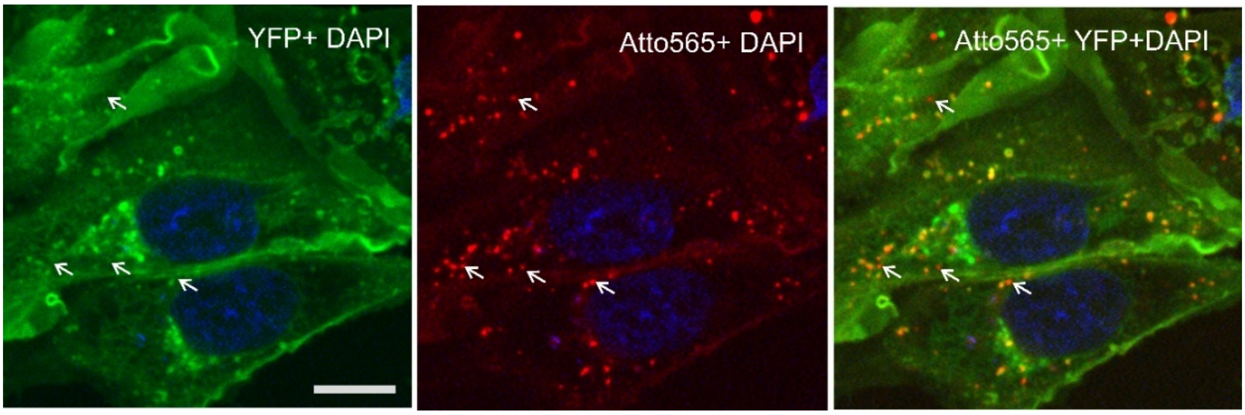

Figure 9-9 **Zoomed confocal microscopy picture of Cys-P1-EGFR-eYFP CHO cells shown in Figure 5f) of main article.** After staining of nuclei with Hoechst 33342, starved cells were treated with **PNA$_{15}$-P2** in HBSS, 4 min before addition of 50 nM **Complex III** (adaptor DNA-105mer + five Atto565-DNA-23mers) in HBSS-BB for 4 min then stimulation with EGF (100 nM) for 15 mins. Toehold mediated strand displacement of Atto565-23mer DNA with 300 nM displacement DNA-23mer in presence of 100 nM EGF (2 x 5 min in HBSS at 30°C). Excitation time:, ATTO565: 200 ms YFP: 100 ms, Hoechst 33342:100 ms Diode lasers: Hoechst 33342) 405 nm; YFP) 488 nm; Atto565) 561 nm; Dichroic emission filters: Hoechst 33342) $\lambda_{em}$ = 460 ± 50 nm; YFP) $\lambda_{em}$ 470 ± 24 nm; Atto565) $\lambda_{em}$ 600 ± 50 nm. Scale bar= 10µm. Experiments were repeated 3 times independently with similar results

##### 9.2.5 Analysis of DNA reporter signal loss after PNA tagging

CHO cells stably expressing doxycycline inducible Cys-P1-EGFR-eYFP (25,000) were seeded and incubated in 200 µL Hams F12 media (10% FBS) overnight at 37°C. Cells were induced with 500 ng/mL doxycycline by switching media (1% FBS, 500 ng/mL doxycyline) 16 hours prior to experiments. 4.5 hours prior to the experiment, media was switched to 0% FBS Hams F12 media without doxycycline. After washing with DPBS, cells were treated with **$PNA_{15}$-P2** (100 nM) for 4 minutes in DPBS. Subsequently, cells were washed with HBSS-BB before 4 min incubation with Atto565-labeled DNA-15mer in HBSS-BB. BB: Blocking Buffer: 0.1mg/mL salmon sperm DNA, 0.2% BSA, 1x ProLong™ Live Antifade in HBSS. Cells were kept at 37 °C, with 5 % $CO_2$, and 95% relative humidity using the Cell Vivo incubation system from *Olympus.* Fluorescence images were taken approximately 5-10 minutes after labelling with Atto565 and at hourly timepoints thereafter, with the same cells in frame for each experiment. Regions of interest (ROIs), encompassing the same cells per experiment and excluding areas without cells, were analyzed (Figure 9-10b) and mean intensity of the ROIs were compared between the Atto565 and YFP channels, over time. The loss of signal in the Atto565 and YFP channels was was plotted in Figure 9-10.

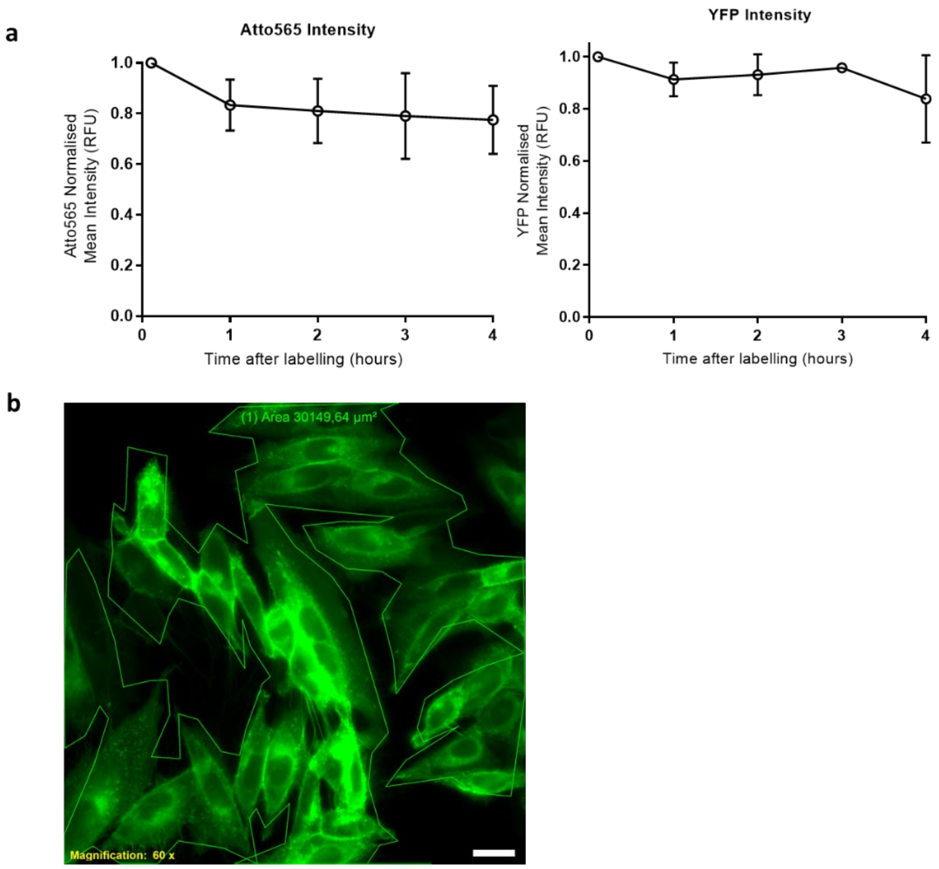

Figure 9-10 **Analysis of DNA reporter signal loss after $PNA_{15}$ tagging of Cys-P1-EGFR-eYFP CHO cells followed by hybridization of Atto565-DNA-15mer. a)** Loss of signal in the Atto565 and YFP channels over time at 37°C. Intensity values calculated as described in b). All data points were normalized to the first time point using GraphPad 7.0. Data is presented as the mean +/- SD of n=3 independent experiments. **b)** Representative ROI taken to calculate mean intensity in the YFP and Atto565 channel in a). YFP brightness was increased before drawing ROI around the borders of the cell population of at least 20 cells Scale bar= 10 µm.

#### 9.3 Labelling of Cys-E3-h$Y_2$R on HEK293-cells: comparison of 16mer and 11mer PNA:DNA duplex stability

Stably transfected HEK293-E3-h$Y_2$R cells[7,8] were seeded out into 8-well µ-slides (*ibidi*), which were prior coated with poly-*D*-lysine (0.01% w/v in DPBS w/o $Ca^{2+}$ and $Mg^{2+}$) and grown to 90% confluence overnight. Labelling and microscopic studies were performed on the next day. For nuclear staining of cells the cell culture medium was exchanged with 20 mM HEPES in HBSS buffer at pH 7 and 2 µL of Hoechst33342, followed by incubation for 10 min at 37 °C. Afterwards, the cells were labeled with a 100 nM **$PNA_{11}$-K3** or **$PNA_{16}$-K3** in HBSS buffer supplemented with 0.1 mM TCEP and 20 mM HEPES at pH 7 for 4 min. After washing with 200 mM $NaHCO_3$ in DPBS buffer (w/o $Ca^{2+}$ and $Mg^{2+}$) at pH 8.4 for 1.5 min, complementary DNA (1 µM) containing 6mer overhangs; **FAM-$DNA_{17}$** or **FAM-$DNA_{22}$**, was added for 5 min in 20 mM HEPES. Cells were washed with OptiMEM before microscopic studies were performed in OptiMEM at 37°C.

FAM-DNA sequences with 6mer overhangs underlined:

**FAM-$DNA_{17}$** FAM-GG ACG AAT AC<u>A CAT CTC</u>

**FAM-$DNA_{22}$** <u>G CGG CA</u>C GGA AAC ATA AGC AGG-FAM

Staining was weak when nucleic acid hybridization was performed at 37°C by allowing the formation of the $PNA_{11}$-$DNA_{17}$ duplex (Extended Data Figure 3a). Intense membrane staining was achieved when nucleic acid hybridization involved the formation of the $PNA_{16}$-$DNA_{22}$ duplex (Extended Data Figure 3b). Based on this result, PNA-15mer and -16mers were used for the further labelling experiments.

### 10 Immunofluorescence

Detection of EGFR phosphorylation in a well plate format was carried out by fluorescent labelling using antibodies against phosphorylated EGFR (Y1068) and using YFP signal as an internal standard. Phospho-EGF Receptor (Tyr1068) ((D7A5), XP® Rabbit mAb #3777 Lot: TB264033) was purchased from *Cell Signalling* and used in an 800x dilution. Goat anti-Rabbit IgG (H+L) Highly Cross-Adsorbed Secondary Antibody, Alexa Fluor Plus 647 (catalogue number A32733, Lot:10) was purchased from *Thermo Fischer Scientific* and used as a 500x dilution.

#### 10.1 Cell preparation

Cys-P1-EGFR-eYFP expressing CHO cells were plated 5000 cells/well on a 96-well culture plate (CulturePlate-96, *Perkin Elmer)*. After 18 hours cells were induced by adding 100 µL doxycycline to a final concentration of 0.1 µg/mL. After five hours serum was switched to serum free media with 0.1 µg/mL doxycycline. Media was removed and cells washed with HBSS.

Cell were incubated with PNA donor **$PNA_{15}$-P2** (50 µL, 100 nM in HBSS, pH 7.0) or PBS for 4 min before washing (x2, HBSS). Either 100 nM DNA 15mer or PBS was added, and cells incubated for 3 min before addition of EGF solution to a final concentration of 100 nM. The cells were incubated at 37° C for 10 min before fixation and analysis by immunofluorescence.

#### 10.2 Immunofluorescence assay

After the relevant experiment, cells were fixed by incubation with fixing solution (4% paraformaldehyde in PBS, 100 µL / well) for 10 min at room temperature. Cells were washed (x3 in PBS) and treated with Blocking Buffer (5% normal goat serum, 0.3% Triton in PBS) for one hour at room temperature. After rinsing (x3 in *Tris*-Hcl Buffered Saline -TBS) the Phospho-EGF Receptor (Tyr1068, D7A5), antibody was added as 50 µL of a 1:800 dilution in buffer (1% BSA, 0.3% Triton in PBS) and incubated overnight at 4 °C. After washing (x3 in TBS) cells were incubated for one hour with AF647-labelled Goat anti-Rabbit IgG as 50 µL of a 1:500 dilution in buffer (1% BSA, 0.3% Triton in PBS). After washing (x3 for 5 min in TBS) the plate was read in a Viktor X5 Multimode Plate Reader (*Perkin Elmer*) using the following channels: YFP: $\lambda_{ex}$ = 420±15, $\lambda_{em}$ = 532±10 nm. AF647: $\lambda_{ex}$ = 665±10 nm $\lambda_{em}$ = 640±10 nm. Without PNA tagging treatment with EGF resulted in a2.8-fold increase of Y1068 phosphorylation. The responsiveness of the EGFR remained after PNA-tagging and DNA hybridisation (3.4-fold increase).

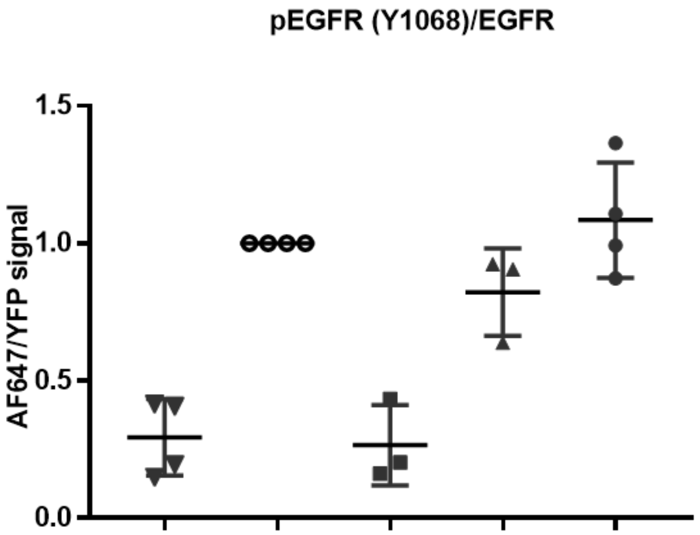

| Condition | 1 | 2 | 3 | 4 | 5 |
|---|---|---|---|---|---|
| $PNA_{15}$-P2 | - | - | + | + | + |
| DNA 15mer | - | - | - | - | + |
| EGF | - | + | - | + | + |

Figure 10-1 **Relative EGFR-Y1068 phosphorylation of Cys-P1-EGFR-eYFP stable CHO cells, after PNA labelling and EGF stimulation, as determined by immunofluorescence 96 well plate assay.** Treatment with 100 nM **$PNA_{15}$-P2**, 100 nM DNA15mer, 100 nM EGF as described in chapter 10.1. Immunofluorescence assay described in chapter 10.2. Readout was measured in ratio of AF647/YFP in relative fluorescence units (RFU). Each replicate was normalized so that condition 2 (+EGF;-PNA;-DNA) was fixed at 1 RFU using GraphPad 7.0. Each data point on the graph is the mean of three wells measured in the same experiment, and for each well 5000 cells were plated. Dot plot is presented as the mean +/- SD of n=3 (conditions 3,4) n=4 (conditions 1,2,5) biological triplicates/quadruplets.

### 11 GPCR characterization by signal transduction assays

Lipofectamine® 2000 was obtained from *Invitrogen* (Carlsbad, CA, USA) and Metafecene® Pro was obtained from *Biontex Laboratories GmbH* (Munich, Germany). IP-One Gq kit was purchased from *Cis-Bio* (Waltham, MA, USA). All cell culture plates were obtained *from Greiner Bio-one* (Kremsmünster, Austria). Endothelin 1 (ET-1) was purchased from *Bachem Holding* (Bubendorf, Switzerland).

#### 11.1 Radioactive inositol triphosphate accumulation assay for characterization of Cys-E3-hY$_2$R

Characterization of Cys-E3-hY$_2$R-eYFP from chapter 9-3 was previously described.[7] It was demonstrated that the introduction of the N-terminal Cys-E3 tag does not impair the biological activity of the hY$_2$R.

#### 11.2 Characterization of Cys-P1-ET$_B$R-GPFspark activation and internalization

##### 11.2.1 Inositol phosphate accumulation assay for characterization of Cys-P1-ET$_B$R-GFPspark

COS-7 cells were grown in T25 cell culture flask until 70 % confluence was reached. For transient transfection, 4000 ng pCMV3-Cys-E3-ET$_B$R-GFPspark or pCMV3-EDNRB-GFPspark and 15 µL Metafecene® Pro in DMEM were used according to the manufacturer's protocol. Cells were incubated with the transfection mix overnight at 37 °C. The next day, cells were washed with PBS and detached by trypsin / EDTA addition before transfer into 384 well plate (10,000 cells per well). Cells were cultured at 37 °C and 95 % humidity for 24 h prior to receptor characterization by IPone Assay (CisBio). Transfected COS-7 cells were incubated with different dilutions of ET-1 for 1 h and inositol monophosphate was detected according to the manufacturer's protocol using Tecan Microplate Reader Spark® plate reader from *Tecan* with the SparkControl™ software, Concentration–response curves and EC$_{50}$ as well as pEC$_{50}$ values were calculated by nonlinear regression using GraphPad Prism 5.0 (Supporting Figure 11A).

##### 11.2.2 Ligand-mediated internalization of Cys-P1-ET$_B$R-GPFspark

HEK293 cells were cultured in DMEM/HAM's F12 (1:1), containing 15 % (v/v) heat-inactivated FCS as monolayers at 37 °C, 5 % $CO_2$ and 95 % humidity. After reaching confluence, HEK293 cells were seeded into 8-well µ-slides, which were prior coated with poly-D-lysine (0.01 % w/v in DPBS). After reaching 70 % confluence, HEK293 cells were transiently transfected with 750 ng of the wildtype pCMV3-ET$_B$R-GFPspark or the pCMV3-Cys-P1-ET$_B$R-GFPspark construct and Lipofectamine® 2000 according to the manufacturer's protocol. Microscopic imaging was performed the next day. The supernatant was exchanged with OptiMEM and cell nuclei were stained with 1 µL Hoechst33342 for 20 min. The cell monolayer was washed once with HBSS and the cells were incubated with 500 nM ET-1 in OptiMEM for 1 h at 37 °C. Fluorescence images were acquired in HBSS after ET-1 removal by washing with HBSS. Used filter sets: Hoechst33342 (filter 02; excitation: G 365, beamsplitter: FT 395; emission: LP 420), and GFP (filter 38; excitation: BP 470/40, beamsplitter: FT 495; emission: BP 525/50).

Introducing the Cys-P1-tag behind the endogenous signal peptide of the $ET_BR$ does not alter the G protein-mediated signalling induced by the peptide ET-1 (Figure S 11A). For both, the wildtype GPCR $ET_BR$-GFPspark and the Cys-P1-$ET_BR$-GFPspark construct a low nanomolar $EC_{50}$ has been detected by signal transduction assays. For the wildtype an $EC_{50}$ of 1.5 nM ($pEC_{50}$ = 8.83 ± 0.05; $E_{max}$ = 100 ± 2 %) was determined, whereas an $EC_{50}$ of 1.4 nM ($pEC_{50}$ = 8.85 ± 0.07; $E_{max}$ = 98 ± 3 %) was found for the tagged $ET_BR$. Additionally, membrane localization studies (Fig. S 11B) revealed the same distribution pattern of both proteins, when expressed in HEK293 cells. Stimulation of these cells with ET-1 lead to complete receptor internalization, indicating the full functionality of the tagged GPCR compared to the wildtype protein.

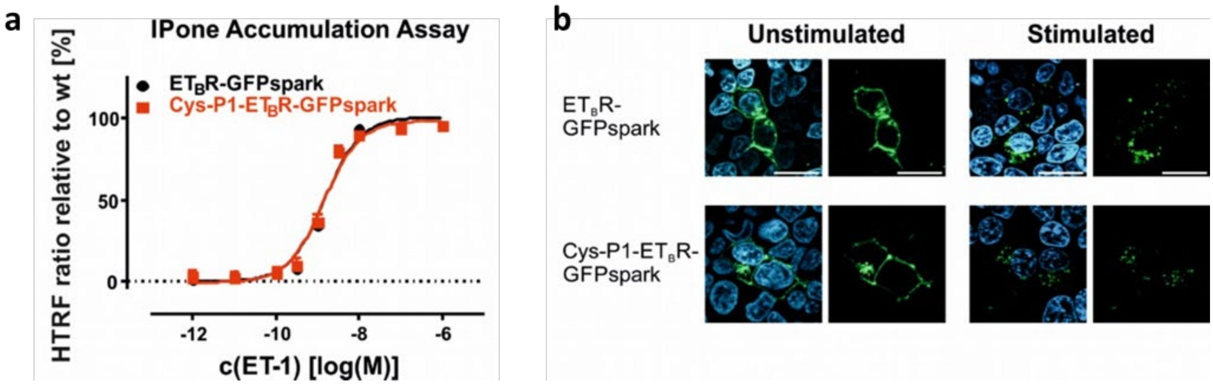

Figure 11-1 **Characterization of Cys-P1-$ET_BR$-GFPspark in comparison to the wildtype $ET_BR$-GFPspark. a)** G-protein activation was tested by IPone accumulation assay in transiently transfected COS-7. Each concentration was measure n=3 times over 5 independent experiments. Data were normalized to the wildtype $ET_BR$-GFPspark protein. Error bars show mean +/- standard error of the mean (SEM). **b)** Fluorescence microscopy images of live HEK293 cells transiently expressing $ET_BR$-GFPspark or Cys-P1-$ET_BR$-GFPspark before and after stimulation with ET-1. Used filter sets: Hoechst33342 (filter 02; excitation: G 365, beamsplitter: FT 395; emission: LP 420), and GFP (filter 38; excitation: BP 470/40, beam splitter: FT 495; emission: BP 525/50). Scale bar: 20 µm. Experiments were repeated 3 times independently with similar results.

### 12 Flow Cytometry Analysis of Multi-fluorophore labelling

Flow cytometry experiments were performed by using an BD Accuri™ C6 (BD Biosciences) with a flow rate of 66 µL/ min for 150 or 260 µL. Data was analyzed using BD CFlow Plus V1.0.227.4 (BD Biosciences). Excitation lasers: YFP) 488 nm; Atto647N) 640 nm. Emission filters: YFP) 533/30 nm; Atto647N) 675/25 nm.

#### 12.1 Multilabelling of Cys-P1-EGFR-eYFP on CHO cells with Atto647N-DNA constructs

Flow cytometry analysis was performed on PNA tagged cells hybridised with **Complex IV**: adaptor DNA-105mer + Atto647N-15mer, **Complex V:** adaptor DNA-105mer 3(Atto647N-15mer) and **Complex VI:** adaptor DNA-105mer +5(Atto647N-15mer) as described in the methods section of the main text.

After gating (Figure 12-1), mean intensities were obtained for YFP and Atto647N. An average of 444 (experiment 1) 6600 (experiment 2) or 9090 (experiment 3) cells were analyzed after gating was applied. Signal was recorded after application of gate 4. Background was defined as the mean intensity obtained from the (-DOX) control after applying gate 2 (Fig 12-1a,1b). For (-Dox) control, cells were treated as described, but without doxycycline induction, addition of **PNA$_{15}$-P2** or addition of DNA complex. The background for the eYFP signal and Atto647N signal were subtracted from the mean intensities. To correct for eYFP expression levels, a ratio of the mean Atto647N intensities (Fig 12-1D) to the mean eYFP intensity was calculated after the gating strategy was applied.

To verify that the DNA complexes do not unspecifically bind to cells an additional control experiment, (-PNA) was carried out. Cells were treated as described, but no **PNA$_{15}$-P2** was added to during the PNA tagging step, and **Complex VI** (5x Atto647N) was added during the hybridization step.

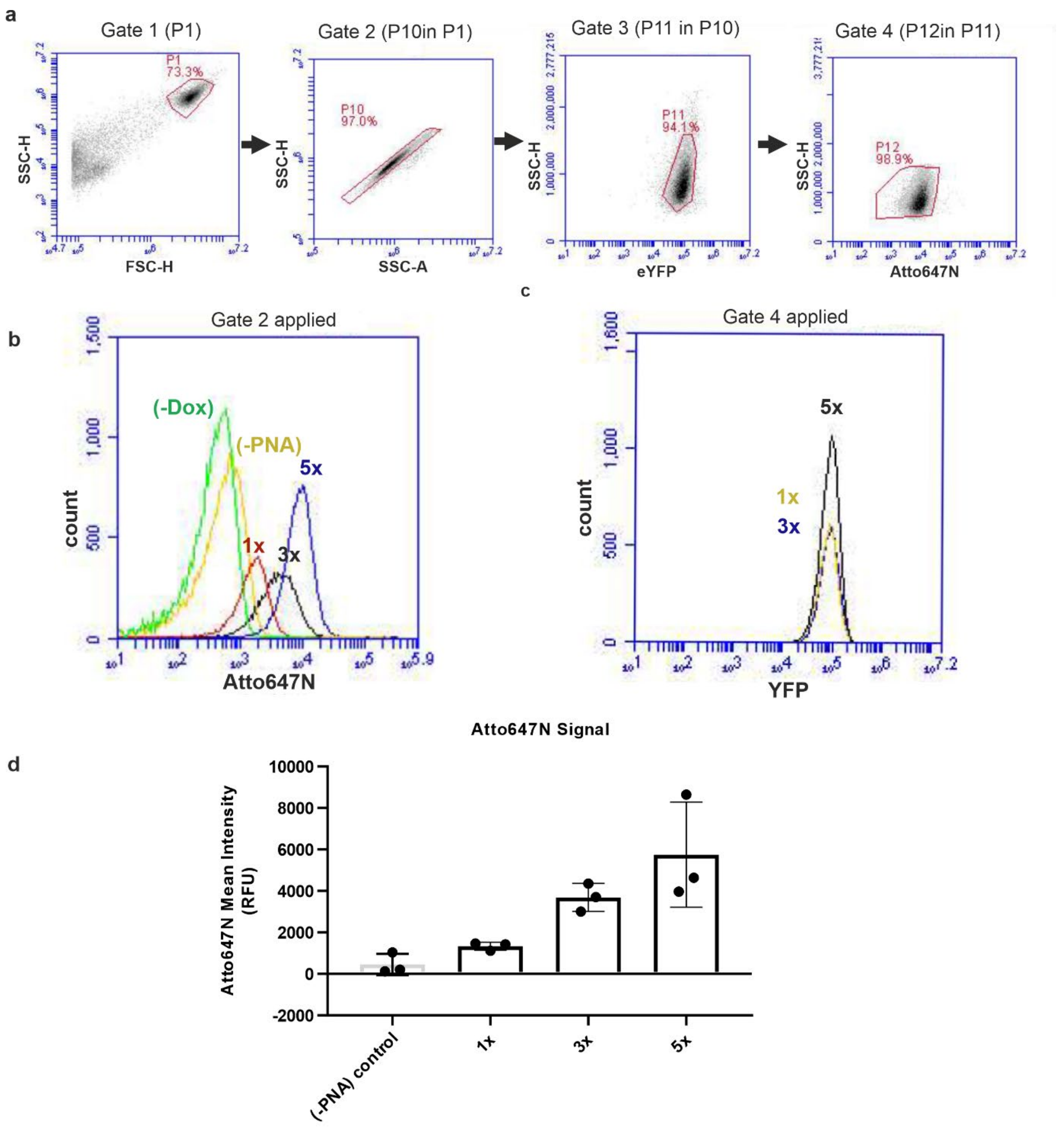

Figure 12-1 **Flow cytometry analysis of PNA labelled Cys-Pa-EGFR-eYFP CHO cells labelled with multiple Atto647N dyes. a)** Exemplified gating strategy for the analysis of CHO cells expressing Cys-P1-EGFR-eYFP upon induction with doxycycline and after tagging with PNA and hybridization with Atto647N-labelled nucleic acid complexes. Debris (FSC-A, SSC-A; Gate 1) and doublets (SSC-H, SSC-W; Gate 2) were excluded and the main populations (eYFP; Gate 3) and (Atto647N; Gate 4) were identified. Histograms were analyzed by fluorescence flow cytometry after PNA labelling and hybridization with 1, 3 or 5 Atto647N DNAs-15mers (complexIV, complexV and complexVI, respectively). (-Dox) control was included to identify background signal and (-PNA) was included as a negative control.**b)** Atto647N channel after applying gate 2 and **c)** YFP channel after applying gate 4. Fluorescence intensity of eYFP and Atto647N is indicated in arbitrary units (A.U.). **d)** Mean intensity of PNA-tagged cells treated with 1, 3 or 5 Atto647N reporter strands, and a (-PNA) control where non PNA- tagged cells were treated 5x reporter strands. For analysis, gate 4 was applied, or gate 2 for (-DOX) control. Data is presented as the mean +/- SD of n=3 three independent experiments.

### 13 References

1 Roloff, A. *Templatgesteuerte Reaktionen von Peptidnukleinsäuren* PhD thesis, Humboldt-Universität zu Berlin, (2014).

2 Mourtas, S., Gatos, D., Kalaitzi, V., Katakalou, C. & Barlos, K. S-4-Methoxytrityl mercapto acids: synthesis and application. *Tetrahedron Letters* **42**, 6965-6967, doi:https://doi.org/10.1016/S0040-4039(01)01423-X (2001).

3 Zitterbart, R. *Chemische Synthese & funktionelle Analyse von immobilisierten Protein-Domänen* PhD thesis, Humboldt-Universität zu Berlin, (2017).

4 Reinhardt, U. *et al.* Peptide-templated acyl transfer: A chemical method for the labeling of membrane proteins on live cells. *Angewandte Chemie - International Edition* **53**, 10237-10241, doi:10.1002/anie.201403214 (2014).

5 Stief, S. M. *et al.* Loss of KDM6A confers drug resistance in acute myeloid leukemia. *Leukemia*, doi:10.1038/s41375-019-0497-6 (2019).

6 Unger, T., Jacobovitch, Y., Dantes, A., Bernheim, R. & Peleg, Y. Applications of the Restriction Free (RF) cloning procedure for molecular manipulations and protein expression. *Journal of Structural Biology* **172**, 34-44, doi:https://doi.org/10.1016/j.jsb.2010.06.016 (2010).

7 Reinhardt, U., Lotze, J., Mörl, K., Beck-Sickinger, A. G. & Seitz, O. Rapid Covalent Fluorescence Labeling of Membrane Proteins on Live Cells via Coiled-Coil Templated Acyl Transfer. *Bioconjugate Chemistry* **26**, 2106-2117, doi:10.1021/acs.bioconjchem.5b00387 (2015).

8 Lotze, J. *et al.* Time-Resolved Tracking of Separately Internalized Neuropeptide Y2 Receptors by Two-Color Pulse-Chase. *ACS Chemical Biology* **13**, 618-627, doi:10.1021/acschembio.7b00999 (2018).
